# Lipid network and moiety analysis for revealing enzymatic dysregulation and mechanistic alterations from lipidomics data

**DOI:** 10.1101/2022.02.04.479101

**Authors:** Tim D. Rose, Nikolai Köhler, Lisa Falk, Lucie Klischat, Olga E. Lazareva, Josch K. Pauling

## Abstract

Lipidomics is of growing importance for clinical and biomedical research due to many associations between lipid metabolism and diseases. The discovery of these associations is facilitated by improved lipid identification and quantification. Sophisticated computational methods are advantageous for interpreting such large-scale data for understanding metabolic processes and their underlying (patho)mechanisms. To generate hypothesis about these mechanisms, the combination of metabolic networks and graph algorithms is a powerful option to pinpoint molecular disease drivers and their interactions. Here we present LINEX^2^ (Lipid Network Explorer), a lipid network analysis framework that fuels biological interpretation of alterations in lipid compositions. By integrating lipid-metabolic reactions from public databases we generate dataset-specific lipid interaction networks. To aid interpretation of these networks we present an enrichment graph algorithm that infers changes in enzymatic activity in the context of their multispecificity from lipidomics data. Our inference method successfully recovered the MBOAT7 enzyme from knock-out data. Furthermore, we mechanistically interpret lipidomic alterations of adipocytes in obesity by leveraging network enrichment and lipid moieties. We address the general lack of lipidomics data mining options to elucidate potential disease mechanisms and make lipidomics more clinically relevant.

**Graphical Abstract:** 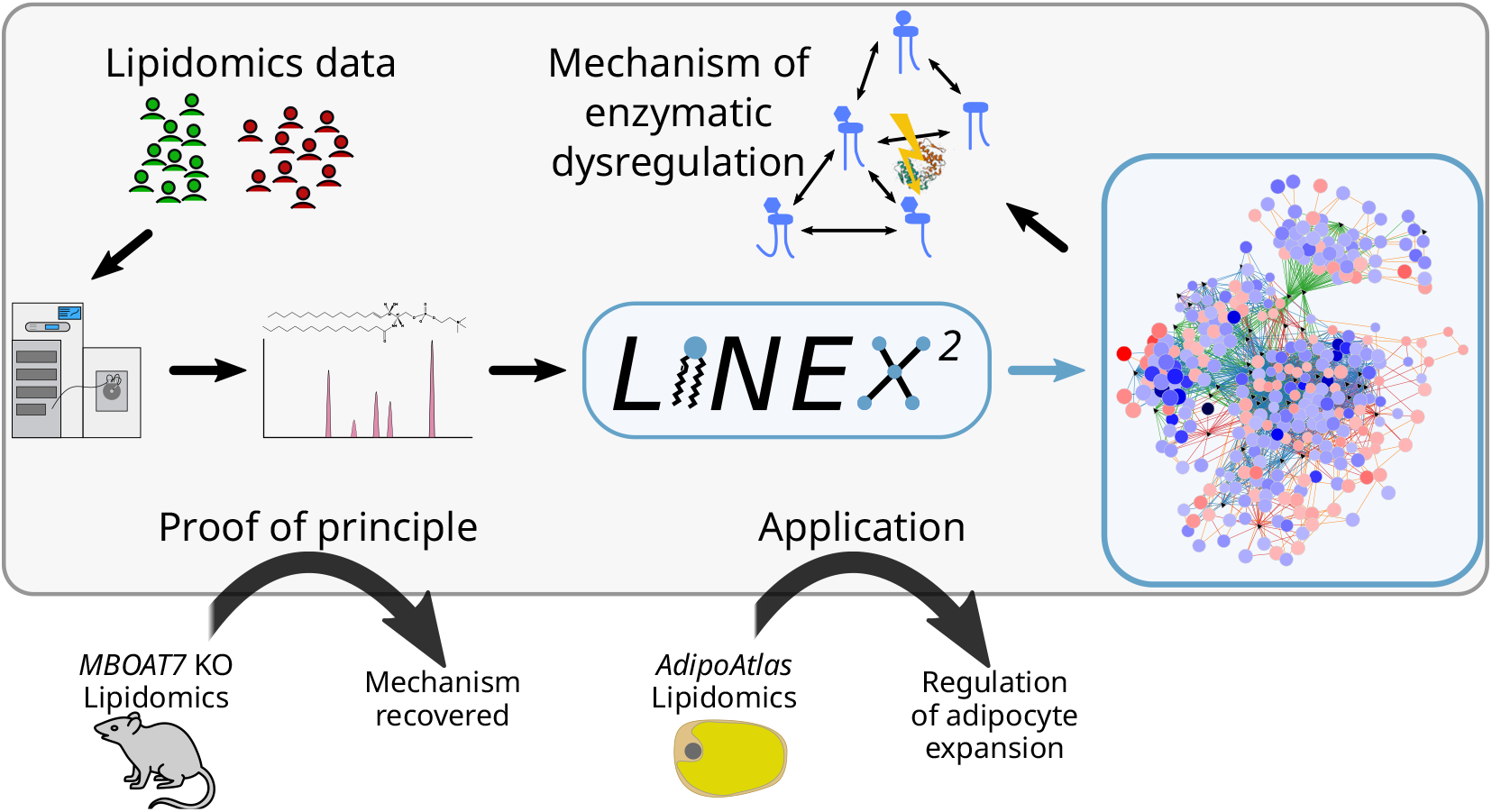

LINEX^2^ (Lipid Network Explorer) is a framework to visualize and analyze quantitative lipidomics data. The included algorithms offer new perspectives on the lipidome and can propose potential mechanisms of dysregulation.

- Using the Reactome and Rhea databases, a comprehensive set of lipid class reactions is included and utilized to map the lipidome on custom data-specific networks.
- With a novel network enrichment method, enzymatic dysregulation can be recovered from lipidomics data.
- We validate its usability on data with a central lipid enzymatic deficiency.
- LINEX^2^ is the first tool capable of such analysis and includes complimentary analysis options for structural lipid analysis. It is freely available as a web service (https://exbio.wzw.tum.de/linex2).

## 1 Introduction

Lipids play a fundamental role in cells across all domains of life. They are not only crucial for the long-term storage of energy but can also influence the activity and occurrence of membrane proteins [1], as well as signalling and inflammatory processes [2, 3]. Therefore, diseases are also influenced by lipids. This is known not only for liver and metabolic diseases [4, 5] but also e.g. various cancers [6, 7, 8, 9]. Despite their essential role in many biological processes, excessive accumulation of lipids, especially in non-adipose tissues can lead to lipotoxicity [10, 11]. Hence, to fully understand diseases on the molecular level, changes in the lipidome have to be characterized and their regulation understood.

Nowadays, an increasing part of the lipidome can be identified and quantified using mass spectrometry (MS). The field, also known as lipidomics, is becoming more relevant for clinical applications and biomarker research [12]. While MS-based lipidomics is not yet used for diagnoses, potential biomarkers have been discussed [13, 14, 15] and disease stratifications based on lipidomics proposed [16, 17]. To gain more insights into disease mechanisms, it is necessary to go beyond classification and prediction by proposing functional interpretations of lipid changes and links to other omics layers. Due to the complexity of both acquired lipidomics data as well as the regulatory mechanisms behind lipid metabolism, dedicated computational tools are of great importance for unraveling these associations.

Such interactions can be studied through biological networks. On the metabolic level, these networks describe reactions between metabolites that are catalyzed by enzymes. When considering lipid metabolic networks an additional constraint is the inherent complexity of the lipidome and its chemical reactions. Lipid enzymes commonly catalyze more than one reaction, this is referred to as multispecificity [18]. This usually means that one enzyme catalyzes a reaction for a group of lipids that e.g. belong to one lipid class but differ in their fatty acyl composition. The combinatorial complexity makes generating lipidome scale metabolic networks for an organism inefficient but instead requires data-specific networks [19, 20, 21].

Metabolic networks are commonly studied with dynamic modeling or constraint based modeling. These techniques allow predictions of the system dynamics, for example the distribution of energy resources. Parameterization of such models requires large amounts of data covering the entire molecular state [22]. Especially metabolic fluxes and well-characterized enzyme kinetics are important, which are often not available in a clinical setting.

Another way to analyze biological networks is through network enrichment. By comparing two experimental conditions, the goal is to find highly connected molecular subnetworks that are enriched with significant genes, proteins, or metabolites. The rationale behind this approach is to propose a mechanistic hypothesis for observed dysregulations. Many algorithms have been developed over the years [23, 24, 25, 26, 27], mainly with a focus on protein-protein interaction (PPI) or gene-regulatory networks. A dedicated method for metabolomics data is included in the MetExplore analysis and visualization software [28]. Their MetaboRank [29] algorithm is an enrichment and network-based fingerprint recommendation method. For lipid networks, an algorithm implemented in the BioPAN software is available, which is capable of creating lipid networks and running de-novo pathway enrichment on them [20, 21]. However, it does not consider reactions involving (de-)esterification. LINEX (Lipid Network Explorer) is a network-based method, which we previously developed [19], addressing this. It combines lipid class and fatty acid metabolism to provide comprehensive networks for computational analysis and lipidomics data interpretation. Using the LINEX framework we previously showed in several studies [19] that new insights into lipidome-wide data can be generated using lipid networks and that central alterations are often metabolically highly related. A limitation of this method is that lipid class reactions have to be entered by users. This requires detailed knowledge about lipid metabolism if reactions beyond the default are required. In contrast to de-novo enrichment on large-scale biological networks, pathway enrichment identifies significantly altered categorized pathways. For metabolites, this can be performed with the KEGG [30] or Reactome database [31]. A recent lipid-specific method is the Lipid Ontology web service (LION/web), which performs an ontology-based enrichment incorporating biological and chemical properties of lipids [32]. So far, no method is available, that puts the multispecifity of lipid enzymes into the center of interpreting lipidomic changes.

Here we present LINEX^2^, a redesigned and extended framework, which addresses the shortcomings of lipid-network based methods. Lipid reactions are based on database information. This provides links to other omics disciplines. Furthermore, we developed a lipid-network enrichment algorithm, that incorporates multispecific enzyme links. The method enables the generation of mechanistic hypothesis from lipidomics data. We successfully applied our method to lipidomics data of a knockout study and reveal potential dysregulations of the lipid metabolism in the adipose tissue of obese humans. This can help to better translate lipidomics into clinical application [33, 34] and improve our understanding of the role of lipid metabolism in disease mechanisms.

## 2 Results

### 2.1 A framework for lipid network analysis

The workflow of a lipidomics experiment can be divided into five steps: sampling, sample preparation, data acquisition, data processing, and data interpretation [35]. LINEX^2^ is aiming at the biological interpretation of lipidomics data (Figure 1). The LINEX^2^ builds data-specific lipid metabolic networks. To obtain these networks, we developed a network extension algorithm (Figure 1, purple box), where metabolic reactions on the lipid class level and fatty acid reactions are extended to the lipid species level. Network extension is possible with molecular species (e.g. DG(16:0_18:1)) or sum species data (e.g. DG(34:1)). Sum species are internally converted to molecular species, to incorporate modifications or additions/removals of fatty acids. This is achieved by finding sets of fatty acyls matching the sum composition using common fatty acids defined per lipid class (e.g. DG(16:0_18:1), DG(16:1_18:0), or DG(14:0_20:1) for DG(34:1)). If molecular species are identified but not quantified they can be used instead of inferring fatty acyl sets, as reported in some studies [16]. An example for the network extension is the lipid class reaction between a Phosphatidylcholine (PC) and a Diacylglycerol (DG) (PC → DG), where the phosphocholine headgroup is cleaved off, is applied to the molecular lipid species PC(16:0_18:1) → DG(16:0_18:1) (for a detailed description see Materials & Methods section Network extension). Also, fatty acid reactions, such as elongation or desaturation can optionally be added to the network as heuristics, e.g. for Lyso-PC(18:0) (LPC(18:0)) → LPC(18:1). Since such reactions usually do not occur on complex lipids directly, but rather as activated fatty acids, they help to visualize fatty acid-specific effects on the network, as previously shown [19], and facilitate computational network analysis.

**Figure 1:**
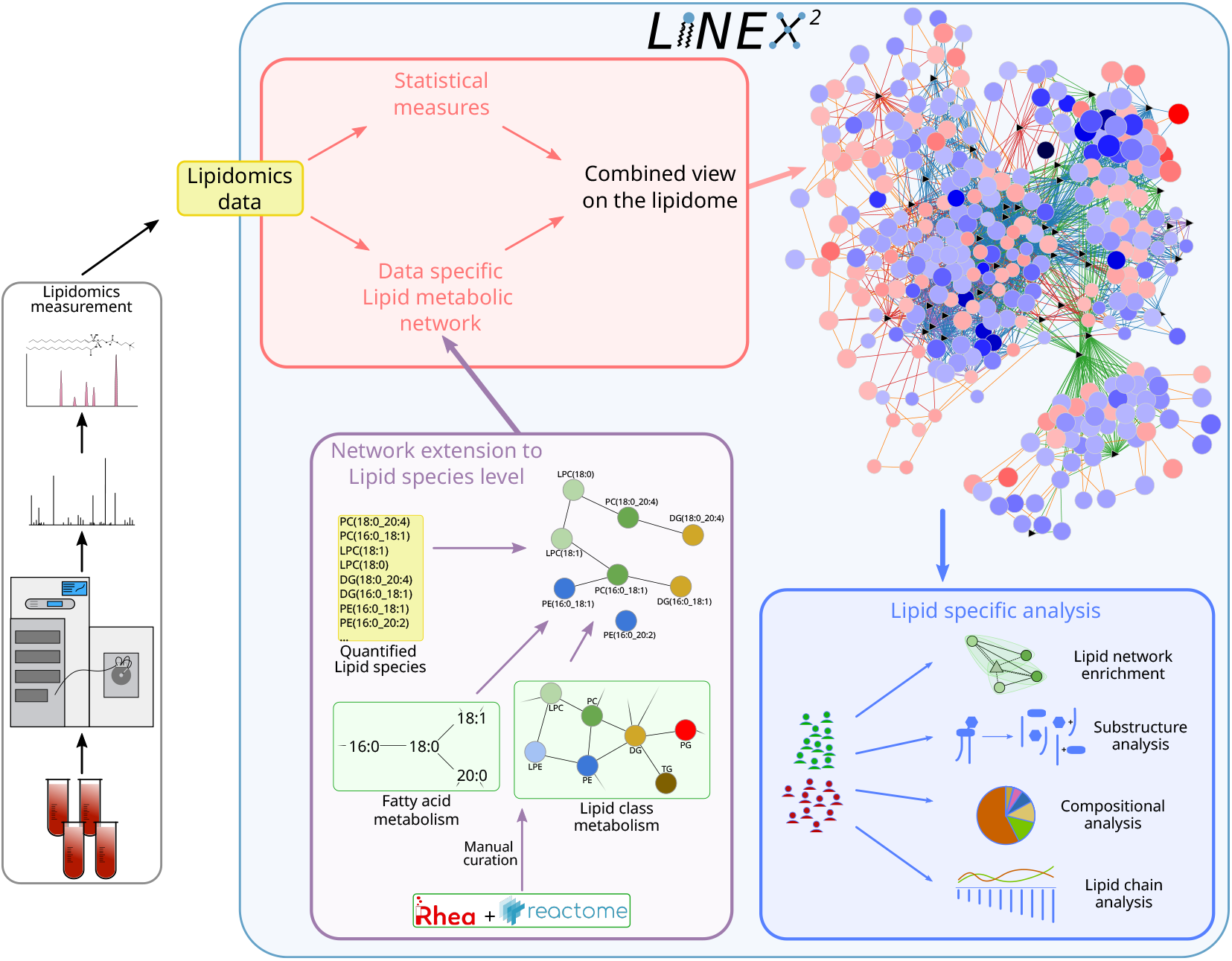
Lipidomics data is used as an input to LINEX^2^. The lipids are then utilized to perform network extension that converts lipid class and fatty acid metabolic networks to lipid species, which are then visualized together with statistical measures such as t-tests or correlations. The network is also used as a basis for lipid substructure, compositional, and lipid chain analysis. A lipid network enrichment algorithm, that takes enzymatic multi specificity into account, can be used to generate hypotheses for enzymatic dysregulation.

### 2.2 Comprehensive curation of lipid-metabolic reactions

The basis for our network extension are publicly available metabolic reaction databases. To provide a comprehensive overview of lipid metabolism, we curated lipid class reactions from the Rhea [36] and Reactome [31] databases (Figure 2A). As a reference for lipid classes, we updated the lipid classes from the ALEX123 lipid database [37] (see Data Availability section). During curation, we removed all transport reactions and specialized modifications such as oxidations or fatty acid branching, which cannot be annotated to standardized lipid classes or are not generalizable for automated network extension. Curation resulted in over 3000 annotated reactions from both databases combined (Figure 2A) across organisms, including organism-specific reactions from Reactome. The top three organisms including the most reactions from Reactome are Homo sapiens (HSA), Rattus norvegicus (RNO), and Mus musculus (MMU) (Figure 2B).

**Figure 2:**
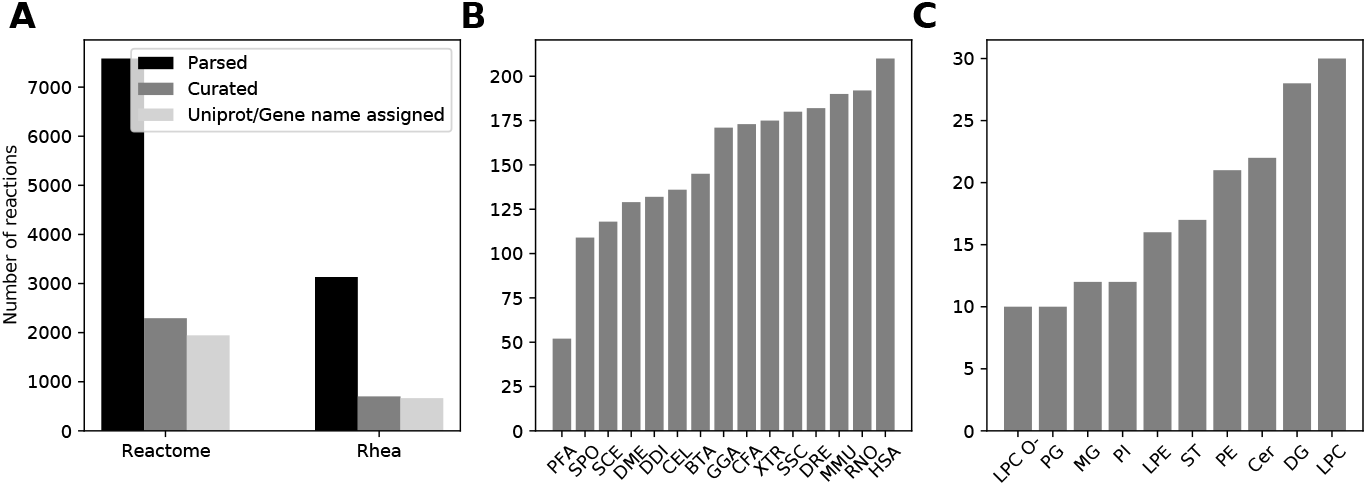
**A** Number of lipid-reactions parsed from Reactome and Rhea databases (black), after curation for available lipid classes and number of curated reactions (dark-grey), which Uniprot or gene name annotations were available (light-grey). **B** Curated reactions per organism from the Reactome database (Rhea does not list details about organisms). **C** Top ten lipid classes with the most curated class reactions.

In cellular lipid metabolism multiple enzymes may catalyze the same lipid class reactions but exhibit different substrate affinities based on the molecular fatty acyl composition. We made all annotated enzymes per class reaction available. After database processing, LPC is the lipid class participating in most reactions (Figure 2C), followed by DG. All reaction identifiers are individually linked, providing a reference to the original database entries in the network.

To keep the freely available LINEX^2^ software up-to-date, user contributions for new lipid classes and lipid-metabolic reactions can be made using an online form (https://exbio.wzw.tum.de/linex2). This way LINEX^2^ can be updated in a community effort to enhance support for less studied parts of the lipidome.

### 2.3 An approach to analyzing lipid networks

For interpreting quantitative changes in molecular networks, network enrichment can be a powerful approach. In the context of metabolic or lipid networks, such methods can reveal underlying changes in enzymatic activity. However, it is more challenging than network enrichment of e.g. proteomics data on PPI networks where changes in protein amounts correspond directly to functional changes of the nodes in the network. In PPI networks changes in protein abundances correspond directly to functional changes of the nodes, representing proteins, in the network. However, when analyzing (lipid-)metabolic networks enzymatic changes can only be approximated from changes in metabolite abundances between experimental conditions. In lipid-metabolic networks, an additional challenge comes from the multispecificity of involved enzymes. In LINEX^2^-networks (as implemented in the network extension) every edge between two lipid species corresponds to an enzymatic reaction, therefore enzymes can correspond to multiple edges.

Our method is designed to explicitly take multispecificity into account. Therefore, a hypernetwork, establishing connections not only between lipids but also reactions, is required. In the hypernetwork more than two nodes can be connected with one (hyper)edge. Based on this representation, the enrichment algorithm can easily connect solutions from the same class reaction, promoting solutions explainable by a few metabolic reactions. Figure 3A shows the workflow of the enrichment analysis (for details see the Materials & Methods section Network enrichment). We start with a LINEX^2^-network, where reactions are represented as edges (1). In the next step, we add lipid class reactions as a second type of nodes to the network (2). Edges between a class reaction node and all lipid species participating in this reaction are introduced, in addition to lipid-lipid edges, that represent conversions. This network is converted to a hypernetwork, where each hyperedge represents a lipid species reaction with lipid-substrates, -products, and reaction nodes (3). For each hyperedge (lipid species reaction), the dysregulation is quantified by the relative change of the lipid substrate-product ratio or difference between two experimental conditions (4). Considering both substrates and products is especially important for reversible reactions [39]. The reaction network is then used to find a maximally dysregulated subnetwork by employing a simulated annealing-supported local search (5). Heuristic reactions are penalized in the objective function of the network enrichment and serve only to increase connectivity. Additionally, the number of class reactions in the network can be penalized to favor parsimonious solutions with a simple mechanistic explanation.

**Figure 3:**
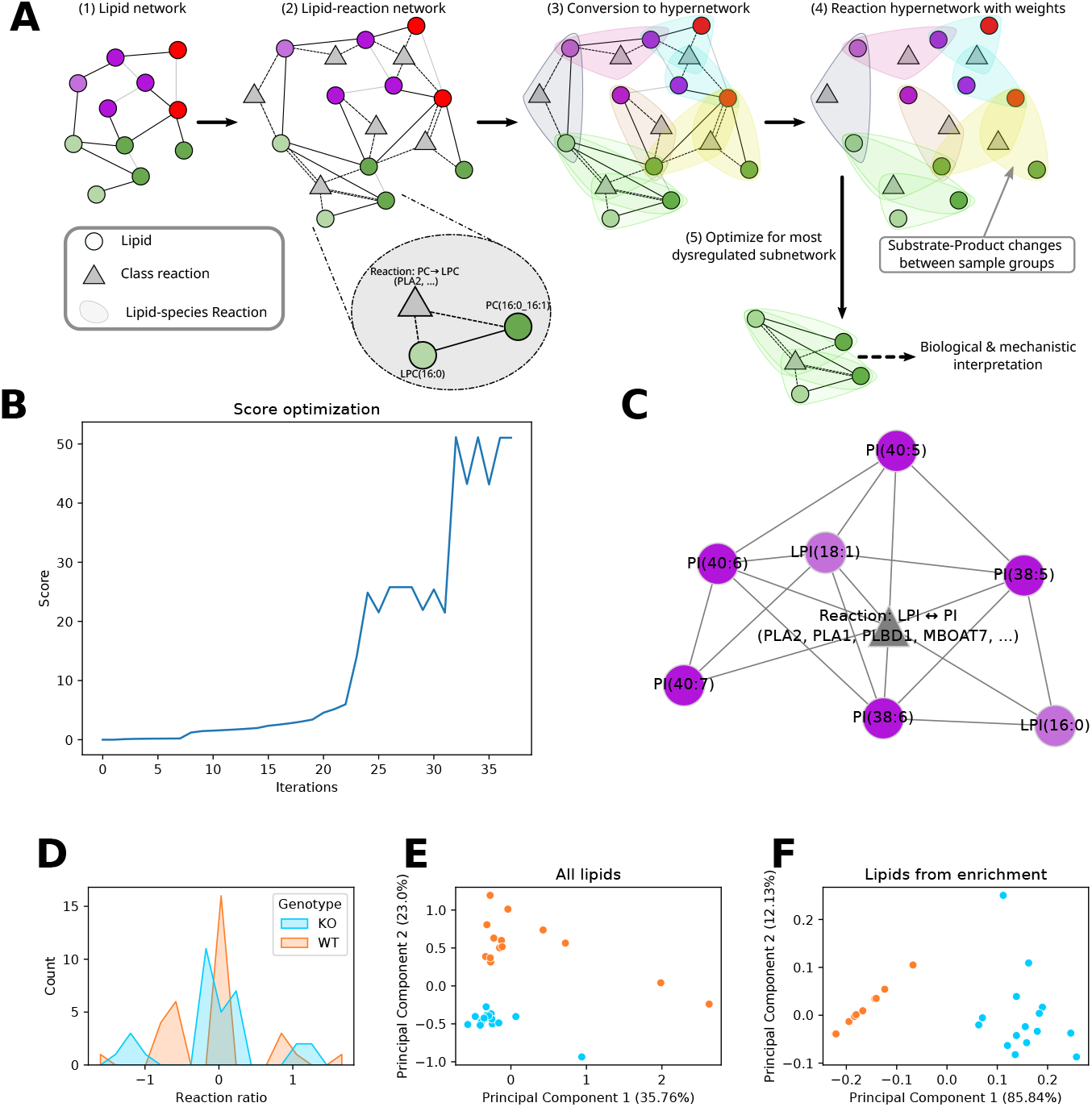
**A** Description of network enrichment workflow. In brief, the lipid network is converted into a hypernetwork, in which hyperedges correspond to lipid species reactions. Based on the computed dysregulation per hyperedge, an optimization algorithm finds the subnetwork with the maximum dysregulation **B** Optimal subnetwork predicted by the enrichment algorithm for mice liver lipidomics data by Thangapandi et al. [38]. The comparison is between wild-type and MBOAT7 knock-out samples. The resulting subnetwork shows the LPI ↔ PI reaction at the center, surrounded by polyunsaturated PI species and two LPI species. **C** Progression of the objective function score during optimization that yielded the subnetwork in B. **D** Substrate-product ratio distribution for the LPI ↔ PI class reaction for all lipid species reactions per genotype (MBOAT7 deficient (KO) and wild type (WT)). **E** Principal component analysis of full lipidomics data and **F** of a subset of the lipidomics data containing only the lipids from the enriched subnetwork from B. The color code is the same as in D for both plots.

### 2.4 Inferring known enzymatic dysregulation from a knock-out study

As a proof of principle for the enrichment, we selected data from Thangapandi et al. [38]. In this study, the authors compared liver lipidomics of mice with a hepatospecific deficiency of MBOAT7 (KO) to wild-type (WT) mice under non-alcoholic fatty liver disease (NAFLD) condition. MBOAT7 catalyzes the class reaction fatty acyl-CoA + LPI → PI + CoA with a specific preference for Arachidonic acid (20:4(ω-6), AA) [40]. The data from Thangapandi et al. [38] is well suited for testing our enrichment algorithm because the enzymatic origin of lipidomic changes in liver tissue is known and the lipidome is affected by the disease.

Figure 3B shows the score progression during the optimization of the algorithm. The temporary plateau at a score of 25 shows the need for global approximation methods such as simulated annealing. In Figure 3C the optimal subnetwork is shown (full network available in the supplement). It consists only of PI, LPI species, and one class reaction. This class reaction represents the transformation between LPI and PI. LINEX^2^ cannot differentiate between the exact enzyme for this reaction. However, in contrast to e.g. PLA2, MBOAT7 only catalyzes LPI → PI class reactions. Additionally, MBOAT7 is known for a higher affinity for AA [40]. This preference can also be observed in the solution in Figure 3C for the edge between LPI(18:1) and PI(38:5), under the assumption that this reaction can only occur if the molecular composition of PI(38:5) is PI(18:1_20:4). Furthermore, all other reactions between LPIs and PIs are only possible for the addition/removal of fatty acyls with at least 20 carbon atoms and 4 double bonds. These results are not surprising, because of the structural similarity of AA to other (very)-long-chain polyunsaturated fatty acids (Supplementary Figure S1). While LINEX^2^ is not able to directly pinpoint MBOAT7, the results demonstrate its capability to find strong hypotheses for enzymatic dysregulation from lipidomics data.

To evaluate the enrichment results, we implemented an empirical p-value estimation procedure (detailed description in Materials & Methods section Network enrichment). This is computed by comparing the score of the final solution to a distribution of sets of unconnected reactions of the same size, to evaluate whether the final connected subnetwork has a significantly higher score. The MBOAT7 enrichment result (Figure 3C) has a p-value of 0.0018, indicating the likeliness of the mechanistic solution.

When investigating the distributions of the LPI ↔ PI class reaction (i.e. over all respective lipid species reactions) per genotype (Figure 3D), no strong distribution shift in one direction can be observed. The distributions show a peak around zero, indicating that many reaction ratios are not influenced by the MBOAT7 knock-out (KO). However, two more peaks around 1 and −1 can be observed for both conditions, where the peaks of the KO are shifted slightly more towards absolutely higher values. Despite these subtle differences, it is not possible to draw a hypothesis towards a mechanistic explanation including fatty acid-specific effects. In Figure 3E we plotted the principal component analysis (PCA) of the full lipidomics data. In contrast, Figure 3F shows the PCA plot based only on the lipidomics data for the lipid species present in the enrichment subnetwork (Figure 3C). In the PCA of all lipids, PC2 reflects the variance corresponding to the genotype, explaining 23% of the total variance. However, after selecting the LPI and PI species from the enrichment solution, the genotypic difference makes up for the majority of the variance with almost 86%. This means that the lipids in the subnetwork (Figure 3C) represent the effect of the MBOAT7 knock-out almost entirely.

These results demonstrate the ability of the enrichment analysis to develop reasonable hypotheses on enzymatic dysregulation based on lipidomics data. The result not only shows an increased variance corresponding to the genotype but also allows mechanistic lipid species-specific explanations.

### 2.5 A mechanistic hypothesis for adipocyte expansion in obesity

We further aimed at improving our understanding of the changes in lipid-metabolism of lipid-related diseases. For this purpose, we selected the AdipoAtlas [41], a reference lipidome of adipose tissue in lean and obese humans. The authors identified 1636 molecular lipid species, out of which 737 were quantified. Out of all semi-absolutely quantified lipid species, only 26 - solely carnitines - could not be mapped out of the box, showing that a full reference lipidome can be analyzed with the LINEX^2^ database integration.

#### 2.5.1 Network analysis indicates a mechanism for adipocyte expansion

We used our network enrichment algorithm, which resulted in the subnetwork shown in Figure 4A. The subnetwork contains three reactions, which all represent an acyl-transferase reaction between Lyso-Phospholipids. Investigating the reaction ratios of these three class reactions over all possible species reactions shows equal distributions between obese and lean (Figure 4C). However, considering the species reactions present in the subnetwork reveals differences between the groups with respect to the reaction ratios (Figure 4D). These reactions are catalyzed by the Phospholipase A2 Group IVC (PLA2G4C) and the asparaginase (ASPG), which both have lipase and acyl-transferase activity. It has been shown that PLA2 Group IV members preferably act on the sn-2 position and that polyunsaturated fatty-acyls are commonly transferred by them [42]. This preference is reflected in the subnetwork. Literature research shows that PLA2G4C has been reported to be differentially expressed in obese individuals [43, 44] and products of (c)PLA2 activity are known mediators of adipose tissue metabolism [45].

**Figure 4:**
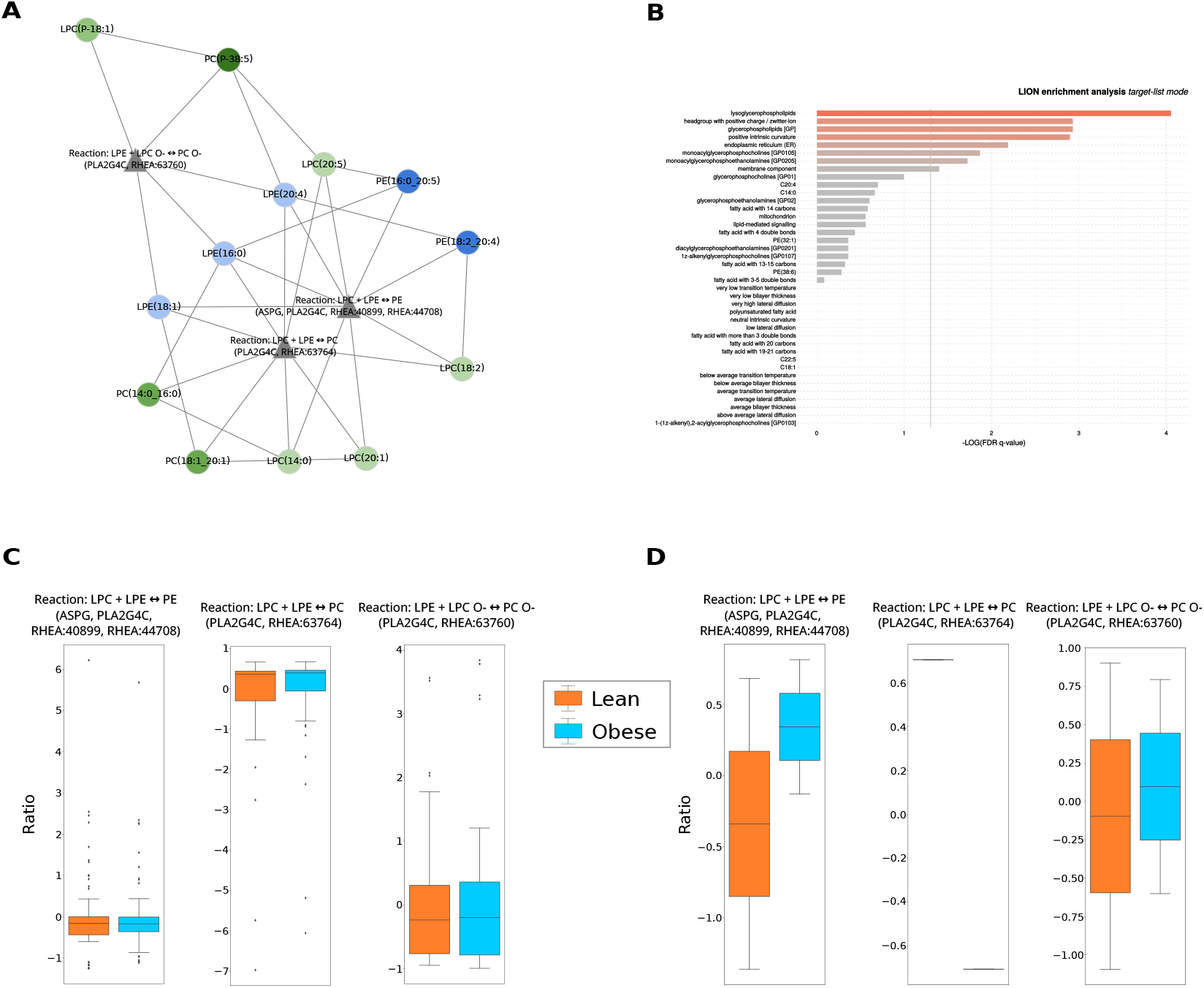
LINEX^2^ application on the AdipoAtlas data. **A** Subnetwork returned by the introduced enrichment algorithm. The enriched subnetwork contains three reaction nodes, all representing fatty acid transfer between lysophospho- and phospholipids. Furthermore, the network shows a preference for long-chain polyunsaturated fatty acids. **B** LION enrichment using the lipids in the subnetwork (A) as targets in the ‘target list mode’. **C** Distribution of the substrate to product changes (see Methods - Substrate-product change calculation) for the three reactions present in A *over all possible lipid species combinations* from the AdipoAtlas data. **D** Distribution of the substrate to product changes using only the *lipid species combinations identified in A*. Both C and D ratios are shown as per-reaction z-scores.

The prevalence of acyl-transferase reactions in the subnetwork suggests a transfer of FAs between lipids with a Phosphocholine and a Phosphoethanolamine headgroup and their respective Lyso-Phospholipid species. The ratio of LPC/LPE to PC/PE as well as the ratio of lipids with a Phos-phocholine headgroup to lipids with a Phosphoethanolamine headgroup influences the membrane curvature [46, 47]. This property is important because adipocytes expand in obesity [48]. A change in this ratio has also been associated with altered membrane integrity and fluidity [49, 50]. We confirmed this with a Lipid Ontology (LION) enrichment analysis [32], where we used the lipids of the enriched subnetworks as a target list (Figure 4B). The analysis resulted in membrane curvature and other membrane-related terms. Additionally, we observed similar behavior in the development of mesenchymal stem cells to adipogenic cells based on data from Levental et al. [51] (Supplementary Figure S2B). These insights further support the practical feasibility of our reaction enrichment approach.

#### 2.5.2 Lipid moieties show alterations in neutral lipid composition

Despite changes in the Glycerophospholipid composition that are an indication for adipocyte expansion, accumulation of neutral storage lipids is a major hallmark for obesity. This is also reflected in the network representation of the AdipoAtlas lipidome (Figure 5). It shows increased TG and DG levels in obese samples, and an overall decrease in Glycerophospholipids. Neutral lipid species containing poly-unsaturated FAs have especially high fold changes (Figure 5, Supplementary Figure S3). Concerning chain length, we observe that TG species with a sum length >30 and <57 are accumulated in obese samples (Supplementary Figure S4A). Since this pathway of the lipid metabolism was not picked up by the network enrichment as the strongest dysregulated part, we wanted to further investigate the compositional changes of neutral lipids. For this, we developed a lipid moiety analysis. It quantifies common substructures of lipids across the lipidome to show trends in changes of the lipidome composition (Supplementary Figure S5). As moieties, we define sum length, number of double bonds, head groups, and their combinations. The results go in hand with the observations on the lipid network. Especially lipid species with a sum length >45 and 2 to 3 double bonds show a sharp increase in obesity, predominantly TG species with a length of 49 and 53. Also Sterol esters show significant changes in disease progression. The observed changes in the TG composition are in accordance with previously published results [52]. This analysis can provide additional insights into the lipid metabolism and complement the network analysis.

**Figure 5:**
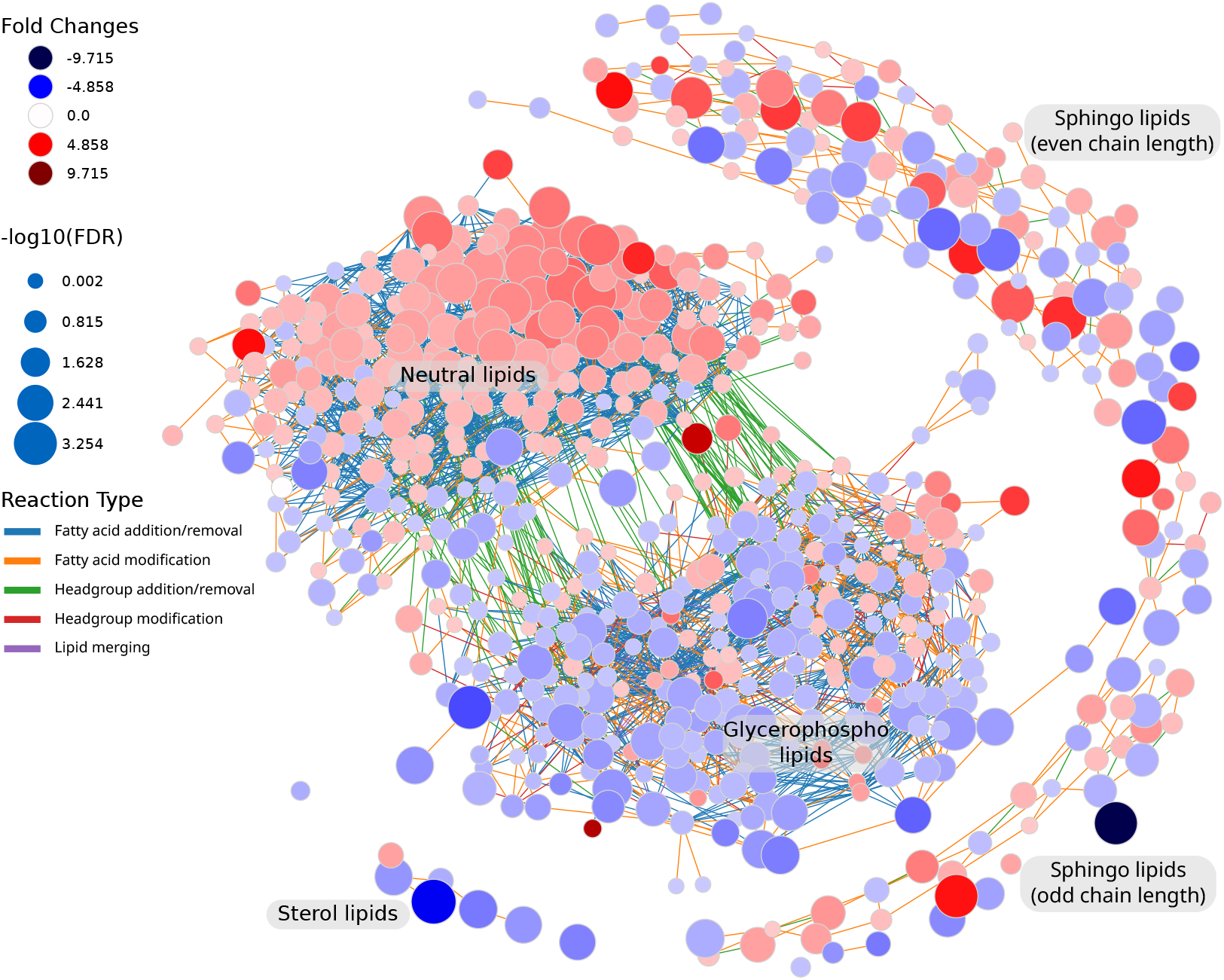
Lipidomics data from the AdipoAtlas visualized with LINEX. In the network lipids are represented as circular nodes. The red color of lipid nodes represents a positive fold change from lean to obese condition, and blue a negative fold change. Edge color indicates the type of reaction connecting two nodes. An interactive version of the network as well as all other analyses conducted with LINEX are available in an HTML file in the supplement.

### 2.6 LINEX^2^ Software

The LINEX^2^ software framework for analysis and visualization of lipid networks is available as a web service at https://exbio.wzw.tum.de/linex2. Lipidomics data and sample annotations can be uploaded as .csv files, to perform not only network enrichment and visualization, but also summarizing statistics, lipid chain analysis [53], and moiety analysis. Results can be viewed and downloaded in an interactive format. For high-throughput analysis, a python package is also available (https://pypi.org/project/linex2/).

## 3 Discussion

We present a method to generate and analyze lipid-metabolic networks. Using curated lipid class reactions from common metabolic databases our method computes data-specific lipid networks. Furthermore, we developed a network enrichment algorithm, to propose hypotheses for enzymatic dysregulation from lipidomics data. As a proof of principle, we applied the approach to liver lipidomics data, where the deficient MBOAT7 enzyme was successfully identified from the data.

Network enrichment for molecular biological data analysis has first been applied in 2002 [54] and a variety of methods have been developed since then. The challenge in generating mechanistic hypothesis from metabolomics or lipidomics data lies in the fact that dysregulation on the enzymatic level is not measured directly. Instead it can only be inferred based on changes in the metabolome, unless full-scale proteomics experiments are run in addition. For lipid networks, only one tool, BioPAN, is available so far [20]. In contrast to our proposed network enrichment algorithm, this method is searching for activated reaction chains between lipids of the same sum composition. The scope of the LINEX^2^ enrichment differs from BioPAN, by searching for dysregulation of multispecific enzymes that likely affect lipids of the same class with different sets of fatty acyls. Another difference is in the network computation. LINEX^2^ includes fatty acyl addition/removal, enabling insights such as the MBOAT7 example we show in this work. To illustrate how LINEX^2^ compares to BioPAN [20], we computed the BioPAN network (Supplementary Figure S6A) as well as the predicted list of active reactions. The results do not include LPI species and only one reaction chain with a PI species (Supplementary Figure S6B). Therefore a hypothesis on MBOAT7 dysregulation cannot be drawn from this method. Nguyen et al. [21] performed a network optimization based on changes in lipid abundances and literature mining of lipid-enzyme interactions. However, they do not infer quantitative values for reactions and no implementation is available.

Hence, LINEX^2^ lipid network enrichment is the only available method that aims at inferring enzymatic dysregulation from lipidomics data. An important aspect of the method is the usage of hypernetworks, to take the multispecificity of lipid enzymes into account, which increases confidence in the retrieved mechanism. Beyond its role in lipid metabolism multispecificity also plays a role in other biological processes, for instance in the glycan metabolism, where enzymes extend various branched glycan structures [55] or DNA methylation [56]. Our principle of network analysis and enrichment could be extended into these fields and help to discover underlying dysregulation.

A limitation of our enrichment algorithm is that it computes substrate-product ratios independent from each other. In reality, however, reactions are linked through shared substrates or products and metabolic changes are propagated through the network. These effects can be due to, e.g. metabolic self-regulation [57] and structural or signaling functions. Since each lipid species takes part in a plethora of reactions, results of altered enzymatic activity might not be observed directly for the substrates and products of that reaction. This is also the case for multiple reactions, which form a consecutive transformation sequence that change at the same time. However, assuming the principle of maximum parsimony, disordered conditions are most likely caused by alterations in only a few enzymatic steps, making the settings for such inaccurate approximations rare cases. Our network extension method depends on generalizable reaction rules. Therefore, manual curation of reaction databases was necessary. Due to a better coverage of commonly measured lipid classes, metabolic databases may be susceptible to research bias. We address this bias by using lipid class reactions instead of enzymes, to prevent well-studied enzymes participating in many reactions from being favourably selected. Additionally, the network enrichment is avoiding bias by correcting for the number of lipid participants in the reaction.

A limitation for the generated hypothesis is the missing knowledge about enzymatic specifity. Therefore, our method is constrained to returning a set of candidate enzymes, which are attributed to the same type of reaction, without pinpointing individual enzymes. With more data available, such as the work from Hayashi et al. [42], better estimates for fatty acid-specific subnetworks can be made.

Lipids exhibit a plethora of structural and signaling functions, beyond energy metabolism. Therefore it is important to not only consider the biosynthesis of lipids, but also the change of biophysical properties. Consequently, a comprehensive computational analysis of the lipidome should include both network-based as well as lipid property-related methods. We also showed this by generating additional insights through the use of LION [32], lipid chain analysis [53], and lipid moiety analysis, which quantifies common lipid substructure features.

With the ability to connect enzymatic activity to lipidomics data, LINEX^2^ provides the basis for a knowledge-driven integration of lipidomics with proteomics data. The inclusion of quantitative proteome information could further improve the performance of the enrichment algorithm presented in this paper and open up the possibility of directly identifying causal proteins. This could be of great value for the causal interpretation of lipidome changes, which would directly translate into relevance for clinical applications, due to the many associations of lipids with various disorders [44, 13, 8, 7, 16].

With our LINEX^2^ web service, we offer new analysis methods for lipidomic data, ranging from network visualization to generating hypotheses for dysregulation. Freely available through a userfriendly interface, lipidomics researchers do not need to be experts in bioinformatics to perform sophisticated analyses of the lipidome in a metabolic context. Moreover, LINEX^2^ networks can be the basis for further methodological developments that help to enhance the biological interpretability of lipidomics experiments by enabling inference of metabolic regulation from lipid data.

## 4 Materials & Methods

### 4.1 Database parsing & curation

We obtained lipid-related reactions from the Rhea [36] and Reactome [31] databases. From Rhea, all reactions involving lipids were parsed (based on ChEBI ontology, a subclass of CHEBI:18059). All reactions included in the category “Metabolism of Lipids” for all available organisms (e.g. R-HSA-556833 for Homo sapiens) were parsed from Reactome.

After parsing, all lipids and reactions were manually curated. Lipids were annotated and assigned to classes according to an updated version of lipid nomenclature from Pauling et al. [37] (Supplementary Table 1). Lipids that could not be annotated were not considered. Lipids are commonly composed of a headgroup, a backbone, and a set of attached fatty acids. From the databases, we extracted reactions showing conversions between common lipid classes, which are usually based on changes in one of these three attributes of lipids. We classified these lipid class reactions with at least one annotated lipid available into different categories: headgroup modification (e.g. PS ↔ PE), headgroup addition/removal (e.g. DG ↔ PA), fatty acid addition/removal (e.g. LPC ↔ PC), lipid merging (e.g. PA + PG ↔ CL) (see next section and Supplementary Figure S7 for more detailed descriptions). Fatty acid reactions on complex lipids are heuristics and can be manually added or banned by the user. Default available reactions are fatty acid elongation (increasing the chain length by 2), fatty acid desaturation (adding one double bond), and hydroxylation/oxidation (adding one hydroxylation/oxidation to a fatty acid).

### 4.2 Network extension to species level

Curated class reactions from databases are used to infer lipid species networks. All steps of this network extension are explained below. To properly evaluate the reactions, molecular lipid species are required. This means that for each lipid the attached fatty acid must be available. Therefore, all lipid species, which are only available as sum species, are converted into a set of possible molecular species. As an example, a PC(40:2) has to be converted into possible molecular species such as PC(20:0_20:2) or PC(22:2_18:0). For this, possible common (class-specific) fatty acids can be added by the user. Only if at least one molecular species can be generated that has the same sum formula as the original sum species, it is considered for the network extension.

Extension of lipid class metabolic networks to lipid species networks can be divided into two steps: extension of the class metabolism and fatty acid metabolism.

#### Extension of class metabolism

Lipid class reactions are evaluated using the defined reaction categories (headgroup removal/addition, headgroup modification, fatty acid addition/removal, and lipid merging) plus ether heuristic. For each reaction, all lipids from the user data, which match the lipid classes that participate in a reaction are selected. Reactions with more than one lipid class as substrate and product are only possible or available for certain reaction categories. If possible, these are explicitly mentioned. The reaction evaluations are under the condition that a lipid class reaction for the substrate-product set exists. For a “Headgroup modification” reaction the substrate and product lipids require the same set of fatty acids, e.g. PS(18:0_16:0) ↔ PE(18:0_16:0, (Supplementary Figure S7A). A “Headgroup addition/removal” also requires the substrate and product lipids to have the same set of fatty acids, e.g. DG(18:0_18:1) ↔ PA(18:0_18:1) (Supplementary Figure S7B). The reaction is also possible for two lipids as substrates and two lipids as products, e.g. PC + Cer ↔ DG + SM (Supplementary Figure S7C). In this case, the headgroup is shifted from one lipid to another. For this evaluation, two substrate product pairs are matched for the lipid donating the headgroup (PC ↔ DG) and the lipid accepting it (Cer ↔ SM). These are then evaluated independently if at least one reaction per pair can be found. The “Fatty acid addition/removal” reactions in the case of one lipid as substrate and product require one lipid with one less fatty acid and the fatty acids of the lipid with fewer fatty acids to be contained in the other lipid, e.g. DG(18:0_18:1) ↔ TG(18:0_18:1_16:0) (Supplementary Figure S7D). For two substrates and two products, a fatty acid is shifted from one lipid to another, e.g. PE + MLCL ↔ CL + LPE. Again, two substrate product pairs are matched for the lipid donating the fatty acid (PC ↔ LPC) and the lipid accepting it (MLCL ↔ CL). They are evaluated independently and the edges are added to the network if two pairs can be found which donate/accept the same fatty acid. Another case exists for reactions with two substrates and one product (e.g. LPC + LPC ↔ PC). Also here, a fatty acid is shifted from one lipid to another, however, the donor is not considered a lipid, after the fatty acid is removed. Similarly, a pair of lipids accepting the fatty acid is formed (LPC ↔ PC). Edges are then added to the network if the accepting and donating lipids have combined the same fatty acids as the resulting lipid. The reaction type “Lipid merging” describes two lipids that are bound together by a reaction, e.g. PG + PA ↔ CL (Supplementary Figure S7E). The molecular species of the substrates require the same combined fatty acids as the resulting lipid for this reaction to occur and be added to the network. We additionally consider fatty acid ether exchange as heuristics. These optional connections are edges between lipid classes and their corresponding ether classes if they share the same set of fatty acids, e.g. LPA(18:1) ↔ LPA(O-18:1), to improve network connectivity and stress fatty acid-specific effects in the network. Edges in the network are undirected since we cannot conclude the net flux of a reaction from the lipidomics data, especially since for most reactions, counterparts in the opposite direction exist.

#### Extension of fatty acid metabolism

Fatty acid synthesis and modification occurs commonly on activated fatty acids and they are not bound to complex lipids. However, to increase network connectivity, fatty acid reactions on complex lipids can be added to the network. As described earlier, this is done through user-defined reactions. A fatty acid reaction, e.g. PC(18:0_16:0) → PC(18:1_16:0), here fatty acid desaturation for the fatty acid 18:0 to 18:1, requires two lipids of the same lipid class and all but one identical fatty acid. Only one fatty acid modification is considered per reaction. In the case of elongation, the non-identical fatty acids require the same amount of double bonds, and other modifications have to differ by the length of two carbon atoms, e.g. 18:0 - 20:0. A desaturation requires fatty acids, which differ by a double bond, with all other attributes being the same. Custom fatty acid metabolism rules can be added, by providing the numeric changes of fatty acid attributes, such as length, double bonds, or modifications. Additionally, reactions between two specific fatty acids can be excluded. For example, the desaturation of fatty acyl 18:2 to 18:3 is not possible in humans.

In the network representation lipids are shown in the provided resolution. In the case of sum species, lipid nodes can also be shown as molecular species (based on the possible molecular species, as explained earlier). Sum species including their statistical properties are then projected onto multiple potential molecular species.

### 4.3 Network enrichment

We developed a novel network enrichment algorithm for lipid networks. It aims to find the most dysregulated lipid subnetwork between two experimental conditions providing a hypothesis for enzymatic alteration/dysregulation. The methodology involves 1. building a reaction network from a standard LINEX network and calculation of substrate-product changes per reaction. 2. Utilization of a local search algorithm to find the heaviest connected subgraph (i.e. the subgraph with the largest average substrate-product change) and 3. an empirical p-value estimation. All steps are described below.

#### Reaction network building

To convert the lipid network to a reaction network, we generate a unique reaction identifier for each reaction (edge) in the network extension. This is especially important for reactions with more than one substrate and product, with multiple edges corresponding to one lipid species reaction. In the next step, all lipid species reactions are converted to a new network representation with reactions as nodes. Edges between two reaction nodes are drawn, if the reaction belongs to the same lipid class reaction or at least one lipid species can be found in both reactions.

#### Substrate-product change calculation

The Substrate-product change is calculated independently for each reaction *r_i_* of the set of all reaction nodes *R* in the network. It describes the relative substrate to product change between two experimental conditions. It can be calculated using absolute or relative substrate-product change. The data for the calculation consists of a set of measured samples *N*. With *C* denoting the subset of samples belonging to the control condition and *D* the samples belonging to the disease condition. The absolute substrate product difference Γ^a^ for reaction *r_i_* for of the disease samples *D* is calculated as

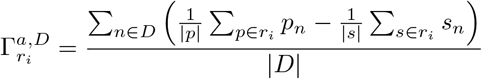

with 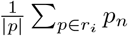 as the mean of all lipid products concentrations from reaction *r_i_* in sample *n* and 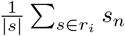 corresponding for substrates. We choose the mean over the sum here to avoid a bias towards reactions with unequal numbers of lipid products or substrates. The relative substrate-product difference Γ^*a*^ is calculated by

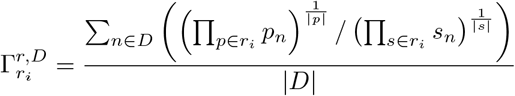

The |*p*|th- and |*s*|th-root are used as bias correction factors in an analogous fashion to using the mean over the sum in the absolute difference calculation. From the user, the absolute or relative score can be used to compute the final reaction score that compares both experimental sets *C* and *D*. It is calculated as follows:

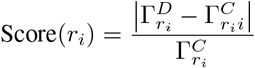

This can be done with the relative or absolute substrate-product change. As previously explained, reactions of the fatty acid metabolism or ether lipid conversions are heuristic, to improve network connectivity. They do not occur directly on the lipid level. For that reason, They are also considered in the network enrichment but penalized (default = −1) to favor the selection of the non-heuristic reactions.

#### Local search and simulated annealing

Local search is a heuristic approach that is usually applied to hard optimization problems [58]. Local search investigates the search space by applying local changes to candidate solutions, such that the objective function value is increasing. The changes are applied until no more local improvements can be made. To avoid stagnation in a local maximum, the simulated annealing procedure [59] allows non-optimal solutions and thus increases the exploration space. The probability of accepting a suboptimal solution depends on the temperature parameter *T*, which decreases over time at rate *α*:

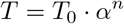

where *T*_0_ is the initial temperature, *α* is the rate of decrease and *n* is the iteration number. If no more local improvements are possible, a random solution is accepted under the following condition:

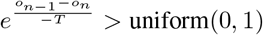

where and are objective function scores at iterations n-1 and n correspondingly.

We employ local search on the reaction network *G* = (*V*, *E*). Starting from a (random) set of connected starting nodes, also called seed, the local search can perform three actions for improvement in the objective function scores: node addition, node deletion, and node substitution. A minimum and maximum size for the subnetwork have to be entered as parameters, preventing the algorithm from selecting too small or big solutions. The action that allows improving the current value of the objective function is accepted, and thus a candidate solution is modified at each iteration. The algorithm terminates when a) no further improvements are possible, b) the simulated annealing condition is not satisfied, or c) the number of maximum iterations is reached. The best-identified subnetwork is returned. The objective function score of a reaction subnetwork *G*\* = (*V*\*,*E*\*) is computed as follows:

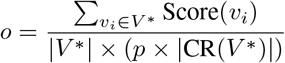

with a user defined penalty *p* for the number of different lipid class reactions in the subnetwork and *CR*(*V* *), the set of different lipid class reactions in the set nodes *V*\*. If the reaction network consists of unconnected components, the local search is run for each component independently and a subgraph for each component is returned.

#### Subnetwork p-value

The network enrichment algorithm results in a subnetwork with a score for each run. To indicate if this subnetwork/score provides a significant insight compared to an equally sized random set of reactions, we compute an empirical p-value. For that, we sample reactions in the range of the minimum and maximum subnetwork size. These reactions are not connected, as in the subnetwork of the enrichment. This creates a distribution of scores. The distribution is then used to estimate a p-value for the solution found by the enrichment. The number of samples can be decided by the user, with more samples giving a better estimate of the distribution at increased runtime. The rationale behind sampling unconnected solutions is to estimate how much the connected (mechanistic) subnetwork scores compared to unconnected (non-mechanistic) solutions.

In the implementation for the LINEX^2^ web service, the local search is run multiple times (default=5), each with random seeds. The best result can then be optionally used as a seed for another local search run, to improve this result. The best score achieved in all local search runs is then returned to the user.

### 4.4 Reaction ratio plots

Visualizations of reaction ratios were performed for each lipid class reaction individually. Reaction ratios per sample are computed in the same as for the substrate-product change calculation, without averaging over all individuals:

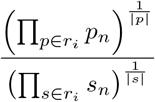

All ratios per experimental condition are compiled into a list and the density for each considered experimental condition is plotted.

### 4.5 Lipid moiety analysis

The (combined) abundance of lipid features was implemented inspired by the glycan substructure method by Bao et al. [55]. We used the same vectorization and weighting as the authors, but with lipid substructures as features. These were: headgroup, backbone, independent fatty acyls, sum length of fatty acyls, sum double bonds of fatty acyls, and fatty acyl hydroxylations. The features were weighted independently or in combination of pairs by occurrence in each lipid per sample. To find the most discriminative feature combinations, we train a regression model with sample groups as target variables and extract its coefficients. A summary of the workflow can be found in Supplementary Figure S5.

### 4.6 Lipid chain analysis

We implemented lipid chain analysis in python according to the proposed method by Mohamed, Molendijk, and Hill [53]. For each lipid class, lipid species with the same sum length of fatty acids are summed up per sample and a mean over all samples of one experimental condition is calculated. After that, the fold change between a selected control and e.g. a disease condition is calculated for each sum length per lipid class. The result is then plotted with an ascending fatty acid length on the x-axis, showing class-specific fatty acid length fold changes between conditions.

### 4.7 Webtool and data upload

The web service is built with the Django web framework (https://www.djangoproject.com/) in the python programming language (version 3.8, https://www.python.org/). PostgreSQL (https://www.postgresql.org/) is used as a back-end database for Django, to store data, networks, and all computed attributes. Cookies are used to connect a browser session to uploaded data, their corresponding computed networks, and analyses. For interactive network visualizations, vis-network [60] is used and other interactive plots are done with Plotly [61]. All other user-site functionalities are implemented in plain JavaScript. PDF versions of networks are generated with the NetworkX package [62] in conjunction with the matplotlib library [63]. The backend was implemented in python. To achieve compatibility across operating systems, LINEX^2^ can be built in a Docker environment. In the public LINEX^2^ version, uploaded user data is temporarily stored on our server for a certain time or can be deleted manually by the user (for further information see https://exbio.wzw.tum.de/linex/request-data-delete). However, using the provided Dockerfiles LINEX^2^ can also be easily run locally on any computer (for instructions check the source code repository). LINEX^2^ is free software, published under the aGPLv3 license. The source code is available at https://gitlab.lrz.de/lipitum-projects/linex. While we adapted the procedure to generate lipid species networks, the original LINEX version can still be accessed through the website (marked as version 1).

Identified and quantified lipidomics data with optional sample labels can be uploaded to LINEX^2^. Lipidomics data must be uploaded as a table with samples, lipids, and their corresponding concentrations/amounts. To convert lipids into our internal programming model, we recommend the LIPID MAPS nomenclature [64]. However, we integrated the LipidLynxX [65] software, which can convert multiple lipid nomenclatures, increasing the compatibility of LINEX^2^ with multiple formats. A tutorial is available on the website (https://exbio.wzw.tum.de/linex/tutorial).

### 4.8 Statistical measures implemented in LINEX^2^

To enable a combined visualization of the biochemical connections between lipid species and quantitative lipidomics measurements, LINEX^2^ offers the possibility to project different statistical metrics onto the species networks.

To characterize the changes in lipid levels between different experimental conditions, we provide unpaired parametric (t-test), non-parametric (Wilcoxon rank-sum test) test options, and a paired parametric test (Wilcoxon signed-rank test). All resulting p-values are automatically false-discovery rate (FDR) corrected using the Benjamini-Hochberg procedure [66]. Furthermore, fold changes are computed to showcase effect size. For the computation of these metrics, we used the scipy package [66, 67] in conjunction with the statsmodels package [68]. All measures can be visualized either as node sizes or node colors.

Measures for lipid connections (i.e. edges in the network) are correlation-based. Specifically, the options provided are spearman’s correlation and partial correlation. All correlations above a user-specified significance threshold (default = 0.05) are set to 0 automatically. Correlation values can be visualized in the network representation through edge colors.

### 4.9 Analyzed data sets

The lipidomics data for MOBAT7 WT and knockout mice were taken from Thangapandi et al. [38]. No further processing was done and the data was analyzed as provided by the authors. Data for the Adipo Atlas was used as provided in the supplement of Lange et al. [41]. The comparison of MCS to adipogenic cells is coming from the supplement of Levental et al. [51]. Lipid species measured in less than 50% of all samples were removed before analysis with LINEX^2^. For all data sets analyzed with LINEX^2^, HTML files with the LINEX^2^ output are available in the supplement.

## Data Availability Statement

LINEX is free software. Source code: GitLab (aGPLv3 License): https://gitlab.lrz.de/lipitum-projects/linex

Figure reproducibility: https://gitlab.lrz.de/lipitum-projects/linex

ALEX123 lipid classes and curated database reaction: https://gitlab.lrz.de/lipitum-projects/LINEX2_package/-/tree/master/LINEX2/data

## Author Contributions

JKP supervised the project and secured the funding. NK, TDR, and JKP planned and conceptualized the work. NK and TDR developed the web service. NK, OEL, and TDR designed and implemented the network enrichment procedure. LF, LK, and TDR parsed and curated the reaction databases, and implemented the network extension. NK and TDR applied, validated, and interpreted the approach on lipidomics data. NK, OEL, TDR, and JKP wrote the manuscript. All authors read, reviewed, and accepted the manuscript in its final form.

## Acknowledgments

This project was funded by the Bavarian State Ministry of Science and the Arts in the framework of the Bavarian Research Institute for Digital Transformation (bidt; JKP, NK, TDR: Junior Research Group LipiTUM; O.L.: Doctoral Fellow).

