## Supplementary material for "Lipid network and moiety analysis for revealing enzymatic dysregulation and mechanistic alterations from lipidomics data": LINEX2_supplement_v2.pdf

### Supplementary Figures

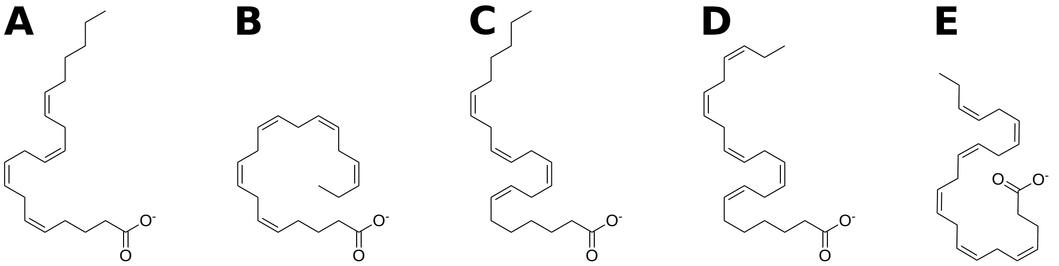


Figure S1: Fatty acid candidates for the acyltransferase reaction catalyzed by MBOAT7 predicted by the enrichment shown in Figure 4. **A** (5Z,8Z,11Z,14Z)-eicosatetraenoate (SwissLipids ID: SLM:000000296), **B** (5Z,8Z,11Z,14Z,17Z)-eicosapentaenoate (SwissLipids ID: SLM:000000409), **C** (7Z,10Z,13Z,16Z)-docosatetraenoate (SwissLipids ID: SLM:000001124), **D** (7Z,10Z,13Z,16Z,19Z)-docosapentaenoate (SwissLipids ID: SLM:000000929), and **E** (4Z,7Z,10Z,13Z,16Z,19Z)-docosahexaenoate (SwissLipids ID: SLM:000001084).

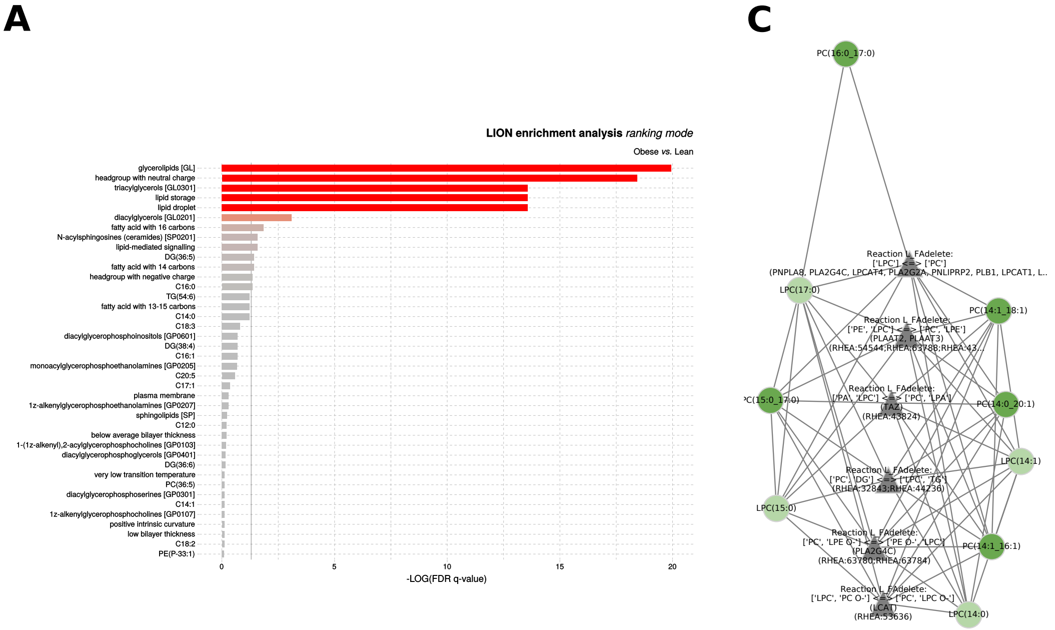


Figure S2: Analysis of cell culture data comparing mesenchymal stem cells and adipogenic cells. **A** LION enrichment with all lipids in the enrichment subnetwork (Main Figure 4A) as LION enrichment based on p-values from a student's t-test. **B** Enrichment result. The subnetwork only comprises lipids with a Phosphocholine head group.

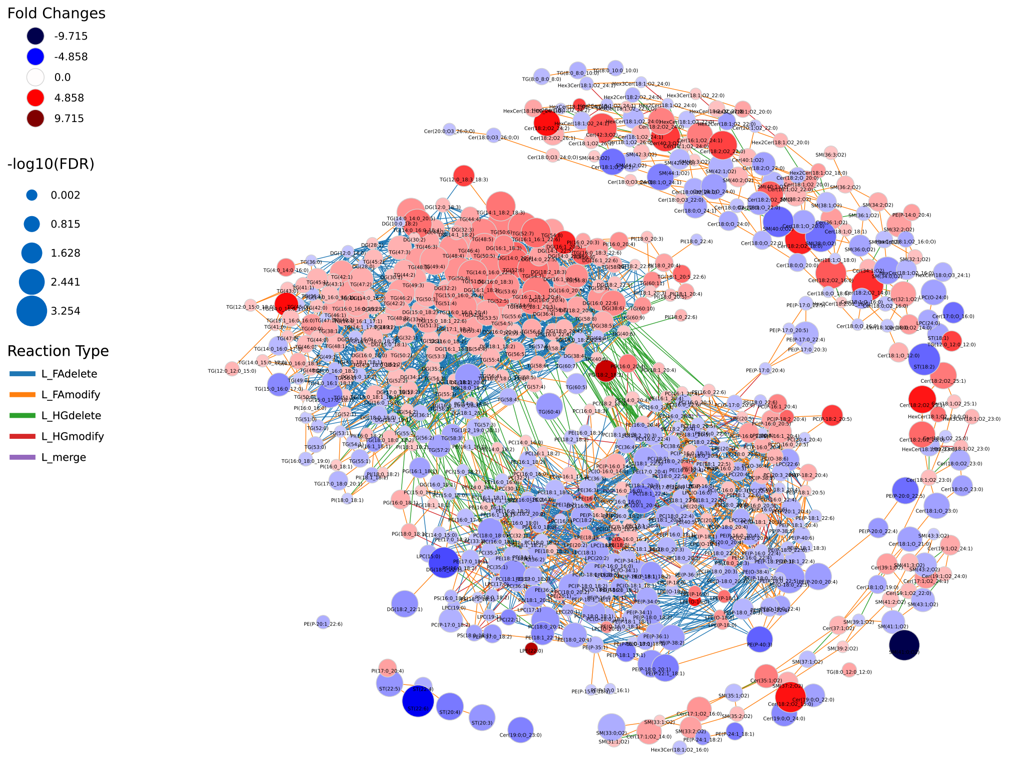


Figure S3: Annotated AdipoAtlas network. For a description of the network see Figure 4 and the Results section.

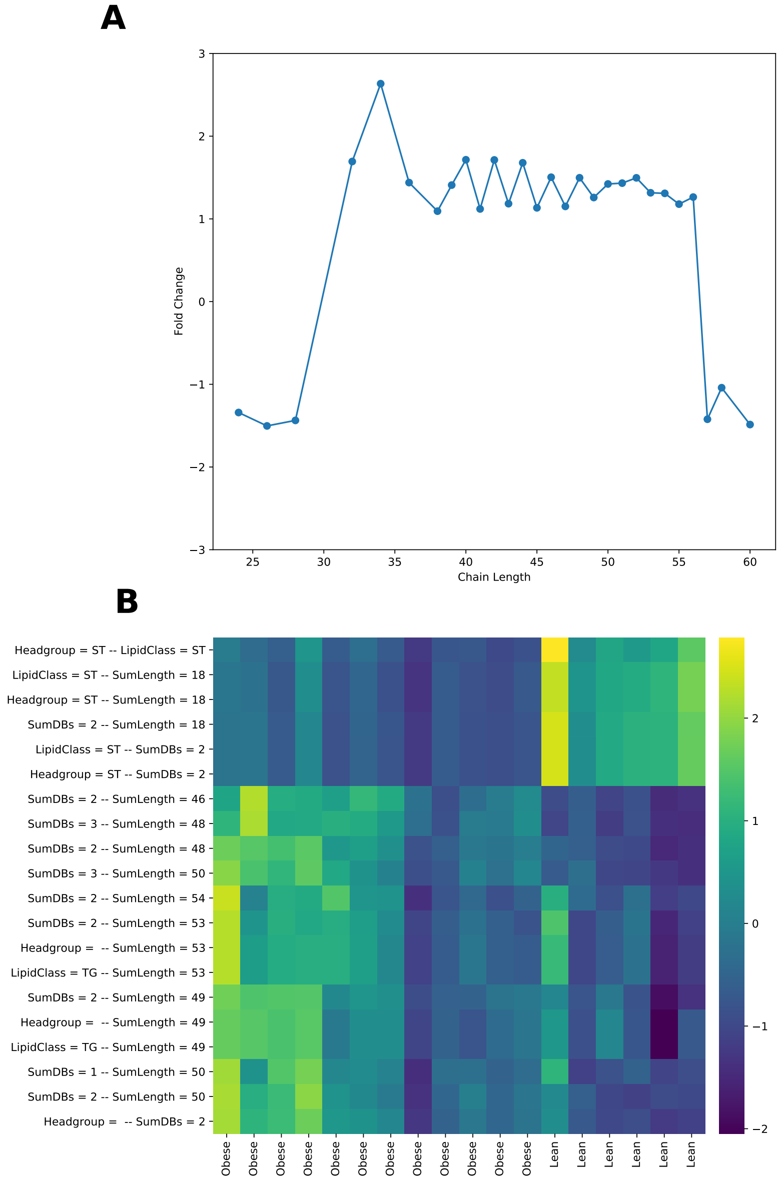


Figure S4: Additional results of LINEX$^2$ analyses on the AdipoAtlas data. **A** Results of the chain length analysis showing fold changes between obese and lean samples for the sum of all TG species per sum length. **B** Substructure analysis results, showing the 20 best property combinations. Features were selected by their absolute coefficients in a regression analysis with the obesity state (i.e. obese or lean) as the target variable. Selected properties include steryl esters, which show lower values in obese samples, and sum length combinations between 46 and 54 only found in TG species in the AdipoAtlas data.

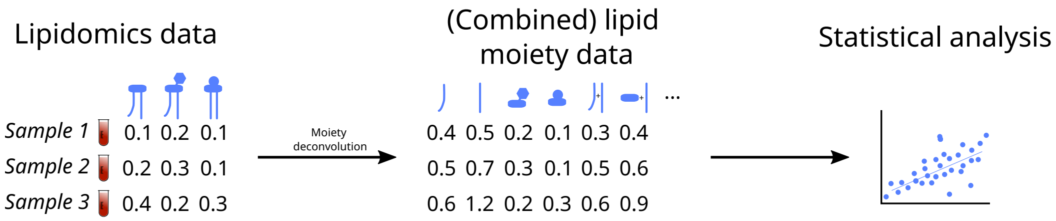


Figure S5: Workflow of lipid moiety analysis. Lipids from lipidomics data are deconvoluted into quantitative moieties. Moiety data is then used for statistical analysis, such as regression.

A

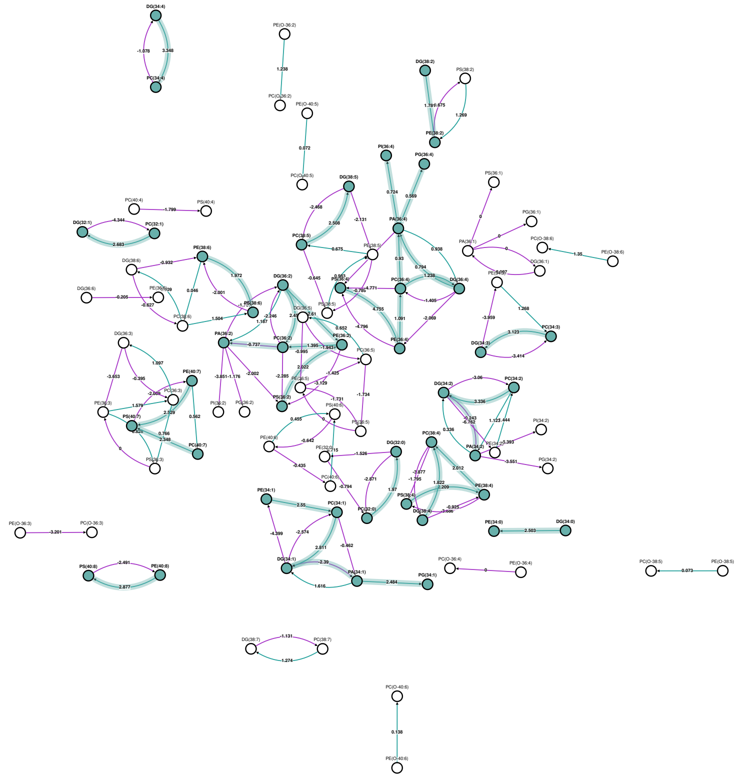

B

lipid species active reaction (WTHFCD vs KOHFCD)

| Reactions chains | Z-score | Predicted genes |
|---|---|---|
| PS(36:4) → PE(36:4) → PC(36:4) → PA(36:4) | 3.912 | *PISD, PEMT, PLD1, PLD2* |
| PS(36:4) → PE(36:4) → PC(36:4) → DG(36:4) → PA(36:4) → PI(36:4) | 3.847 | *PISD, PEMT, DGKA, DGKB, DGKD, DGKE, DGKG, DGKH, DGKI, DGKK, DGKQ, DGKZ, CDS1, CDS2, CDIPT* |
| PS(36:4) → PE(36:4) → PC(36:4) → DG(36:4) → PA(36:4) → PG(36:4) | 3.778 | *PISD, PEMT, DGKA, DGKB, DGKD, DGKE, DGKG, DGKH, DGKI, DGKK, DGKQ, DGKZ, CDS1, CDS2, PTPMT1* |
| DG(34:4) → PC(34:4) | 3.348 | *CHPT1* |
| PC(34:3) → DG(34:3) | 3.123 | No genes have yet been identified |
| PS(38:4) → PE(38:4) → PC(38:4) | 2.985 | *PISD, PEMT* |
| PE(40:8) → PS(40:8) | 2.877 | *PTDSS2* |
| PC(34:1) → DG(34:1) | 2.811 | No genes have yet been identified |
| PE(34:1) → PC(34:1) → DG(34:1) → PA(34:1) → PG(34:1) | 2.728 | *PEMT, DGKA, DGKB, DGKD, DGKE, DGKG, DGKH, DGKI, DGKK, DGKQ, DGKZ, CDS1, CDS2, PTPMT1* |
| PC(32:1) → DG(32:1) | 2.683 | No genes have yet been identified |
| PE(40:7) → PS(40:7) | 2.529 | *PTDSS2* |
| PC(38:5) → DG(38:5) | 2.508 | No genes have yet been identified |
| DG(34:0) → PE(34:0) | 2.503 | *CEPT1* |
| PA(34:1) → PG(34:1) | 2.484 | *CDS1, CDS2, PTPMT1* |
| DG(36:2) → PC(36:2) | 2.456 | *CHPT1* |
| PS(36:2) → PE(36:2) → PC(36:2) | 2.416 | *PISD, PEMT* |
| PC(40:7) → PS(40:7) | 2.348 | *PTDSS1* |
| PC(34:2) → DG(34:2) → PA(34:2) | 2.187 | *DGKA, DGKB, DGKD, DGKE, DGKG, DGKH, DGKI, DGKK, DGKQ, DGKZ* |
| PE(38:4) → PC(38:4) | 2.012 | *PEMT* |
| PE(38:6) → PS(38:6) | 1.972 | *PTDSS2* |
| PC(32:0) → DG(32:0) | 1.97 | No genes have yet been identified |
| DG(36:2) → PE(36:2) → PC(36:2) → PA(36:2) | 1.887 | *CEPT1, PEMT, PLD1, PLD2* |
| DG(38:4) → PC(38:4) | 1.822 | *CHPT1* |
| DG(38:2) → PE(38:2) | 1.781 | *CEPT1* |

Figure S6: BioPAN results for the MBOAT7 knock-out data. **A** Computed lipid species network with z-scores for active reactions. **B** Predicted active lipid species reactions with corresponding z-scores and predicted genes.

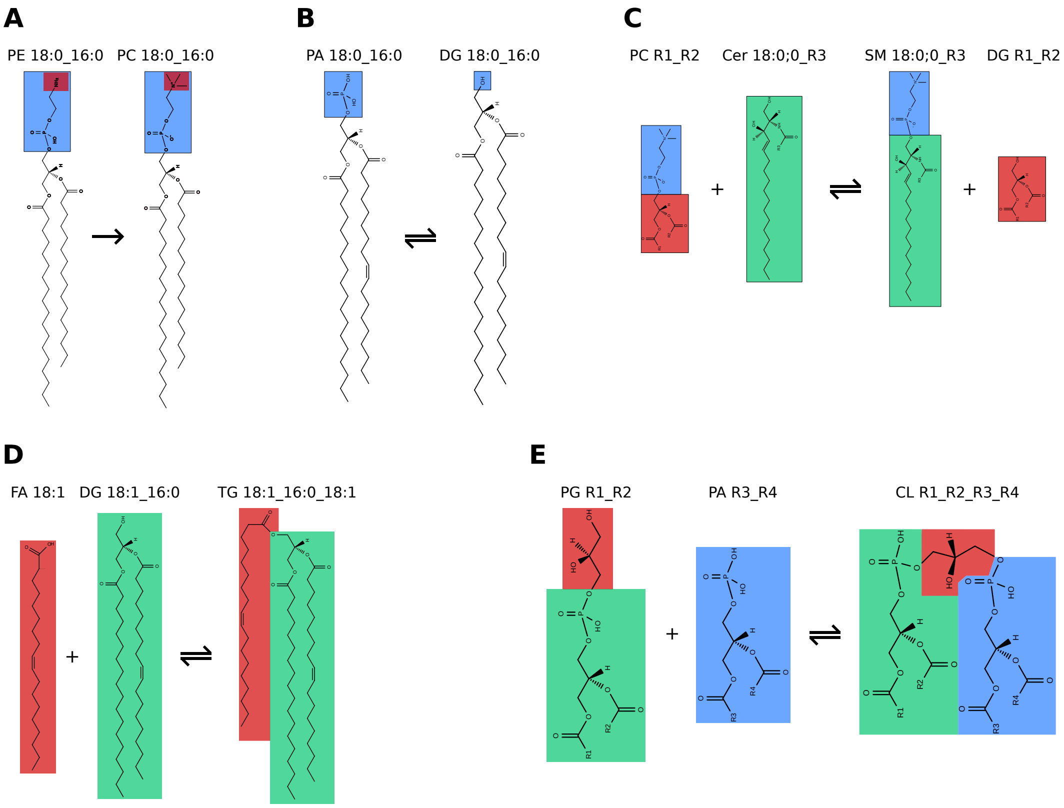


Figure S7: Schematic depiction of common biochemical lipid modifications/transformations. **A** Tri-methylation of the ethanolamine moiety of a PE produces the corresponding (i.e. same fatty acid combination) PC species **B** De-phosphorylation of a PA transforms it into a DG. **C** The phosphocholine group of PC is transferred onto a Cer, the PC-corresponding DG and the Cer-corresponding SM species. **D** Esterification of a free fatty acid onto a DG produces a TG. **E** PG and PA are linked via esterification of the glycerol and the phosphate group. The Cardiolipin product contains the same fatty acid residues as the substrates.
