## Supplementary material for "Lipid network and moiety analysis for revealing enzymatic dysregulation and mechanistic alterations from lipidomics data": LINEX_network_AdipoAtlas.html

LINEX - LipidNetworkExplorer


- Visit the LINEX website

Network

Network Enrichment

Lipidome Summary

Version 2

##### Network Options

**Network Type**

Native Network
Enzyme Network
Native Molecular Network
Enzyme Molecular Network

Recompute Network
  


---


**Node Colours**

Lipid Class
Desaturation
Chain Length
C Index
DB Index
-log10(FDR)
Fold Changes
Closeness Centrality
OH Index
Betweenness Centrality
Degree
Hydroxylation

**Edge Colours**

Reaction Type
Correlations
Correlation Changes

**Node Sizes**

-log10(FDR)
Fold Changes
Closeness Centrality
OH Index
Betweenness Centrality
Chain Length
Degree
Desaturation
DB Index
C Index
Hydroxylation


---


**Comparison**

Obese\_Lean

**Group**

Obese
Lean


---


**Find Lipid Species**


Find
  

**Find by Substring**

Find


---


  
  
**Shown reaction types**

all
only Class reactions
only FA reactions

  
  
**Significance Threshold**

Update
  

**Highlight**

**Enable physics**

**Unified Scales**

---


Download View

Downloads the network as a .json file, which can be uploaded to 127.0.0.1:8000/download to generate a pdf view

##### Legend

Hide Legend Navigation

##### Network

0%


Enrichment settings not available in downloaded file

Lipidome Overview


Substructure Analysis


Chain Length Analysis

##### Substructure Features

##### Regression-Based Feature Selection

| Lean | | |
| --- | --- | --- |
| **Feature** | **Coefficient** | **Absolute Coefficient** |
| Headgroup = -- SumLength = 49 | 0.0 | 0.0 |
| LipidClass = TG -- SumLength = 49 | 0.0 | 0.0 |
| SumDBs = 1 -- SumLength = 50 | -0.0 | 0.0 |
| Headgroup = ST -- LipidClass = ST | 0.0 | 0.0 |
| SumDBs = 2 -- SumLength = 46 | -0.0 | 0.0 |
| LipidClass = TG -- SumLength = 53 | -0.0 | 0.0 |
| Headgroup = -- SumLength = 53 | -0.0 | 0.0 |
| SumDBs = 2 -- SumLength = 53 | -0.0 | 0.0 |
| SumDBs = 3 -- SumLength = 48 | -0.0 | 0.0 |
| SumDBs = 2 -- SumLength = 48 | 0.0 | 0.0 |
| SumDBs = 2 -- SumLength = 50 | 0.0 | 0.0 |
| LipidClass = ST -- SumLength = 18 | 0.0 | 0.0 |
| Headgroup = ST -- SumLength = 18 | 0.0 | 0.0 |
| SumDBs = 2 -- SumLength = 54 | -0.0 | 0.0 |
| SumDBs = 2 -- SumLength = 18 | 0.0 | 0.0 |
| Headgroup = ST -- SumDBs = 2 | 0.0 | 0.0 |
| LipidClass = ST -- SumDBs = 2 | 0.0 | 0.0 |
| SumDBs = 3 -- SumLength = 50 | 0.0 | 0.0 |
| LipidClass = TG -- SumDBs = 2 | -0.0 | 0.0 |
| SumDBs = 1 -- SumLength = 46 | 0.0 | 0.0 |
| SumDBs = 2 -- SumLength = 49 | 0.0 | 0.0 |
| Headgroup = -- SumDBs = 2 | -0.0 | 0.0 |
| SumDBs = 4 -- SumLength = 50 | -0.0 | 0.0 |
| SumDBs = 5 -- SumLength = 56 | 0.0 | 0.0 |
| LipidClass = TG -- SumLength = 45 | 0.0 | 0.0 |
| Headgroup = -- SumLength = 45 | 0.0 | 0.0 |
| SumDBs = 6 -- SumLength = 58 | -0.0 | 0.0 |
| SumDBs = 4 -- SumLength = 54 | 0.0 | 0.0 |
| LipidClass = TG -- SumDBs = 1 | 0.0 | 0.0 |
| Headgroup = -- SumLength = 46 | -0.0 | 0.0 |
| Headgroup = -- SumDBs = 1 | 0.0 | 0.0 |
| LipidClass = TG -- SumLength = 46 | -0.0 | 0.0 |
| Headgroup = -- SumLength = 51 | 0.0 | 0.0 |
| LipidClass = TG -- SumLength = 51 | 0.0 | 0.0 |
| SumDBs = 3 -- SumLength = 51 | 0.0 | 0.0 |
| SumDBs = 2 -- SumLength = 56 | -0.0 | 0.0 |
| SumDBs = 1 -- SumLength = 47 | 0.0 | 0.0 |
| Headgroup = -- SumDBs = 3 | 0.0 | 0.0 |
| SumDBs = 5 -- SumLength = 54 | -0.0 | 0.0 |
| Headgroup = -- SumLength = 55 | -0.0 | 0.0 |
| LipidClass = TG -- SumLength = 55 | -0.0 | 0.0 |
| SumDBs = 2 -- SumLength = 52 | 0.0 | 0.0 |
| Headgroup = -- SumLength = 56 | 0.0 | 0.0 |
| LipidClass = TG -- SumLength = 56 | 0.0 | 0.0 |
| SumDBs = 0 -- SumLength = 45 | 0.0 | 0.0 |
| SumDBs = 2 -- SumLength = 51 | -0.0 | 0.0 |
| SumDBs = 3 -- SumLength = 58 | 0.0 | 0.0 |
| SumDBs = 1 -- SumLength = 51 | 0.0 | 0.0 |
| SumDBs = 3 -- SumLength = 55 | -0.0 | 0.0 |
| LipidClass = TG -- SumDBs = 3 | 0.0 | 0.0 |
| SumDBs = 0 -- SumLength = 47 | -0.0 | 0.0 |
| Headgroup = -- LipidClass = TG | 0.0 | 0.0 |
| SumDBs = 4 -- SumLength = 58 | 0.0 | 0.0 |
| LipidClass = TG -- SumDBs = 6 | -0.0 | 0.0 |
| SumDBs = 1 -- SumLength = 44 | -0.0 | 0.0 |
| SumDBs = 4 -- SumLength = 52 | -0.0 | 0.0 |
| SumDBs = 3 -- SumLength = 54 | 0.0 | 0.0 |
| Headgroup = -- SumDBs = 6 | -0.0 | 0.0 |
| LipidClass = TG -- SumDBs = 9 | 0.0 | 0.0 |
| Headgroup = -- SumDBs = 9 | 0.0 | 0.0 |
| LipidClass = ST -- SumLength = 20 | 0.0 | 0.0 |
| Headgroup = ST -- SumLength = 20 | 0.0 | 0.0 |
| LipidClass = TG -- SumLength = 47 | 0.0 | 0.0 |
| Headgroup = -- SumLength = 47 | 0.0 | 0.0 |
| Headgroup = -- SumLength = 44 | -0.0 | 0.0 |
| SumDBs = 0 -- SumLength = 36 | -0.0 | 0.0 |
| Headgroup = -- SumLength = 52 | -0.0 | 0.0 |
| LipidClass = TG -- SumLength = 52 | -0.0 | 0.0 |
| LipidClass = TG -- SumLength = 44 | -0.0 | 0.0 |
| SumDBs = 1 -- SumLength = 49 | 0.0 | 0.0 |
| Headgroup = -- SumLength = 50 | -0.0 | 0.0 |
| LipidClass = TG -- SumLength = 50 | -0.0 | 0.0 |
| Headgroup = -- SumLength = 36 | -0.0 | 0.0 |
| LipidClass = TG -- SumLength = 36 | -0.0 | 0.0 |
| SumDBs = 9 -- SumLength = 58 | 0.0 | 0.0 |
| LipidClass = TG -- SumLength = 40 | 0.0 | 0.0 |
| Headgroup = -- SumLength = 40 | 0.0 | 0.0 |
| SumDBs = 4 -- SumLength = 53 | 0.0 | 0.0 |
| Headgroup = -- SumLength = 54 | -0.0 | 0.0 |
| LipidClass = TG -- SumLength = 54 | -0.0 | 0.0 |
| SumDBs = 3 -- SumLength = 49 | 0.0 | 0.0 |
| SumDBs = 7 -- SumLength = 58 | -0.0 | 0.0 |
| SumDBs = 7 -- SumLength = 56 | 0.0 | 0.0 |
| SumDBs = 5 -- SumLength = 52 | -0.0 | 0.0 |
| SumDBs = 0 -- SumLength = 53 | -0.0 | 0.0 |
| SumDBs = 4 -- SumLength = 48 | -0.0 | 0.0 |
| SumDBs = 1 -- SumLength = 45 | 0.0 | 0.0 |
| Headgroup = -- SumDBs = 0 | -0.0 | 0.0 |
| SumDBs = 3 -- SumLength = 46 | -0.0 | 0.0 |
| LipidClass = TG -- SumLength = 60 | 0.0 | 0.0 |
| Headgroup = -- SumLength = 60 | 0.0 | 0.0 |
| SumDBs = 3 -- SumLength = 52 | -0.0 | 0.0 |
| Headgroup = -- SumLength = 43 | 0.0 | 0.0 |
| SumDBs = 6 -- SumLength = 54 | 0.0 | 0.0 |
| LipidClass = TG -- SumLength = 43 | 0.0 | 0.0 |
| SumDBs = 5 -- SumLength = 58 | -0.0 | 0.0 |
| SumDBs = 1 -- SumLength = 53 | -0.0 | 0.0 |
| SumDBs = 0 -- SumLength = 46 | 0.0 | 0.0 |
| LipidClass = TG -- SumDBs = 0 | -0.0 | 0.0 |
| SumDBs = 2 -- SumLength = 47 | 0.0 | 0.0 |
| Headgroup = ST -- SumDBs = 3 | 0.0 | 0.0 |
| LipidClass = ST -- SumDBs = 3 | 0.0 | 0.0 |
| SumDBs = 3 -- SumLength = 20 | 0.0 | 0.0 |
| LipidClass = TG -- SumLength = 48 | -0.0 | 0.0 |
| Headgroup = -- SumLength = 48 | -0.0 | 0.0 |
| SumDBs = 4 -- SumLength = 20 | 0.0 | 0.0 |
| LipidClass = TG -- SumLength = 42 | 0.0 | 0.0 |
| Headgroup = -- SumLength = 42 | 0.0 | 0.0 |
| LipidClass = ST -- SumDBs = 4 | 0.0 | 0.0 |
| Headgroup = ST -- SumDBs = 4 | 0.0 | 0.0 |
| LipidClass = TG -- SumDBs = 8 | 0.0 | 0.0 |
| Headgroup = -- SumDBs = 8 | 0.0 | 0.0 |
| SumDBs = 1 -- SumLength = 48 | 0.0 | 0.0 |
| SumDBs = 4 -- SumLength = 60 | 0.0 | 0.0 |
| SumDBs = 0 -- SumLength = 51 | -0.0 | 0.0 |
| SumDBs = 1 -- SumLength = 55 | -0.0 | 0.0 |
| SumDBs = 0 -- SumLength = 40 | 0.0 | 0.0 |
| SumDBs = 4 -- SumLength = 51 | 0.0 | 0.0 |
| Headgroup = ST -- SumLength = 22 | 0.0 | 0.0 |
| LipidClass = ST -- SumLength = 22 | 0.0 | 0.0 |
| SumDBs = 1 -- SumLength = 42 | 0.0 | 0.0 |
| SumDBs = 0 -- SumLength = 43 | 0.0 | 0.0 |
| SumDBs = 8 -- SumLength = 54 | 0.0 | 0.0 |
| Headgroup = PS -- LipidClass = PS | 0.0 | 0.0 |
| SumDBs = 3 -- SumLength = 56 | -0.0 | 0.0 |
| SumDBs = 0 -- SumLength = 49 | 0.0 | 0.0 |
| Headgroup = -- SumDBs = 4 | 0.0 | 0.0 |
| SumDBs = 1 -- SumLength = 40 | 0.0 | 0.0 |
| LipidClass = TG -- SumDBs = 7 | 0.0 | 0.0 |
| SumDBs = 4 -- SumLength = 55 | -0.0 | 0.0 |
| SumDBs = 5 -- SumLength = 53 | 0.0 | 0.0 |
| Headgroup = -- SumDBs = 7 | 0.0 | 0.0 |
| LipidClass = TG -- SumLength = 41 | 0.0 | 0.0 |
| Headgroup = -- SumLength = 41 | 0.0 | 0.0 |
| Headgroup = PS -- SumLength = 36 | 0.0 | 0.0 |
| LipidClass = PS -- SumLength = 36 | 0.0 | 0.0 |
| SumDBs = 0 -- SumLength = 50 | -0.0 | 0.0 |
| LipidClass = Cer -- SumLength = 38 | -0.0 | 0.0 |
| SumDBs = 6 -- SumLength = 56 | 0.0 | 0.0 |
| Headgroup = -- SumDBs = 5 | 0.0 | 0.0 |
| SumDBs = 6 -- SumLength = 22 | 0.0 | 0.0 |
| Headgroup = PC -- SumLength = 34 | 0.0 | 0.0 |
| Headgroup = ST -- SumDBs = 6 | 0.0 | 0.0 |
| LipidClass = ST -- SumDBs = 6 | 0.0 | 0.0 |
| LipidClass = TG -- SumDBs = 5 | 0.0 | 0.0 |
| SumDBs = 5 -- SumLength = 50 | 0.0 | 0.0 |
| LipidClass = TG -- SumDBs = 10 | 0.0 | 0.0 |
| Headgroup = -- SumDBs = 10 | 0.0 | 0.0 |
| SumDBs = 5 -- SumLength = 22 | 0.0 | 0.0 |
| Headgroup = ST -- SumDBs = 5 | 0.0 | 0.0 |
| LipidClass = ST -- SumDBs = 5 | 0.0 | 0.0 |
| SumDBs = 0 -- SumLength = 16 | -0.0 | 0.0 |
| SumDBs = 10 -- SumLength = 58 | 0.0 | 0.0 |
| SumDBs = 1 -- SumLength = 43 | 0.0 | 0.0 |
| SumDBs = 1 -- SumLength = 36 | 0.0 | 0.0 |
| SumDBs = 0 -- SumLength = 52 | -0.0 | 0.0 |
| LipidClass = PS -- SumDBs = 1 | 0.0 | 0.0 |
| Headgroup = PS -- SumDBs = 1 | 0.0 | 0.0 |
| SumDBs = 2 -- SumLength = 45 | 0.0 | 0.0 |
| Headgroup = PC -- LipidClass = PC | 0.0 | 0.0 |
| Headgroup = -- SumLength = 58 | 0.0 | 0.0 |
| LipidClass = TG -- SumLength = 58 | 0.0 | 0.0 |
| Headgroup = PC -- LipidClass = LPC | -0.0 | 0.0 |
| Headgroup = PE -- SumDBs = 0 | -0.0 | 0.0 |
| Headgroup = PC -- LipidClass = PC O- | 0.0 | 0.0 |
| SumDBs = 2 -- SumLength = 40 | 0.0 | 0.0 |
| LipidClass = LPC -- SumDBs = 0 | -0.0 | 0.0 |
| Headgroup = PC -- SumDBs = 0 | -0.0 | 0.0 |
| SumDBs = 0 -- SumLength = 41 | 0.0 | 0.0 |
| SumDBs = 2 -- SumLength = 55 | 0.0 | 0.0 |
| Headgroup = PE -- LipidClass = PE O- | 0.0 | 0.0 |
| Headgroup = PE -- SumDBs = 4 | 0.0 | 0.0 |
| LipidClass = TG -- SumLength = 57 | -0.0 | 0.0 |
| Headgroup = -- SumLength = 57 | -0.0 | 0.0 |
| Headgroup = PE -- SumLength = 18 | -0.0 | 0.0 |
| SumDBs = 4 -- SumLength = 36 | 0.0 | 0.0 |
| SumDBs = 0 -- SumLength = 18 | -0.0 | 0.0 |
| SumDBs = 5 -- SumLength = 55 | 0.0 | 0.0 |
| Headgroup = PC -- SumDBs = 1 | 0.0 | 0.0 |
| Headgroup = PC -- SumDBs = 2 | 0.0 | 0.0 |
| Headgroup = PE -- LipidClass = LPE O- | -0.0 | 0.0 |
| LipidClass = TG -- SumDBs = 4 | 0.0 | 0.0 |
| Headgroup = PC -- SumLength = 36 | 0.0 | 0.0 |
| Headgroup = -- SumLength = 38 | 0.0 | 0.0 |
| LipidClass = TG -- SumLength = 38 | 0.0 | 0.0 |
| SumDBs = 7 -- SumLength = 54 | 0.0 | 0.0 |
| Headgroup = PE -- SumLength = 38 | 0.0 | 0.0 |
| Headgroup = -- SumLength = 34 | -0.0 | 0.0 |
| SumDBs = 7 -- SumLength = 52 | 0.0 | 0.0 |
| SumDBs = 4 -- SumLength = 38 | 0.0 | 0.0 |
| SumDBs = 1 -- SumLength = 34 | 0.0 | 0.0 |
| Headgroup = PC -- LipidClass = SM | 0.0 | 0.0 |
| SumDBs = 8 -- SumLength = 56 | 0.0 | 0.0 |
| Headgroup = PC -- SumLength = 16 | -0.0 | 0.0 |
| LipidClass = PE O- -- SumDBs = 4 | 0.0 | 0.0 |
| LipidClass = SM -- SumDBs = 1 | 0.0 | 0.0 |
| LipidClass = LPC -- SumLength = 16 | -0.0 | 0.0 |
| SumDBs = 2 -- SumLength = 57 | -0.0 | 0.0 |
| Headgroup = PE -- SumDBs = 1 | -0.0 | 0.0 |
| SumDBs = 0 -- SumLength = 34 | -0.0 | 0.0 |
| Headgroup = PE -- LipidClass = PE | 0.0 | 0.0 |
| SumDBs = 3 -- SumLength = 44 | 0.0 | 0.0 |
| SumDBs = 3 -- SumLength = 47 | 0.0 | 0.0 |
| SumDBs = 0 -- SumLength = 48 | 0.0 | 0.0 |
| Headgroup = PE -- SumDBs = 5 | 0.0 | 0.0 |
| Headgroup = PE -- LipidClass = LPE | -0.0 | 0.0 |
| LipidClass = LPE O- -- SumDBs = 0 | -0.0 | 0.0 |
| LipidClass = TG -- SumLength = 34 | -0.0 | 0.0 |
| SumDBs = 2 -- SumLength = 42 | 0.0 | 0.0 |
| LipidClass = PC -- SumLength = 34 | 0.0 | 0.0 |
| SumDBs = 6 -- SumLength = 52 | -0.0 | 0.0 |
| Headgroup = PE -- SumLength = 16 | -0.0 | 0.0 |
| Headgroup = PC -- SumDBs = 4 | 0.0 | 0.0 |
| SumDBs = 9 -- SumLength = 56 | 0.0 | 0.0 |
| Headgroup = -- SumLength = 28 | -0.0 | 0.0 |
| SumDBs = 0 -- SumLength = 28 | -0.0 | 0.0 |
| LipidClass = TG -- SumLength = 28 | -0.0 | 0.0 |
| SumDBs = 1 -- SumLength = 38 | 0.0 | 0.0 |
| Headgroup = PS -- SumDBs = 2 | 0.0 | 0.0 |
| LipidClass = PS -- SumDBs = 2 | 0.0 | 0.0 |
| SumDBs = 8 -- SumLength = 58 | -0.0 | 0.0 |
| SumDBs = 4 -- SumLength = 56 | -0.0 | 0.0 |
| LipidClass = PE O- -- SumLength = 38 | 0.0 | 0.0 |
| SumDBs = 1 -- SumLength = 41 | 0.0 | 0.0 |
| LipidClass = LPE -- SumDBs = 0 | -0.0 | 0.0 |
| SumDBs = 2 -- SumLength = 34 | 0.0 | 0.0 |
| LipidClass = PC O- -- SumLength = 36 | 0.0 | 0.0 |
| LipidClass = LPE O- -- SumLength = 18 | -0.0 | 0.0 |
| SumDBs = 3 -- SumLength = 53 | 0.0 | 0.0 |
| SumDBs = 6 -- SumLength = 50 | 0.0 | 0.0 |
| SumDBs = 2 -- SumLength = 36 | 0.0 | 0.0 |
| Headgroup = PC -- SumLength = 38 | 0.0 | 0.0 |
| LipidClass = LPE -- SumLength = 18 | -0.0 | 0.0 |
| Headgroup = PC -- SumLength = 18 | -0.0 | 0.0 |
| LipidClass = PE O- -- SumDBs = 5 | 0.0 | 0.0 |
| LipidClass = LPC -- SumLength = 18 | -0.0 | 0.0 |
| SumDBs = 4 -- SumLength = 49 | 0.0 | 0.0 |
| LipidClass = PC -- SumDBs = 2 | 0.0 | 0.0 |
| SumDBs = 5 -- SumLength = 60 | 0.0 | 0.0 |
| SumDBs = 5 -- SumLength = 38 | 0.0 | 0.0 |
| LipidClass = LPE O- -- SumLength = 16 | -0.0 | 0.0 |
| LipidClass = TG -- SumLength = 26 | -0.0 | 0.0 |
| Headgroup = -- SumLength = 26 | -0.0 | 0.0 |
| SumDBs = 0 -- SumLength = 26 | -0.0 | 0.0 |
| LipidClass = PC O- -- SumLength = 34 | 0.0 | 0.0 |
| LipidClass = PC O- -- SumDBs = 2 | 0.0 | 0.0 |
| Headgroup = PE -- SumLength = 36 | 0.0 | 0.0 |
| LipidClass = PC -- SumLength = 36 | 0.0 | 0.0 |
| Headgroup = PE -- SumDBs = 3 | 0.0 | 0.0 |
| Headgroup = -- LipidClass = DG | -0.0 | 0.0 |
| LipidClass = SM -- SumLength = 34 | 0.0 | 0.0 |
| LipidClass = PC O- -- SumDBs = 4 | 0.0 | 0.0 |
| SumDBs = 9 -- SumLength = 60 | 0.0 | 0.0 |
| Headgroup = PE -- SumLength = 37 | 0.0 | 0.0 |
| LipidClass = PE -- SumLength = 38 | 0.0 | 0.0 |
| LipidClass = PE O- -- SumLength = 37 | 0.0 | 0.0 |
| SumDBs = 0 -- SumLength = 44 | 0.0 | 0.0 |
| SumDBs = 8 -- SumLength = 60 | 0.0 | 0.0 |
| LipidClass = TG -- SumLength = 39 | 0.0 | 0.0 |
| SumDBs = 0 -- SumLength = 39 | 0.0 | 0.0 |
| Headgroup = -- SumLength = 39 | 0.0 | 0.0 |
| SumDBs = 3 -- SumLength = 57 | -0.0 | 0.0 |
| Headgroup = ST -- SumDBs = 1 | 0.0 | 0.0 |
| LipidClass = ST -- SumDBs = 1 | 0.0 | 0.0 |
| SumDBs = 2 -- SumLength = 38 | 0.0 | 0.0 |
| LipidClass = PE O- -- SumDBs = 2 | -0.0 | 0.0 |
| Headgroup = PS -- SumDBs = 4 | 0.0 | 0.0 |
| LipidClass = PS -- SumDBs = 4 | 0.0 | 0.0 |
| LipidClass = PE O- -- SumDBs = 3 | 0.0 | 0.0 |
| LipidClass = PE O- -- SumLength = 36 | 0.0 | 0.0 |
| Headgroup = PS -- SumLength = 38 | 0.0 | 0.0 |
| LipidClass = PS -- SumLength = 38 | 0.0 | 0.0 |
| SumDBs = 7 -- SumLength = 55 | 0.0 | 0.0 |
| Headgroup = PE -- SumDBs = 2 | -0.0 | 0.0 |
| SumDBs = 3 -- SumLength = 37 | 0.0 | 0.0 |
| LipidClass = PE O- -- SumDBs = 1 | -0.0 | 0.0 |
| LipidClass = LPE O- -- SumDBs = 1 | -0.0 | 0.0 |
| Headgroup = PC -- SumLength = 40 | 0.0 | 0.0 |
| SumDBs = 4 -- SumLength = 22 | -0.0 | 0.0 |
| LipidClass = PE -- SumDBs = 4 | 0.0 | 0.0 |
| SumDBs = 6 -- SumLength = 38 | 0.0 | 0.0 |
| Headgroup = PS -- SumLength = 34 | 0.0 | 0.0 |
| LipidClass = PS -- SumLength = 34 | 0.0 | 0.0 |
| Headgroup = PE -- SumLength = 40 | 0.0 | 0.0 |
| Headgroup = PC -- SumDBs = 3 | 0.0 | 0.0 |
| LipidClass = SM -- SumLength = 40 | 0.0 | 0.0 |
| LipidClass = PC -- SumDBs = 1 | 0.0 | 0.0 |
| LipidClass = PC -- SumLength = 38 | 0.0 | 0.0 |
| SumDBs = 6 -- SumLength = 55 | 0.0 | 0.0 |
| SumDBs = 1 -- SumLength = 18 | -0.0 | 0.0 |
| LipidClass = PC -- SumDBs = 4 | 0.0 | 0.0 |
| SumDBs = 5 -- SumLength = 36 | 0.0 | 0.0 |
| LipidClass = PE O- -- SumLength = 34 | -0.0 | 0.0 |
| Headgroup = -- LipidClass = Cer | -0.0 | 0.0 |
| SumDBs = 1 -- SumLength = 52 | -0.0 | 0.0 |
| SumDBs = 5 -- SumLength = 48 | 0.0 | 0.0 |
| Headgroup = PE -- SumDBs = 6 | 0.0 | 0.0 |
| Headgroup = -- SumLength = 32 | -0.0 | 0.0 |
| SumDBs = 5 -- SumLength = 51 | 0.0 | 0.0 |
| LipidClass = PE -- SumDBs = 5 | 0.0 | 0.0 |
| LipidClass = TG -- SumLength = 24 | 0.0 | 0.0 |
| Headgroup = -- SumLength = 24 | 0.0 | 0.0 |
| SumDBs = 0 -- SumLength = 24 | 0.0 | 0.0 |
| LipidClass = PC O- -- SumLength = 38 | 0.0 | 0.0 |
| SumDBs = 3 -- SumLength = 36 | 0.0 | 0.0 |
| SumDBs = 6 -- SumLength = 48 | 0.0 | 0.0 |
| LipidClass = DG -- SumDBs = 2 | -0.0 | 0.0 |
| Headgroup = PC -- SumDBs = 5 | 0.0 | 0.0 |
| LipidClass = LPE -- SumLength = 16 | -0.0 | 0.0 |
| SumDBs = 5 -- SumLength = 40 | 0.0 | 0.0 |
| LipidClass = LPE -- SumDBs = 1 | -0.0 | 0.0 |
| LipidClass = DG -- SumLength = 34 | -0.0 | 0.0 |
| LipidClass = PE -- SumLength = 40 | 0.0 | 0.0 |
| Headgroup = PE -- SumLength = 34 | -0.0 | 0.0 |
| SumDBs = 6 -- SumLength = 40 | 0.0 | 0.0 |
| LipidClass = PC O- -- SumDBs = 1 | 0.0 | 0.0 |
| Headgroup = PC -- SumDBs = 6 | 0.0 | 0.0 |
| LipidClass = PC -- SumDBs = 3 | 0.0 | 0.0 |
| LipidClass = LPC -- SumDBs = 1 | -0.0 | 0.0 |
| LipidClass = TG -- SumLength = 32 | -0.0 | 0.0 |
| LipidClass = DG -- SumLength = 36 | -0.0 | 0.0 |
| LipidClass = PC O- -- SumDBs = 3 | 0.0 | 0.0 |
| LipidClass = Cer -- SumDBs = 1 | -0.0 | 0.0 |
| SumDBs = 1 -- SumLength = 54 | 0.0 | 0.0 |
| LipidClass = PE O- -- SumLength = 40 | 0.0 | 0.0 |
| LipidClass = DG -- SumDBs = 1 | -0.0 | 0.0 |
| LipidClass = PE O- -- SumDBs = 6 | 0.0 | 0.0 |
| LipidClass = PE -- SumLength = 36 | 0.0 | 0.0 |
| SumDBs = 3 -- SumLength = 38 | 0.0 | 0.0 |
| LipidClass = PC -- SumDBs = 5 | 0.0 | 0.0 |
| SumDBs = 0 -- SumLength = 42 | -0.0 | 0.0 |
| SumDBs = 4 -- SumLength = 46 | -0.0 | 0.0 |
| LipidClass = PE -- SumDBs = 6 | 0.0 | 0.0 |
| SumDBs = 6 -- SumLength = 53 | 0.0 | 0.0 |
| SumDBs = 1 -- SumLength = 32 | -0.0 | 0.0 |
| LipidClass = PE -- SumDBs = 3 | 0.0 | 0.0 |
| LipidClass = SM -- SumLength = 38 | 0.0 | 0.0 |
| Headgroup = HEX2 -- LipidClass = Hex2Cer | -0.0 | 0.0 |
| LipidClass = PC O- -- SumDBs = 5 | 0.0 | 0.0 |
| LipidClass = PE -- SumLength = 34 | 0.0 | 0.0 |
| LipidClass = SM -- SumDBs = 2 | 0.0 | 0.0 |
| Headgroup = HEX2 -- SumLength = 42 | -0.0 | 0.0 |
| LipidClass = Hex2Cer -- SumLength = 42 | -0.0 | 0.0 |
| LipidClass = PS -- SumLength = 40 | 0.0 | 0.0 |
| LipidClass = PS -- SumDBs = 6 | 0.0 | 0.0 |
| Headgroup = PS -- SumLength = 40 | 0.0 | 0.0 |
| Headgroup = PS -- SumDBs = 6 | 0.0 | 0.0 |
| LipidClass = DG -- SumLength = 32 | -0.0 | 0.0 |
| LipidClass = PC O- -- SumDBs = 6 | 0.0 | 0.0 |
| SumDBs = 10 -- SumLength = 60 | 0.0 | 0.0 |
| SumDBs = 0 -- SumLength = 32 | -0.0 | 0.0 |
| LipidClass = DG -- SumDBs = 3 | -0.0 | 0.0 |
| LipidClass = PC -- SumDBs = 6 | 0.0 | 0.0 |
| SumDBs = 2 -- SumLength = 44 | -0.0 | 0.0 |
| Headgroup = HEX2 -- SumDBs = 2 | -0.0 | 0.0 |
| LipidClass = Hex2Cer -- SumDBs = 2 | -0.0 | 0.0 |
| Headgroup = PI -- LipidClass = PI | 0.0 | 0.0 |
| Headgroup = PC -- LipidClass = LPC O- | -0.0 | 0.0 |
| LipidClass = Cer -- SumLength = 34 | -0.0 | 0.0 |
| Headgroup = PI -- SumDBs = 4 | 0.0 | 0.0 |
| LipidClass = PI -- SumDBs = 4 | 0.0 | 0.0 |
| Headgroup = PE -- SumLength = 39 | 0.0 | 0.0 |
| LipidClass = PE O- -- SumLength = 39 | 0.0 | 0.0 |
| SumDBs = 4 -- SumLength = 44 | 0.0 | 0.0 |
| LipidClass = PC -- SumLength = 32 | -0.0 | 0.0 |
| LipidClass = Cer -- SumLength = 42 | -0.0 | 0.0 |
| SumDBs = 7 -- SumLength = 40 | 0.0 | 0.0 |
| LipidClass = PI -- SumLength = 38 | 0.0 | 0.0 |
| Headgroup = PI -- SumLength = 38 | 0.0 | 0.0 |
| Headgroup = PE -- SumDBs = 7 | 0.0 | 0.0 |
| Headgroup = PC -- SumLength = 41 | 0.0 | 0.0 |
| LipidClass = SM -- SumLength = 41 | 0.0 | 0.0 |
| SumDBs = 4 -- SumLength = 37 | 0.0 | 0.0 |
| SumDBs = 3 -- SumLength = 42 | -0.0 | 0.0 |
| Headgroup = -- SumDBs = 12 | -0.0 | 0.0 |
| SumDBs = 12 -- SumLength = 60 | -0.0 | 0.0 |
| LipidClass = TG -- SumDBs = 12 | -0.0 | 0.0 |
| SumDBs = 4 -- SumLength = 39 | 0.0 | 0.0 |
| LipidClass = PC -- SumDBs = 0 | 0.0 | 0.0 |
| LipidClass = PC -- SumLength = 40 | 0.0 | 0.0 |
| LipidClass = PC O- -- SumLength = 32 | 0.0 | 0.0 |
| Headgroup = HEX3 -- LipidClass = Hex3Cer | 0.0 | 0.0 |
| LipidClass = LPC O- -- SumDBs = 0 | -0.0 | 0.0 |
| LipidClass = PE -- SumLength = 32 | 0.0 | 0.0 |
| SumDBs = 2 -- SumLength = 32 | -0.0 | 0.0 |
| LipidClass = PE -- SumDBs = 7 | 0.0 | 0.0 |
| LipidClass = LPC O- -- SumLength = 16 | -0.0 | 0.0 |
| LipidClass = Cer -- SumDBs = 2 | 0.0 | 0.0 |
| LipidClass = Hex3Cer -- SumDBs = 1 | 0.0 | 0.0 |
| Headgroup = HEX3 -- SumDBs = 1 | 0.0 | 0.0 |
| Headgroup = PC -- SumLength = 33 | 0.0 | 0.0 |
| Headgroup = HEX -- LipidClass = HexCer | -0.0 | 0.0 |
| LipidClass = SM -- SumLength = 33 | 0.0 | 0.0 |
| LipidClass = PE -- SumDBs = 2 | 0.0 | 0.0 |
| LipidClass = PE O- -- SumLength = 32 | -0.0 | 0.0 |
| LipidClass = LPC O- -- SumLength = 18 | -0.0 | 0.0 |
| LipidClass = Cer -- SumLength = 33 | 0.0 | 0.0 |
| LipidClass = Cer -- SumDBs = 3 | -0.0 | 0.0 |
| Headgroup = HEX -- SumDBs = 1 | -0.0 | 0.0 |
| LipidClass = HexCer -- SumDBs = 1 | -0.0 | 0.0 |
| SumDBs = 1 -- SumLength = 20 | -0.0 | 0.0 |
| LipidClass = LPC O- -- SumDBs = 1 | -0.0 | 0.0 |
| LipidClass = Hex2Cer -- SumDBs = 1 | -0.0 | 0.0 |
| Headgroup = HEX2 -- SumDBs = 1 | -0.0 | 0.0 |
| LipidClass = LPE -- SumDBs = 4 | 0.0 | 0.0 |
| LipidClass = SM -- SumLength = 32 | 0.0 | 0.0 |
| SumDBs = 11 -- SumLength = 60 | -0.0 | 0.0 |
| Headgroup = -- SumDBs = 11 | -0.0 | 0.0 |
| LipidClass = TG -- SumDBs = 11 | -0.0 | 0.0 |
| Headgroup = PC -- SumLength = 35 | 0.0 | 0.0 |
| LipidClass = SM -- SumDBs = 0 | -0.0 | 0.0 |
| SumDBs = 2 -- SumLength = 41 | 0.0 | 0.0 |
| LipidClass = PC O- -- SumDBs = 0 | 0.0 | 0.0 |
| Headgroup = HEX -- SumLength = 34 | -0.0 | 0.0 |
| LipidClass = HexCer -- SumLength = 34 | -0.0 | 0.0 |
| LipidClass = DG -- SumLength = 30 | -0.0 | 0.0 |
| Headgroup = -- SumLength = 30 | -0.0 | 0.0 |
| SumDBs = 4 -- SumLength = 57 | 0.0 | 0.0 |
| LipidClass = Cer -- SumLength = 40 | 0.0 | 0.0 |
| LipidClass = DG -- SumDBs = 0 | -0.0 | 0.0 |
| SumDBs = 3 -- SumLength = 34 | 0.0 | 0.0 |
| LipidClass = PE O- -- SumLength = 35 | -0.0 | 0.0 |
| SumDBs = 1 -- SumLength = 16 | -0.0 | 0.0 |
| LipidClass = LPC -- SumDBs = 2 | 0.0 | 0.0 |
| Headgroup = PG -- SumDBs = 2 | -0.0 | 0.0 |
| LipidClass = PG -- SumDBs = 2 | -0.0 | 0.0 |
| Headgroup = PG -- SumLength = 36 | -0.0 | 0.0 |
| LipidClass = PG -- SumLength = 36 | -0.0 | 0.0 |
| Headgroup = PE -- SumLength = 35 | -0.0 | 0.0 |
| LipidClass = PE O- -- SumDBs = 7 | 0.0 | 0.0 |
| SumDBs = 5 -- SumLength = 39 | 0.0 | 0.0 |
| SumDBs = 0 -- SumLength = 17 | -0.0 | 0.0 |
| Headgroup = PG -- LipidClass = PG | -0.0 | 0.0 |
| LipidClass = LPE -- SumDBs = 2 | -0.0 | 0.0 |
| LipidClass = PE O- -- SumLength = 33 | -0.0 | 0.0 |
| Headgroup = PE -- SumLength = 33 | -0.0 | 0.0 |
| SumDBs = 4 -- SumLength = 40 | 0.0 | 0.0 |
| LipidClass = PE O- -- SumDBs = 0 | -0.0 | 0.0 |
| LipidClass = LPC -- SumDBs = 4 | 0.0 | 0.0 |
| SumDBs = 1 -- SumLength = 33 | 0.0 | 0.0 |
| LipidClass = PS -- SumDBs = 3 | 0.0 | 0.0 |
| Headgroup = PS -- SumDBs = 3 | 0.0 | 0.0 |
| SumDBs = 1 -- SumLength = 30 | -0.0 | 0.0 |
| LipidClass = Hex3Cer -- SumLength = 42 | 0.0 | 0.0 |
| Headgroup = HEX3 -- SumLength = 42 | 0.0 | 0.0 |
| LipidClass = PE -- SumDBs = 1 | 0.0 | 0.0 |
| LipidClass = Hex3Cer -- SumLength = 40 | 0.0 | 0.0 |
| Headgroup = HEX3 -- SumLength = 40 | 0.0 | 0.0 |
| LipidClass = PE O- -- SumLength = 42 | -0.0 | 0.0 |
| Headgroup = PE -- SumLength = 42 | -0.0 | 0.0 |
| LipidClass = LPC -- SumLength = 17 | -0.0 | 0.0 |
| Headgroup = PC -- SumLength = 17 | -0.0 | 0.0 |
| LipidClass = PC O- -- SumLength = 30 | 0.0 | 0.0 |
| LipidClass = SM -- SumLength = 43 | 0.0 | 0.0 |
| Headgroup = PC -- SumLength = 43 | 0.0 | 0.0 |
| LipidClass = PC O- -- SumLength = 35 | 0.0 | 0.0 |
| Headgroup = PC -- SumLength = 39 | 0.0 | 0.0 |
| LipidClass = SM -- SumLength = 39 | 0.0 | 0.0 |
| LipidClass = Cer -- SumLength = 43 | 0.0 | 0.0 |
| SumDBs = 1 -- SumLength = 39 | 0.0 | 0.0 |
| Headgroup = PI -- SumDBs = 5 | -0.0 | 0.0 |
| LipidClass = PI -- SumDBs = 5 | -0.0 | 0.0 |
| Headgroup = PE -- SumLength = 32 | 0.0 | 0.0 |
| SumDBs = 2 -- SumLength = 35 | 0.0 | 0.0 |
| Headgroup = PC -- SumLength = 42 | -0.0 | 0.0 |
| LipidClass = SM -- SumLength = 42 | -0.0 | 0.0 |
| LipidClass = Hex3Cer -- SumLength = 34 | 0.0 | 0.0 |
| Headgroup = HEX3 -- SumLength = 34 | 0.0 | 0.0 |
| Headgroup = PI -- SumDBs = 2 | 0.0 | 0.0 |
| LipidClass = PI -- SumDBs = 2 | 0.0 | 0.0 |
| LipidClass = PE -- SumLength = 37 | 0.0 | 0.0 |
| LipidClass = PE -- SumDBs = 0 | 0.0 | 0.0 |
| LipidClass = PC -- SumLength = 35 | 0.0 | 0.0 |
| LipidClass = Cer -- SumDBs = 0 | -0.0 | 0.0 |
| SumDBs = 2 -- SumLength = 20 | -0.0 | 0.0 |
| Headgroup = PI -- SumLength = 36 | 0.0 | 0.0 |
| LipidClass = PI -- SumLength = 36 | 0.0 | 0.0 |
| Headgroup = PC -- SumLength = 30 | 0.0 | 0.0 |
| LipidClass = DG -- SumDBs = 4 | -0.0 | 0.0 |
| SumDBs = 5 -- SumLength = 37 | 0.0 | 0.0 |
| SumDBs = 7 -- SumLength = 60 | -0.0 | 0.0 |
| Headgroup = HEX -- SumLength = 42 | -0.0 | 0.0 |
| LipidClass = HexCer -- SumLength = 42 | -0.0 | 0.0 |
| Headgroup = -- SumLength = 33 | -0.0 | 0.0 |
| Headgroup = PC -- SumDBs = 7 | 0.0 | 0.0 |
| SumDBs = 0 -- SumLength = 33 | -0.0 | 0.0 |
| LipidClass = PC -- SumDBs = 7 | 0.0 | 0.0 |
| LipidClass = SM -- SumLength = 35 | 0.0 | 0.0 |
| SumDBs = 2 -- SumLength = 43 | 0.0 | 0.0 |
| LipidClass = Cer -- SumLength = 36 | -0.0 | 0.0 |
| SumDBs = 2 -- SumLength = 33 | 0.0 | 0.0 |
| Headgroup = PE -- SumLength = 17 | -0.0 | 0.0 |
| LipidClass = LPE -- SumLength = 17 | -0.0 | 0.0 |
| LipidClass = SM -- SumDBs = 3 | -0.0 | 0.0 |
| Headgroup = HEX2 -- SumDBs = 3 | -0.0 | 0.0 |
| LipidClass = Hex2Cer -- SumDBs = 3 | -0.0 | 0.0 |
| LipidClass = HexCer -- SumDBs = 2 | -0.0 | 0.0 |
| Headgroup = HEX -- SumDBs = 2 | -0.0 | 0.0 |
| LipidClass = Hex3Cer -- SumDBs = 2 | 0.0 | 0.0 |
| Headgroup = HEX3 -- SumDBs = 2 | 0.0 | 0.0 |
| Headgroup = PC -- SumLength = 32 | 0.0 | 0.0 |
| SumDBs = 5 -- SumLength = 42 | -0.0 | 0.0 |
| Headgroup = PC -- SumLength = 37 | 0.0 | 0.0 |
| SumDBs = 3 -- SumLength = 32 | -0.0 | 0.0 |
| SumDBs = 1 -- SumLength = 35 | -0.0 | 0.0 |
| SumDBs = 2 -- SumLength = 30 | -0.0 | 0.0 |
| Headgroup = HEX3 -- SumLength = 41 | 0.0 | 0.0 |
| LipidClass = Hex3Cer -- SumLength = 41 | 0.0 | 0.0 |
| LipidClass = PC -- SumDBs = 8 | 0.0 | 0.0 |
| Headgroup = PC -- SumDBs = 8 | 0.0 | 0.0 |
| LipidClass = LPC -- SumLength = 20 | 0.0 | 0.0 |
| SumDBs = 8 -- SumLength = 40 | 0.0 | 0.0 |
| LipidClass = DG -- SumLength = 38 | -0.0 | 0.0 |
| LipidClass = PC -- SumLength = 37 | 0.0 | 0.0 |
| LipidClass = DG -- SumLength = 28 | -0.0 | 0.0 |
| LipidClass = Cer -- SumLength = 41 | -0.0 | 0.0 |
| LipidClass = PC -- SumLength = 30 | -0.0 | 0.0 |
| Headgroup = PC -- SumLength = 31 | 0.0 | 0.0 |
| LipidClass = PI -- SumDBs = 3 | 0.0 | 0.0 |
| Headgroup = PI -- SumDBs = 3 | 0.0 | 0.0 |
| SumDBs = 0 -- SumLength = 30 | 0.0 | 0.0 |
| LipidClass = SM -- SumLength = 44 | 0.0 | 0.0 |
| Headgroup = PC -- SumLength = 44 | 0.0 | 0.0 |
| LipidClass = PC -- SumLength = 31 | 0.0 | 0.0 |
| Headgroup = -- SumLength = 35 | -0.0 | 0.0 |
| Headgroup = -- SumLength = 31 | -0.0 | 0.0 |
| LipidClass = DG -- SumDBs = 5 | -0.0 | 0.0 |
| Headgroup = PC -- SumLength = 14 | -0.0 | 0.0 |
| SumDBs = 0 -- SumLength = 14 | -0.0 | 0.0 |
| LipidClass = LPC -- SumLength = 14 | -0.0 | 0.0 |
| LipidClass = SM -- SumLength = 36 | -0.0 | 0.0 |
| SumDBs = 3 -- SumLength = 41 | 0.0 | 0.0 |
| LipidClass = DG -- SumLength = 31 | -0.0 | 0.0 |
| Headgroup = PC -- SumLength = 20 | 0.0 | 0.0 |
| SumDBs = 6 -- SumLength = 57 | -0.0 | 0.0 |
| LipidClass = HexCer -- SumLength = 40 | -0.0 | 0.0 |
| Headgroup = HEX -- SumLength = 40 | -0.0 | 0.0 |
| Headgroup = HEX2 -- SumLength = 41 | -0.0 | 0.0 |
| LipidClass = Hex2Cer -- SumLength = 41 | -0.0 | 0.0 |
| SumDBs = 1 -- SumLength = 28 | -0.0 | 0.0 |
| Headgroup = PG -- SumLength = 34 | 0.0 | 0.0 |
| LipidClass = PG -- SumLength = 34 | 0.0 | 0.0 |
| LipidClass = Cer -- SumLength = 32 | -0.0 | 0.0 |
| LipidClass = Hex3Cer -- SumLength = 38 | 0.0 | 0.0 |
| Headgroup = HEX3 -- SumLength = 38 | 0.0 | 0.0 |
| LipidClass = DG -- SumLength = 35 | -0.0 | 0.0 |
| Headgroup = PG -- SumDBs = 1 | 0.0 | 0.0 |
| LipidClass = PG -- SumDBs = 1 | 0.0 | 0.0 |
| LipidClass = PC -- SumLength = 33 | 0.0 | 0.0 |
| SumDBs = 3 -- SumLength = 43 | 0.0 | 0.0 |
| LipidClass = Hex2Cer -- SumLength = 38 | -0.0 | 0.0 |
| Headgroup = HEX2 -- SumLength = 38 | -0.0 | 0.0 |
| Headgroup = HEX2 -- SumLength = 36 | -0.0 | 0.0 |
| LipidClass = Hex2Cer -- SumLength = 36 | -0.0 | 0.0 |
| LipidClass = PE -- SumLength = 35 | 0.0 | 0.0 |
| LipidClass = LPC -- SumLength = 19 | -0.0 | 0.0 |
| Headgroup = PC -- SumLength = 19 | -0.0 | 0.0 |
| LipidClass = LPC -- SumDBs = 6 | 0.0 | 0.0 |
| Headgroup = PI -- SumLength = 37 | 0.0 | 0.0 |
| LipidClass = PI -- SumLength = 37 | 0.0 | 0.0 |
| SumDBs = 0 -- SumLength = 15 | 0.0 | 0.0 |
| Headgroup = PC -- SumLength = 15 | 0.0 | 0.0 |
| LipidClass = LPC -- SumLength = 15 | 0.0 | 0.0 |
| LipidClass = DG -- SumDBs = 6 | -0.0 | 0.0 |
| LipidClass = DG -- SumLength = 33 | -0.0 | 0.0 |
| LipidClass = LPC -- SumLength = 22 | 0.0 | 0.0 |
| Headgroup = PC -- SumLength = 22 | 0.0 | 0.0 |
| SumDBs = 4 -- SumLength = 42 | -0.0 | 0.0 |
| Headgroup = PG -- SumDBs = 5 | 0.0 | 0.0 |
| Headgroup = PG -- SumLength = 38 | 0.0 | 0.0 |
| LipidClass = PG -- SumDBs = 5 | 0.0 | 0.0 |
| LipidClass = PG -- SumLength = 38 | 0.0 | 0.0 |
| LipidClass = Cer -- SumDBs = 4 | -0.0 | 0.0 |
| Headgroup = PI -- SumLength = 34 | 0.0 | 0.0 |
| LipidClass = PI -- SumLength = 34 | 0.0 | 0.0 |
| SumDBs = 0 -- SumLength = 20 | -0.0 | 0.0 |
| LipidClass = LPC O- -- SumLength = 20 | -0.0 | 0.0 |
| SumDBs = 5 -- SumLength = 20 | 0.0 | 0.0 |
| SumDBs = 3 -- SumLength = 35 | -0.0 | 0.0 |
| SumDBs = 1 -- SumLength = 17 | -0.0 | 0.0 |
| SumDBs = 0 -- SumLength = 19 | -0.0 | 0.0 |
| SumDBs = 1 -- SumLength = 19 | -0.0 | 0.0 |
| SumDBs = 4 -- SumLength = 34 | -0.0 | 0.0 |
| LipidClass = PC O- -- SumLength = 37 | 0.0 | 0.0 |
| LipidClass = LPC -- SumDBs = 5 | 0.0 | 0.0 |
| LipidClass = PC -- SumLength = 29 | 0.0 | 0.0 |
| Headgroup = PC -- SumLength = 29 | 0.0 | 0.0 |
| SumDBs = 0 -- SumLength = 29 | 0.0 | 0.0 |
| Headgroup = PE -- SumLength = 22 | -0.0 | 0.0 |
| LipidClass = LPE -- SumLength = 22 | -0.0 | 0.0 |
| SumDBs = 0 -- SumLength = 22 | -0.0 | 0.0 |
| LipidClass = Cer -- SumLength = 35 | -0.0 | 0.0 |
| LipidClass = Cer -- SumLength = 39 | -0.0 | 0.0 |
| LipidClass = LPC -- SumDBs = 3 | -0.0 | 0.0 |
| LipidClass = DG -- SumLength = 40 | -0.0 | 0.0 |
| Headgroup = PG -- SumDBs = 4 | 0.0 | 0.0 |
| LipidClass = PG -- SumDBs = 4 | 0.0 | 0.0 |
| Headgroup = PE -- SumLength = 20 | 0.0 | 0.0 |
| LipidClass = LPE -- SumLength = 20 | 0.0 | 0.0 |
| LipidClass = Cer -- SumLength = 37 | 0.0 | 0.0 |
| LipidClass = Hex2Cer -- SumLength = 34 | -0.0 | 0.0 |
| Headgroup = HEX2 -- SumLength = 34 | -0.0 | 0.0 |
| SumDBs = 3 -- SumLength = 40 | -0.0 | 0.0 |
| LipidClass = PC -- SumLength = 28 | -0.0 | 0.0 |
| Headgroup = PC -- SumLength = 28 | -0.0 | 0.0 |
| Headgroup = HEX2 -- SumLength = 40 | -0.0 | 0.0 |
| LipidClass = Hex2Cer -- SumLength = 40 | -0.0 | 0.0 |
| SumDBs = 3 -- SumLength = 30 | -0.0 | 0.0 |
| SumDBs = 0 -- SumLength = 35 | -0.0 | 0.0 |
| Headgroup = PI -- SumDBs = 1 | 0.0 | 0.0 |
| LipidClass = PI -- SumDBs = 1 | 0.0 | 0.0 |
| Headgroup = PI -- SumDBs = 6 | -0.0 | 0.0 |
| LipidClass = PI -- SumDBs = 6 | -0.0 | 0.0 |
| SumDBs = 6 -- SumLength = 36 | -0.0 | 0.0 |
| Headgroup = -- SumLength = 37 | 0.0 | 0.0 |
| LipidClass = LPE -- SumDBs = 3 | 0.0 | 0.0 |
| Headgroup = PC -- SumLength = 24 | -0.0 | 0.0 |
| LipidClass = LPE -- SumDBs = 5 | 0.0 | 0.0 |
| SumDBs = 1 -- SumLength = 37 | 0.0 | 0.0 |
| SumDBs = 1 -- SumLength = 22 | -0.0 | 0.0 |
| LipidClass = DG -- SumLength = 37 | -0.0 | 0.0 |
| LipidClass = LPC O- -- SumLength = 24 | -0.0 | 0.0 |
| LipidClass = DG -- SumDBs = 7 | -0.0 | 0.0 |
| LipidClass = Cer -- SumLength = 31 | 0.0 | 0.0 |
| Headgroup = HEX -- SumLength = 41 | -0.0 | 0.0 |
| LipidClass = HexCer -- SumLength = 41 | -0.0 | 0.0 |
| LipidClass = PI -- SumLength = 32 | -0.0 | 0.0 |
| LipidClass = PI -- SumDBs = 0 | -0.0 | 0.0 |
| Headgroup = PI -- SumLength = 32 | -0.0 | 0.0 |
| Headgroup = PI -- SumDBs = 0 | -0.0 | 0.0 |
| SumDBs = 2 -- SumLength = 39 | -0.0 | 0.0 |
| SumDBs = 0 -- SumLength = 38 | 0.0 | 0.0 |
| SumDBs = 2 -- SumLength = 28 | -0.0 | 0.0 |
| LipidClass = PI -- SumLength = 40 | -0.0 | 0.0 |
| Headgroup = PI -- SumLength = 40 | -0.0 | 0.0 |
| SumDBs = 1 -- SumLength = 31 | 0.0 | 0.0 |
| LipidClass = SM -- SumLength = 31 | 0.0 | 0.0 |
| LipidClass = LPC -- SumLength = 24 | -0.0 | 0.0 |
| LipidClass = DG -- SumLength = 26 | -0.0 | 0.0 |
| LipidClass = DG -- SumDBs = 8 | -0.0 | 0.0 |
| LipidClass = Cer -- SumLength = 44 | -0.0 | 0.0 |
| LipidClass = PC O- -- SumDBs = 7 | 0.0 | 0.0 |
| SumDBs = 0 -- SumLength = 31 | 0.0 | 0.0 |
| SumDBs = 7 -- SumLength = 42 | 0.0 | 0.0 |
| LipidClass = SM -- SumLength = 37 | -0.0 | 0.0 |
| SumDBs = 2 -- SumLength = 37 | 0.0 | 0.0 |
| SumDBs = 7 -- SumLength = 38 | -0.0 | 0.0 |
| LipidClass = Cer -- SumLength = 46 | -0.0 | 0.0 |
| LipidClass = Cer -- SumLength = 30 | 0.0 | 0.0 |

  


##### Best Substructure Features
