## Supplementary material for "Lipid network and moiety analysis for revealing enzymatic dysregulation and mechanistic alterations from lipidomics data": LINEX_network_MBOAT7.html

LINEX - LipidNetworkExplorer


- Visit the LINEX website

Network

Network Enrichment

Lipidome Summary

Version 2

##### Network Options

**Network Type**

Native Network
Enzyme Network
Native Molecular Network
Enzyme Molecular Network

Recompute Network
  


---


**Node Colours**

Lipid Class
Desaturation
Chain Length
C Index
DB Index
-log10(FDR)
Fold Changes
OH Index
Betweenness Centrality
Closeness Centrality
Degree
Hydroxylation

**Edge Colours**

Reaction Type
Correlations
Correlation Changes

**Node Sizes**

-log10(FDR)
Fold Changes
DB Index
C Index
OH Index
Betweenness Centrality
Closeness Centrality
Degree
Hydroxylation
Desaturation
Chain Length


---


**Comparison**

WT HFCD\_KO HFCD
WT HFCD\_WT CHOW
WT HFCD\_KO CHOW
KO HFCD\_WT CHOW
KO HFCD\_KO CHOW
WT CHOW\_KO CHOW

**Group**

WT HFCD
KO HFCD
WT CHOW
KO CHOW


---


**Find Lipid Species**


Find
  

**Find by Substring**

Find


---


  
  
**Shown reaction types**

all
only Class reactions
only FA reactions

  
  
**Significance Threshold**

Update
  

**Highlight**

**Enable physics**

**Unified Scales**

---


Download View

Downloads the network as a .json file, which can be uploaded to 127.0.0.1:8000/download to generate a pdf view

##### Legend

Hide Legend Navigation

##### Network

0%


Enrichment settings not available in downloaded file

Lipidome Overview


Substructure Analysis


Chain Length Analysis

##### Substructure Features

##### Regression-Based Feature Selection

| KO CHOW | | | KO HFCD | | | WT CHOW | | | WT HFCD | | |
| --- | --- | --- | --- | --- | --- | --- | --- | --- | --- | --- | --- |
| **Feature** | **Coefficient** | **Absolute Coefficient** | **Feature** | **Coefficient** | **Absolute Coefficient** | **Feature** | **Coefficient** | **Absolute Coefficient** | **Feature** | **Coefficient** | **Absolute Coefficient** |
| SumDBs = 7 -- SumLength = 40 | -0.004 | 0.004 | Headgroup = PI -- SumDBs = 2 | -0.008 | 0.008 | Headgroup = -- SumLength = 40 | -0.005 | 0.005 | Headgroup = PI -- SumDBs = 1 | -0.007 | 0.007 |
| Headgroup = PI -- SumLength = 38 | -0.004 | 0.004 | LipidClass = PI -- SumDBs = 2 | -0.008 | 0.008 | LipidClass = Cer -- SumLength = 40 | -0.005 | 0.005 | LipidClass = LPI -- SumDBs = 1 | -0.007 | 0.007 |
| LipidClass = PI -- SumLength = 38 | -0.004 | 0.004 | SumDBs = 1 -- SumLength = 40 | -0.007 | 0.007 | LipidClass = PI -- SumDBs = 2 | -0.005 | 0.005 | SumDBs = 2 -- SumLength = 41 | -0.006 | 0.006 |
| LipidClass = LPI -- SumDBs = 1 | -0.004 | 0.004 | SumDBs = 4 -- SumLength = 38 | -0.007 | 0.007 | Headgroup = PI -- SumDBs = 2 | -0.005 | 0.005 | SumDBs = 7 -- SumLength = 40 | -0.005 | 0.005 |
| Headgroup = PI -- SumDBs = 1 | -0.004 | 0.004 | LipidClass = PE -- SumDBs = 4 | -0.006 | 0.006 | SumDBs = 1 -- SumLength = 40 | -0.005 | 0.005 | Headgroup = PI -- LipidClass = LPI | -0.005 | 0.005 |
| Headgroup = PE -- SumDBs = 7 | -0.003 | 0.003 | SumDBs = 6 -- SumLength = 40 | -0.006 | 0.006 | SumDBs = 4 -- SumLength = 38 | -0.004 | 0.004 | LipidClass = PC -- SumLength = 34 | -0.004 | 0.004 |
| LipidClass = PI -- SumDBs = 4 | -0.003 | 0.003 | LipidClass = PC O- -- SumDBs = 4 | -0.006 | 0.006 | Headgroup = PE -- SumLength = 38 | -0.004 | 0.004 | Headgroup = -- SumDBs = 4 | -0.004 | 0.004 |
| SumDBs = 2 -- SumLength = 41 | -0.003 | 0.003 | SumDBs = 1 -- SumLength = 41 | -0.005 | 0.005 | Headgroup = -- SumDBs = 1 | -0.004 | 0.004 | Headgroup = PI -- SumDBs = 5 | -0.004 | 0.004 |
| LipidClass = TG -- SumDBs = 4 | -0.003 | 0.003 | Headgroup = PE -- SumLength = 38 | -0.005 | 0.005 | Headgroup = PE -- SumDBs = 4 | -0.004 | 0.004 | LipidClass = PI -- SumDBs = 5 | -0.004 | 0.004 |
| Headgroup = PC -- SumDBs = 7 | -0.003 | 0.003 | Headgroup = -- SumLength = 40 | -0.005 | 0.005 | SumDBs = 2 -- SumLength = 52 | -0.003 | 0.003 | Headgroup = PI -- SumLength = 16 | -0.004 | 0.004 |
| LipidClass = PC O- -- SumDBs = 7 | -0.003 | 0.003 | LipidClass = Cer -- SumLength = 40 | -0.005 | 0.005 | LipidClass = PI -- SumLength = 36 | -0.003 | 0.003 | LipidClass = LPI -- SumLength = 16 | -0.004 | 0.004 |
| Headgroup = -- SumDBs = 4 | -0.003 | 0.003 | LipidClass = PI -- SumLength = 36 | -0.005 | 0.005 | Headgroup = PI -- SumLength = 36 | -0.003 | 0.003 | LipidClass = PI -- SumLength = 38 | -0.004 | 0.004 |
| Headgroup = PE -- SumLength = 40 | -0.002 | 0.002 | Headgroup = PI -- SumLength = 36 | -0.005 | 0.005 | SumDBs = 5 -- SumLength = 38 | -0.003 | 0.003 | Headgroup = PI -- SumLength = 38 | -0.004 | 0.004 |
| Headgroup = P -- SumLength = 36 | -0.002 | 0.002 | Headgroup = P -- SumDBs = 4 | -0.005 | 0.005 | LipidClass = LPC -- SumDBs = 2 | -0.003 | 0.003 | LipidClass = PI -- SumLength = 40 | -0.004 | 0.004 |
| LipidClass = PA -- SumLength = 36 | -0.002 | 0.002 | LipidClass = PA -- SumDBs = 4 | -0.005 | 0.005 | Headgroup = PC -- SumDBs = 1 | -0.003 | 0.003 | Headgroup = PI -- SumLength = 40 | -0.004 | 0.004 |
| Headgroup = PE -- SumDBs = 6 | -0.002 | 0.002 | Headgroup = PG -- SumDBs = 5 | -0.005 | 0.005 | SumDBs = 2 -- SumLength = 36 | -0.003 | 0.003 | LipidClass = PC -- SumDBs = 4 | -0.004 | 0.004 |
| LipidClass = PE O- -- SumDBs = 7 | -0.002 | 0.002 | LipidClass = PG -- SumDBs = 5 | -0.005 | 0.005 | LipidClass = LPG -- SumDBs = 1 | -0.002 | 0.002 | Headgroup = PE -- SumLength = 34 | -0.004 | 0.004 |
| Headgroup = PC -- SumLength = 34 | -0.002 | 0.002 | Headgroup = PE -- SumDBs = 4 | -0.005 | 0.005 | LipidClass = PG -- SumLength = 38 | -0.002 | 0.002 | LipidClass = PI -- SumDBs = 6 | -0.004 | 0.004 |
| LipidClass = DG -- SumDBs = 3 | -0.002 | 0.002 | SumDBs = 5 -- SumLength = 38 | -0.004 | 0.004 | Headgroup = PG -- SumLength = 38 | -0.002 | 0.002 | Headgroup = PI -- SumDBs = 6 | -0.004 | 0.004 |
| Headgroup = CE -- SumDBs = 0 | -0.002 | 0.002 | Headgroup = PC -- SumDBs = 6 | -0.004 | 0.004 | Headgroup = P -- SumDBs = 4 | -0.002 | 0.002 | Headgroup = PC -- SumDBs = 7 | -0.004 | 0.004 |
| LipidClass = CE -- SumDBs = 0 | -0.002 | 0.002 | LipidClass = LPC -- SumDBs = 0 | -0.004 | 0.004 | LipidClass = PA -- SumDBs = 4 | -0.002 | 0.002 | LipidClass = PC O- -- SumDBs = 3 | -0.004 | 0.004 |
| SumDBs = 0 -- SumLength = 32 | -0.002 | 0.002 | Headgroup = PS -- SumDBs = 6 | -0.004 | 0.004 | LipidClass = PG -- SumDBs = 2 | -0.002 | 0.002 | SumDBs = 1 -- SumLength = 18 | -0.004 | 0.004 |
| SumDBs = 7 -- SumLength = 38 | -0.002 | 0.002 | LipidClass = PS -- SumDBs = 6 | -0.004 | 0.004 | SumDBs = 1 -- SumLength = 41 | -0.002 | 0.002 | Headgroup = PG -- SumLength = 34 | -0.004 | 0.004 |
| SumDBs = 4 -- SumLength = 52 | -0.002 | 0.002 | Headgroup = PC -- SumLength = 40 | -0.004 | 0.004 | LipidClass = PC O- -- SumDBs = 4 | -0.002 | 0.002 | LipidClass = PG -- SumLength = 34 | -0.004 | 0.004 |
| LipidClass = SM -- SumLength = 34 | -0.002 | 0.002 | Headgroup = PI -- SumLength = 34 | -0.004 | 0.004 | LipidClass = PE -- SumLength = 38 | -0.002 | 0.002 | SumDBs = 3 -- SumLength = 34 | -0.003 | 0.003 |
| SumDBs = 3 -- SumLength = 42 | -0.002 | 0.002 | LipidClass = PI -- SumLength = 34 | -0.004 | 0.004 | LipidClass = SM -- SumDBs = 1 | -0.002 | 0.002 | LipidClass = Cer -- SumDBs = 2 | -0.003 | 0.003 |
| Headgroup = PI -- SumLength = 18 | -0.002 | 0.002 | LipidClass = PG -- SumLength = 38 | -0.003 | 0.003 | LipidClass = SM -- SumLength = 34 | -0.002 | 0.002 | SumDBs = 5 -- SumLength = 40 | -0.003 | 0.003 |
| LipidClass = LPI -- SumLength = 18 | -0.002 | 0.002 | Headgroup = PG -- SumLength = 38 | -0.003 | 0.003 | SumDBs = 3 -- SumLength = 54 | -0.002 | 0.002 | LipidClass = TG -- SumDBs = 4 | -0.003 | 0.003 |
| LipidClass = PI -- SumLength = 40 | -0.002 | 0.002 | LipidClass = PC -- SumDBs = 6 | -0.003 | 0.003 | Headgroup = PC -- SumDBs = 5 | -0.002 | 0.002 | LipidClass = PC O- -- SumDBs = 7 | -0.003 | 0.003 |
| Headgroup = PI -- SumLength = 40 | -0.002 | 0.002 | Headgroup = PI -- SumLength = 20 | -0.003 | 0.003 | LipidClass = PE O- -- SumLength = 38 | -0.002 | 0.002 | SumDBs = 1 -- SumLength = 42 | -0.003 | 0.003 |
| LipidClass = PC -- SumDBs = 4 | -0.002 | 0.002 | LipidClass = LPI -- SumDBs = 4 | -0.003 | 0.003 | Headgroup = PC -- SumLength = 16 | -0.002 | 0.002 | LipidClass = CE -- SumLength = 16 | -0.003 | 0.003 |
| LipidClass = LPI -- SumLength = 16 | -0.002 | 0.002 | LipidClass = LPI -- SumLength = 20 | -0.003 | 0.003 | LipidClass = LPC -- SumLength = 16 | -0.002 | 0.002 | Headgroup = CE -- SumLength = 16 | -0.003 | 0.003 |
| Headgroup = PI -- SumLength = 16 | -0.002 | 0.002 | LipidClass = Cer -- SumDBs = 1 | -0.003 | 0.003 | Headgroup = PS -- SumDBs = 6 | -0.002 | 0.002 | Headgroup = PI -- SumLength = 18 | -0.003 | 0.003 |
| LipidClass = PA -- SumDBs = 1 | -0.002 | 0.002 | SumDBs = 2 -- SumLength = 34 | -0.003 | 0.003 | LipidClass = PS -- SumDBs = 6 | -0.002 | 0.002 | LipidClass = LPI -- SumLength = 18 | -0.003 | 0.003 |
| Headgroup = P -- SumDBs = 1 | -0.002 | 0.002 | SumDBs = 2 -- SumLength = 42 | -0.003 | 0.003 | LipidClass = DG -- SumDBs = 1 | -0.002 | 0.002 | SumDBs = 6 -- SumLength = 38 | -0.003 | 0.003 |
| Headgroup = PC -- SumDBs = 3 | -0.002 | 0.002 | LipidClass = TG -- SumDBs = 2 | -0.003 | 0.003 | Headgroup = P -- SumLength = 34 | -0.002 | 0.002 | SumDBs = 1 -- SumLength = 32 | -0.003 | 0.003 |
| LipidClass = PE -- SumDBs = 6 | -0.002 | 0.002 | Headgroup = PC -- SumDBs = 2 | -0.003 | 0.003 | LipidClass = PA -- SumLength = 34 | -0.002 | 0.002 | Headgroup = PC -- SumLength = 32 | -0.003 | 0.003 |
| LipidClass = PI -- SumDBs = 5 | -0.002 | 0.002 | LipidClass = PC O- -- SumDBs = 6 | -0.003 | 0.003 | LipidClass = PI -- SumLength = 34 | -0.002 | 0.002 | LipidClass = PS -- SumDBs = 4 | -0.003 | 0.003 |
| Headgroup = PI -- SumDBs = 5 | -0.002 | 0.002 | LipidClass = PC -- SumLength = 38 | -0.003 | 0.003 | Headgroup = PI -- SumLength = 34 | -0.002 | 0.002 | Headgroup = PS -- SumDBs = 4 | -0.003 | 0.003 |
| Headgroup = PI -- LipidClass = LPI | -0.002 | 0.002 | SumDBs = 2 -- SumLength = 52 | -0.003 | 0.003 | LipidClass = TG -- SumDBs = 2 | -0.002 | 0.002 | Headgroup = PE -- SumDBs = 6 | -0.003 | 0.003 |
| LipidClass = Cer -- SumLength = 38 | -0.002 | 0.002 | LipidClass = PA -- SumLength = 34 | -0.003 | 0.003 | LipidClass = PE -- SumDBs = 4 | -0.002 | 0.002 | LipidClass = PE -- SumLength = 34 | -0.003 | 0.003 |
| LipidClass = PE O- -- SumLength = 40 | -0.002 | 0.002 | Headgroup = P -- SumLength = 34 | -0.003 | 0.003 | LipidClass = PC O- -- SumDBs = 5 | -0.002 | 0.002 | SumDBs = 0 -- SumLength = 32 | -0.003 | 0.003 |
| LipidClass = PS -- SumDBs = 4 | -0.002 | 0.002 | SumDBs = 0 -- SumLength = 18 | -0.003 | 0.003 | LipidClass = PC -- SumLength = 36 | -0.002 | 0.002 | LipidClass = PA -- SumLength = 36 | -0.003 | 0.003 |
| Headgroup = PS -- SumDBs = 4 | -0.002 | 0.002 | Headgroup = PC -- SumLength = 16 | -0.003 | 0.003 | Headgroup = PE -- SumDBs = 5 | -0.002 | 0.002 | Headgroup = P -- SumLength = 36 | -0.003 | 0.003 |
| Headgroup = PI -- SumDBs = 6 | -0.002 | 0.002 | LipidClass = LPC -- SumLength = 16 | -0.003 | 0.003 | LipidClass = PE O- -- SumDBs = 5 | -0.001 | 0.001 | Headgroup = PS -- SumLength = 36 | -0.003 | 0.003 |
| LipidClass = PI -- SumDBs = 6 | -0.002 | 0.002 | LipidClass = PE O- -- SumLength = 38 | -0.003 | 0.003 | Headgroup = -- SumDBs = 2 | -0.001 | 0.001 | LipidClass = PS -- SumLength = 36 | -0.003 | 0.003 |
| LipidClass = PG -- SumDBs = 6 | -0.002 | 0.002 | LipidClass = TG -- SumDBs = 3 | -0.003 | 0.003 | SumDBs = 5 -- SumLength = 40 | -0.001 | 0.001 | Headgroup = PC -- SumLength = 34 | -0.003 | 0.003 |
| LipidClass = PC O- -- SumDBs = 0 | -0.002 | 0.002 | Headgroup = PC -- SumDBs = 1 | -0.003 | 0.003 | SumDBs = 2 -- SumLength = 18 | -0.001 | 0.001 | LipidClass = PE -- SumDBs = 6 | -0.003 | 0.003 |
| LipidClass = PC O- -- SumLength = 32 | -0.002 | 0.002 | LipidClass = LPE -- SumDBs = 1 | -0.003 | 0.003 | Headgroup = PG -- SumDBs = 4 | -0.001 | 0.001 | Headgroup = PE -- SumDBs = 2 | -0.002 | 0.002 |
| SumDBs = 0 -- SumLength = 16 | -0.002 | 0.002 | Headgroup = PE -- SumDBs = 5 | -0.003 | 0.003 | LipidClass = PC -- SumLength = 38 | -0.001 | 0.001 | Headgroup = PG -- SumDBs = 2 | -0.002 | 0.002 |
| Headgroup = PC -- SumLength = 32 | -0.001 | 0.001 | LipidClass = PE -- SumLength = 38 | -0.002 | 0.002 | LipidClass = TG -- SumDBs = 3 | -0.001 | 0.001 | SumDBs = 0 -- SumLength = 16 | -0.002 | 0.002 |
| LipidClass = PA -- SumDBs = 2 | -0.001 | 0.001 | SumDBs = 2 -- SumLength = 36 | -0.002 | 0.002 | LipidClass = PE O- -- SumDBs = 4 | -0.001 | 0.001 | SumDBs = 4 -- SumLength = 52 | -0.002 | 0.002 |
| Headgroup = P -- SumDBs = 2 | -0.001 | 0.001 | LipidClass = PC O- -- SumDBs = 2 | -0.002 | 0.002 | Headgroup = CE -- SumLength = 22 | -0.001 | 0.001 | LipidClass = Cer -- SumLength = 38 | -0.002 | 0.002 |
| Headgroup = PI -- SumDBs = 4 | -0.001 | 0.001 | Headgroup = PE -- SumDBs = 1 | -0.002 | 0.002 | Headgroup = CE -- SumDBs = 6 | -0.001 | 0.001 | LipidClass = PG -- SumDBs = 2 | -0.002 | 0.002 |
| LipidClass = PC O- -- SumLength = 38 | -0.001 | 0.001 | LipidClass = PE O- -- SumDBs = 5 | -0.002 | 0.002 | LipidClass = CE -- SumDBs = 6 | -0.001 | 0.001 | Headgroup = PC -- SumLength = 22 | -0.002 | 0.002 |
| LipidClass = PC O- -- SumDBs = 3 | -0.001 | 0.001 | Headgroup = PC -- SumLength = 38 | -0.002 | 0.002 | LipidClass = CE -- SumLength = 22 | -0.001 | 0.001 | LipidClass = LPC -- SumDBs = 6 | -0.002 | 0.002 |
| LipidClass = Cer -- SumDBs = 2 | -0.001 | 0.001 | Headgroup = -- SumDBs = 1 | -0.002 | 0.002 | Headgroup = PS -- SumLength = 38 | -0.001 | 0.001 | LipidClass = LPC -- SumLength = 22 | -0.002 | 0.002 |
| LipidClass = LPC -- SumDBs = 6 | -0.001 | 0.001 | LipidClass = PI -- SumDBs = 3 | -0.002 | 0.002 | LipidClass = PS -- SumLength = 38 | -0.001 | 0.001 | LipidClass = LPG -- SumDBs = 3 | -0.002 | 0.002 |
| Headgroup = PC -- SumLength = 22 | -0.001 | 0.001 | Headgroup = PI -- SumDBs = 3 | -0.002 | 0.002 | SumDBs = 0 -- SumLength = 18 | -0.001 | 0.001 | LipidClass = PG -- SumDBs = 1 | -0.002 | 0.002 |
| LipidClass = LPC -- SumLength = 22 | -0.001 | 0.001 | LipidClass = DG -- SumDBs = 6 | -0.002 | 0.002 | LipidClass = TG -- SumDBs = 1 | -0.001 | 0.001 | Headgroup = PE -- SumDBs = 7 | -0.002 | 0.002 |
| SumDBs = 3 -- SumLength = 36 | -0.001 | 0.001 | LipidClass = PC -- SumDBs = 3 | -0.002 | 0.002 | SumDBs = 4 -- SumLength = 20 | -0.001 | 0.001 | LipidClass = SM -- SumDBs = 2 | -0.002 | 0.002 |
| SumDBs = 5 -- SumLength = 36 | -0.001 | 0.001 | LipidClass = PC O- -- SumLength = 40 | -0.002 | 0.002 | LipidClass = PE O- -- SumDBs = 6 | -0.001 | 0.001 | Headgroup = -- SumLength = 42 | -0.002 | 0.002 |
| LipidClass = CE -- SumLength = 16 | -0.001 | 0.001 | SumDBs = 8 -- SumLength = 56 | -0.002 | 0.002 | SumDBs = 1 -- SumLength = 50 | -0.001 | 0.001 | LipidClass = Cer -- SumLength = 42 | -0.002 | 0.002 |
| Headgroup = -- SumLength = 38 | -0.001 | 0.001 | LipidClass = PC O- -- SumLength = 36 | -0.002 | 0.002 | LipidClass = Cer -- SumDBs = 1 | -0.001 | 0.001 | Headgroup = PC -- SumDBs = 3 | -0.002 | 0.002 |
| Headgroup = CE -- SumLength = 16 | -0.001 | 0.001 | Headgroup = PI -- SumDBs = 4 | -0.002 | 0.002 | LipidClass = LPG -- SumLength = 20 | -0.001 | 0.001 | SumDBs = 3 -- SumLength = 18 | -0.002 | 0.002 |
| LipidClass = TG -- SumDBs = 6 | -0.001 | 0.001 | Headgroup = -- SumDBs = 2 | -0.002 | 0.002 | LipidClass = LPG -- SumDBs = 4 | -0.001 | 0.001 | Headgroup = P -- SumDBs = 1 | -0.002 | 0.002 |
| SumDBs = 3 -- SumLength = 34 | -0.001 | 0.001 | Headgroup = CE -- SumLength = 18 | -0.002 | 0.002 | Headgroup = PG -- SumLength = 20 | -0.001 | 0.001 | LipidClass = PA -- SumDBs = 1 | -0.002 | 0.002 |
| Headgroup = -- SumDBs = 3 | -0.001 | 0.001 | LipidClass = CE -- SumLength = 18 | -0.002 | 0.002 | LipidClass = PC -- SumDBs = 3 | -0.001 | 0.001 | Headgroup = PC -- SumLength = 20 | -0.002 | 0.002 |
| Headgroup = PE -- SumLength = 36 | -0.001 | 0.001 | LipidClass = SM -- SumLength = 42 | -0.002 | 0.002 | Headgroup = -- SumLength = 54 | -0.001 | 0.001 | LipidClass = LPC -- SumLength = 20 | -0.002 | 0.002 |
| LipidClass = PC -- SumDBs = 2 | -0.001 | 0.001 | Headgroup = PC -- SumLength = 42 | -0.002 | 0.002 | LipidClass = TG -- SumLength = 54 | -0.001 | 0.001 | LipidClass = LPC -- SumDBs = 4 | -0.002 | 0.002 |
| LipidClass = PC -- SumLength = 34 | -0.001 | 0.001 | LipidClass = LPE -- SumLength = 16 | -0.002 | 0.002 | LipidClass = SM -- SumLength = 38 | -0.001 | 0.001 | SumDBs = 1 -- SumLength = 16 | -0.002 | 0.002 |
| LipidClass = PC -- SumLength = 36 | -0.001 | 0.001 | Headgroup = PE -- SumLength = 16 | -0.002 | 0.002 | LipidClass = DG -- SumDBs = 2 | -0.001 | 0.001 | LipidClass = PE O- -- SumDBs = 7 | -0.002 | 0.002 |
| LipidClass = PE -- SumDBs = 7 | -0.001 | 0.001 | LipidClass = PC O- -- SumDBs = 1 | -0.002 | 0.002 | Headgroup = PG -- SumDBs = 2 | -0.001 | 0.001 | Headgroup = P -- SumDBs = 2 | -0.002 | 0.002 |
| LipidClass = SM -- SumDBs = 3 | -0.001 | 0.001 | LipidClass = PC O- -- SumLength = 34 | -0.002 | 0.002 | Headgroup = -- SumLength = 60 | -0.001 | 0.001 | LipidClass = PA -- SumDBs = 2 | -0.002 | 0.002 |
| Headgroup = -- SumDBs = 5 | -0.001 | 0.001 | SumDBs = 3 -- SumLength = 52 | -0.001 | 0.001 | LipidClass = TG -- SumLength = 60 | -0.001 | 0.001 | LipidClass = TG -- SumDBs = 6 | -0.002 | 0.002 |
| LipidClass = Cer -- SumLength = 34 | -0.001 | 0.001 | Headgroup = PS -- SumDBs = 3 | -0.001 | 0.001 | LipidClass = LPE -- SumDBs = 0 | -0.001 | 0.001 | LipidClass = DG -- SumLength = 32 | -0.002 | 0.002 |
| LipidClass = TG -- SumDBs = 7 | -0.001 | 0.001 | LipidClass = PS -- SumDBs = 3 | -0.001 | 0.001 | Headgroup = PC -- SumLength = 38 | -0.001 | 0.001 | Headgroup = -- SumLength = 32 | -0.002 | 0.002 |
| LipidClass = LPE -- SumDBs = 4 | -0.001 | 0.001 | Headgroup = CE -- SumDBs = 1 | -0.001 | 0.001 | LipidClass = PS -- SumDBs = 8 | -0.001 | 0.001 | SumDBs = 1 -- SumLength = 34 | -0.002 | 0.002 |
| LipidClass = LPE -- SumLength = 20 | -0.001 | 0.001 | LipidClass = DG -- SumDBs = 2 | -0.001 | 0.001 | Headgroup = PS -- SumDBs = 8 | -0.001 | 0.001 | LipidClass = PC O- -- SumDBs = 0 | -0.002 | 0.002 |
| Headgroup = PE -- SumLength = 20 | -0.001 | 0.001 | LipidClass = CE -- SumDBs = 1 | -0.001 | 0.001 | SumDBs = 2 -- SumLength = 34 | -0.001 | 0.001 | LipidClass = PC O- -- SumLength = 32 | -0.002 | 0.002 |
| SumDBs = 1 -- SumLength = 34 | -0.001 | 0.001 | SumDBs = 3 -- SumLength = 40 | -0.001 | 0.001 | SumDBs = 8 -- SumLength = 40 | -0.001 | 0.001 | LipidClass = LPG -- SumLength = 16 | -0.002 | 0.002 |
| Headgroup = PG -- SumDBs = 6 | -0.001 | 0.001 | Headgroup = PS -- SumDBs = 1 | -0.001 | 0.001 | Headgroup = PE -- SumDBs = 0 | -0.001 | 0.001 | LipidClass = LPG -- SumDBs = 0 | -0.002 | 0.002 |
| Headgroup = PG -- SumDBs = 3 | -0.001 | 0.001 | LipidClass = PS -- SumDBs = 1 | -0.001 | 0.001 | LipidClass = LPE -- SumLength = 20 | -0.001 | 0.001 | Headgroup = PG -- SumLength = 16 | -0.002 | 0.002 |
| Headgroup = -- SumLength = 32 | -0.001 | 0.001 | LipidClass = SM -- SumLength = 40 | -0.001 | 0.001 | LipidClass = LPE -- SumDBs = 4 | -0.001 | 0.001 | Headgroup = PG -- SumDBs = 0 | -0.002 | 0.002 |
| LipidClass = DG -- SumLength = 32 | -0.001 | 0.001 | LipidClass = LPG -- SumDBs = 6 | -0.001 | 0.001 | Headgroup = PE -- SumLength = 20 | -0.001 | 0.001 | LipidClass = PC O- -- SumLength = 38 | -0.002 | 0.002 |
| SumDBs = 5 -- SumLength = 52 | -0.001 | 0.001 | Headgroup = PG -- SumLength = 22 | -0.001 | 0.001 | Headgroup = CE -- SumLength = 18 | -0.001 | 0.001 | Headgroup = -- SumDBs = 5 | -0.002 | 0.002 |
| Headgroup = PG -- SumDBs = 0 | -0.001 | 0.001 | LipidClass = LPG -- SumLength = 22 | -0.001 | 0.001 | LipidClass = CE -- SumLength = 18 | -0.001 | 0.001 | LipidClass = SM -- SumLength = 36 | -0.002 | 0.002 |
| LipidClass = LPG -- SumLength = 16 | -0.001 | 0.001 | LipidClass = DG -- SumLength = 34 | -0.001 | 0.001 | Headgroup = PG -- SumDBs = 5 | -0.001 | 0.001 | Headgroup = PG -- SumLength = 32 | -0.002 | 0.002 |
| Headgroup = PG -- SumLength = 16 | -0.001 | 0.001 | LipidClass = LPE -- SumLength = 18 | -0.001 | 0.001 | LipidClass = PG -- SumDBs = 5 | -0.001 | 0.001 | LipidClass = PG -- SumLength = 32 | -0.002 | 0.002 |
| LipidClass = LPG -- SumDBs = 0 | -0.001 | 0.001 | Headgroup = PE -- SumLength = 18 | -0.001 | 0.001 | Headgroup = PI -- SumLength = 20 | -0.001 | 0.001 | LipidClass = PE O- -- SumLength = 40 | -0.001 | 0.001 |
| SumDBs = 6 -- SumLength = 38 | -0.001 | 0.001 | SumDBs = 5 -- SumLength = 54 | -0.001 | 0.001 | LipidClass = LPI -- SumLength = 20 | -0.001 | 0.001 | LipidClass = PE -- SumDBs = 2 | -0.001 | 0.001 |
| SumDBs = 1 -- SumLength = 32 | -0.001 | 0.001 | SumDBs = 2 -- SumLength = 50 | -0.001 | 0.001 | LipidClass = LPI -- SumDBs = 4 | -0.001 | 0.001 | SumDBs = 5 -- SumLength = 36 | -0.001 | 0.001 |
| LipidClass = Cer -- SumDBs = 3 | -0.001 | 0.001 | Headgroup = PC -- SumLength = 36 | -0.001 | 0.001 | LipidClass = PG -- SumDBs = 3 | -0.001 | 0.001 | LipidClass = Cer -- SumLength = 34 | -0.001 | 0.001 |
| LipidClass = PG -- SumDBs = 1 | -0.001 | 0.001 | SumDBs = 3 -- SumLength = 54 | -0.001 | 0.001 | SumDBs = 1 -- SumLength = 36 | -0.001 | 0.001 | SumDBs = 6 -- SumLength = 22 | -0.001 | 0.001 |
| SumDBs = 4 -- SumLength = 54 | -0.001 | 0.001 | SumDBs = 4 -- SumLength = 40 | -0.001 | 0.001 | Headgroup = PC -- SumLength = 36 | -0.001 | 0.001 | Headgroup = -- SumLength = 38 | -0.001 | 0.001 |
| Headgroup = -- SumLength = 56 | -0.001 | 0.001 | Headgroup = PC -- SumLength = 41 | -0.001 | 0.001 | Headgroup = PI -- SumDBs = 0 | -0.001 | 0.001 | LipidClass = TG -- SumLength = 50 | -0.001 | 0.001 |
| LipidClass = TG -- SumLength = 56 | -0.001 | 0.001 | LipidClass = SM -- SumLength = 41 | -0.001 | 0.001 | LipidClass = LPI -- SumDBs = 0 | -0.001 | 0.001 | Headgroup = -- SumLength = 50 | -0.001 | 0.001 |
| Headgroup = -- SumLength = 34 | -0.001 | 0.001 | Headgroup = -- SumLength = 34 | -0.001 | 0.001 | SumDBs = 2 -- SumLength = 42 | -0.001 | 0.001 | SumDBs = 1 -- SumLength = 36 | -0.001 | 0.001 |
| Headgroup = PG -- SumLength = 32 | -0.001 | 0.001 | LipidClass = LPG -- SumDBs = 1 | -0.001 | 0.001 | LipidClass = DG -- SumDBs = 7 | -0.001 | 0.001 | Headgroup = PG -- SumDBs = 3 | -0.001 | 0.001 |
| LipidClass = PG -- SumLength = 32 | -0.001 | 0.001 | LipidClass = DG -- SumLength = 38 | -0.001 | 0.001 | LipidClass = PC O- -- SumDBs = 1 | -0.001 | 0.001 | SumDBs = 4 -- SumLength = 36 | -0.001 | 0.001 |
| SumDBs = 1 -- SumLength = 38 | -0.001 | 0.001 | Headgroup = -- SumDBs = 8 | -0.001 | 0.001 | LipidClass = PC O- -- SumLength = 34 | -0.001 | 0.001 | SumDBs = 5 -- SumLength = 52 | -0.001 | 0.001 |
| LipidClass = PE -- SumLength = 40 | -0.001 | 0.001 | LipidClass = TG -- SumDBs = 8 | -0.001 | 0.001 | LipidClass = PI -- SumDBs = 3 | -0.001 | 0.001 | LipidClass = DG -- SumDBs = 3 | -0.001 | 0.001 |
| LipidClass = PE O- -- SumLength = 36 | -0.001 | 0.001 | LipidClass = SM -- SumDBs = 2 | -0.001 | 0.001 | Headgroup = PI -- SumDBs = 3 | -0.001 | 0.001 | SumDBs = 7 -- SumLength = 38 | -0.001 | 0.001 |
| SumDBs = 8 -- SumLength = 40 | -0.001 | 0.001 | Headgroup = PS -- SumLength = 38 | -0.001 | 0.001 | SumDBs = 3 -- SumLength = 36 | -0.0 | 0.0 | Headgroup = CE -- SumDBs = 0 | -0.001 | 0.001 |
| Headgroup = -- SumDBs = 0 | -0.001 | 0.001 | LipidClass = PS -- SumLength = 38 | -0.001 | 0.001 | Headgroup = PC -- SumLength = 40 | -0.0 | 0.0 | LipidClass = CE -- SumDBs = 0 | -0.001 | 0.001 |
| LipidClass = DG -- SumDBs = 5 | -0.001 | 0.001 | Headgroup = CE -- SumLength = 20 | -0.001 | 0.001 | LipidClass = LPC -- SumLength = 18 | -0.0 | 0.0 | Headgroup = PS -- SumDBs = 2 | -0.001 | 0.001 |
| LipidClass = DG -- SumDBs = 0 | -0.001 | 0.001 | LipidClass = CE -- SumLength = 20 | -0.001 | 0.001 | Headgroup = PC -- SumLength = 18 | -0.0 | 0.0 | LipidClass = PS -- SumDBs = 2 | -0.001 | 0.001 |
| LipidClass = PS -- SumDBs = 2 | -0.001 | 0.001 | Headgroup = PG -- SumDBs = 1 | -0.001 | 0.001 | SumDBs = 5 -- SumLength = 56 | -0.0 | 0.0 | LipidClass = DG -- SumDBs = 5 | -0.001 | 0.001 |
| Headgroup = PS -- SumDBs = 2 | -0.001 | 0.001 | LipidClass = DG -- SumDBs = 7 | -0.001 | 0.001 | Headgroup = PE -- LipidClass = PE | -0.0 | 0.0 | LipidClass = PC -- SumLength = 32 | -0.001 | 0.001 |
| SumDBs = 3 -- SumLength = 18 | -0.001 | 0.001 | Headgroup = PS -- SumLength = 40 | -0.001 | 0.001 | Headgroup = PC -- SumDBs = 0 | -0.0 | 0.0 | Headgroup = PE -- SumDBs = 3 | -0.001 | 0.001 |
| SumDBs = 7 -- SumLength = 58 | -0.001 | 0.001 | LipidClass = PS -- SumLength = 40 | -0.001 | 0.001 | Headgroup = PE -- SumDBs = 6 | -0.0 | 0.0 | LipidClass = PE O- -- SumLength = 34 | -0.001 | 0.001 |
| SumDBs = 6 -- SumLength = 56 | -0.001 | 0.001 | LipidClass = PC -- SumDBs = 2 | -0.001 | 0.001 | SumDBs = 7 -- SumLength = 58 | -0.0 | 0.0 | SumDBs = 8 -- SumLength = 40 | -0.001 | 0.001 |
| Headgroup = PS -- SumLength = 40 | -0.001 | 0.001 | SumDBs = 3 -- SumLength = 36 | -0.001 | 0.001 | SumDBs = 5 -- SumLength = 54 | -0.0 | 0.0 | LipidClass = Cer -- SumLength = 41 | -0.001 | 0.001 |
| LipidClass = PS -- SumLength = 40 | -0.001 | 0.001 | LipidClass = SM -- SumLength = 38 | -0.001 | 0.001 | LipidClass = LPG -- SumDBs = 6 | -0.0 | 0.0 | LipidClass = PE O- -- SumDBs = 2 | -0.001 | 0.001 |
| LipidClass = PG -- SumDBs = 3 | -0.001 | 0.001 | LipidClass = LPC -- SumLength = 18 | -0.001 | 0.001 | Headgroup = PG -- SumLength = 22 | -0.0 | 0.0 | Headgroup = -- SumLength = 41 | -0.001 | 0.001 |
| LipidClass = TG -- SumLength = 58 | -0.001 | 0.001 | Headgroup = PC -- SumLength = 18 | -0.001 | 0.001 | LipidClass = LPG -- SumLength = 22 | -0.0 | 0.0 | Headgroup = PE -- SumDBs = 1 | -0.001 | 0.001 |
| Headgroup = -- SumLength = 58 | -0.001 | 0.001 | LipidClass = TG -- SumDBs = 12 | -0.001 | 0.001 | SumDBs = 3 -- SumLength = 56 | -0.0 | 0.0 | LipidClass = PE -- SumLength = 36 | -0.001 | 0.001 |
| Headgroup = PG -- SumLength = 34 | -0.001 | 0.001 | Headgroup = -- SumDBs = 12 | -0.001 | 0.001 | Headgroup = PE -- LipidClass = PE O- | -0.0 | 0.0 | Headgroup = PE -- SumLength = 40 | -0.001 | 0.001 |
| LipidClass = PG -- SumLength = 34 | -0.001 | 0.001 | LipidClass = LPG -- SumDBs = 2 | -0.001 | 0.001 | Headgroup = PS -- LipidClass = PS | -0.0 | 0.0 | LipidClass = DG -- SumDBs = 4 | -0.001 | 0.001 |
| LipidClass = PG -- SumDBs = 4 | -0.001 | 0.001 | Headgroup = PG -- SumDBs = 6 | -0.001 | 0.001 | LipidClass = PC O- -- SumLength = 40 | -0.0 | 0.0 | LipidClass = PS -- SumDBs = 7 | -0.001 | 0.001 |
| Headgroup = PE -- SumLength = 22 | -0.001 | 0.001 | Headgroup = -- SumDBs = 3 | -0.001 | 0.001 | Headgroup = PE -- LipidClass = LPE | -0.0 | 0.0 | Headgroup = PS -- SumDBs = 7 | -0.001 | 0.001 |
| LipidClass = LPE -- SumDBs = 6 | -0.001 | 0.001 | LipidClass = CE -- SumDBs = 4 | -0.001 | 0.001 | LipidClass = PC -- SumDBs = 1 | -0.0 | 0.0 | LipidClass = SM -- SumLength = 41 | -0.001 | 0.001 |
| LipidClass = LPE -- SumLength = 22 | -0.001 | 0.001 | Headgroup = CE -- SumDBs = 4 | -0.001 | 0.001 | LipidClass = PC -- SumDBs = 5 | -0.0 | 0.0 | Headgroup = PC -- SumLength = 41 | -0.001 | 0.001 |
| SumDBs = 6 -- SumLength = 22 | -0.001 | 0.001 | LipidClass = LPG -- SumLength = 18 | -0.001 | 0.001 | SumDBs = 2 -- SumLength = 40 | -0.0 | 0.0 | SumDBs = 3 -- SumLength = 50 | -0.001 | 0.001 |
| LipidClass = LPG -- SumDBs = 2 | -0.001 | 0.001 | Headgroup = PG -- SumLength = 18 | -0.001 | 0.001 | Headgroup = PC -- LipidClass = SM | -0.0 | 0.0 | LipidClass = PI -- SumDBs = 4 | -0.001 | 0.001 |
| Headgroup = -- SumDBs = 7 | -0.001 | 0.001 | LipidClass = PC -- SumLength = 40 | -0.001 | 0.001 | Headgroup = P -- LipidClass = PA | -0.0 | 0.0 | Headgroup = -- SumDBs = 6 | -0.001 | 0.001 |
| LipidClass = PS -- SumLength = 36 | -0.0 | 0.0 | Headgroup = -- SumLength = 62 | -0.001 | 0.001 | Headgroup = CE -- SumLength = 20 | -0.0 | 0.0 | LipidClass = PS -- SumDBs = 8 | -0.001 | 0.001 |
| Headgroup = PS -- SumLength = 36 | -0.0 | 0.0 | LipidClass = TG -- SumLength = 62 | -0.001 | 0.001 | LipidClass = CE -- SumLength = 20 | -0.0 | 0.0 | Headgroup = PS -- SumDBs = 8 | -0.001 | 0.001 |
| LipidClass = LPC -- SumDBs = 4 | -0.0 | 0.0 | Headgroup = -- SumLength = 52 | -0.001 | 0.001 | Headgroup = CE -- SumDBs = 3 | -0.0 | 0.0 | Headgroup = PC -- SumDBs = 4 | -0.001 | 0.001 |
| Headgroup = PC -- SumLength = 20 | -0.0 | 0.0 | LipidClass = TG -- SumLength = 52 | -0.001 | 0.001 | LipidClass = CE -- SumDBs = 3 | -0.0 | 0.0 | LipidClass = SM -- SumDBs = 3 | -0.001 | 0.001 |
| LipidClass = LPC -- SumLength = 20 | -0.0 | 0.0 | SumDBs = 12 -- SumLength = 60 | -0.001 | 0.001 | LipidClass = LPE -- SumLength = 16 | -0.0 | 0.0 | SumDBs = 2 -- SumLength = 32 | -0.001 | 0.001 |
| Headgroup = PC -- SumLength = 36 | -0.0 | 0.0 | LipidClass = TG -- SumLength = 54 | -0.001 | 0.001 | Headgroup = PE -- SumLength = 16 | -0.0 | 0.0 | Headgroup = -- SumDBs = 0 | -0.001 | 0.001 |
| LipidClass = PS -- SumDBs = 8 | -0.0 | 0.0 | Headgroup = -- SumLength = 54 | -0.001 | 0.001 | Headgroup = PG -- LipidClass = PG | -0.0 | 0.0 | Headgroup = PE -- SumLength = 18 | -0.001 | 0.001 |
| Headgroup = PS -- SumDBs = 8 | -0.0 | 0.0 | LipidClass = PE O- -- SumDBs = 4 | -0.0 | 0.0 | LipidClass = DG -- SumLength = 38 | -0.0 | 0.0 | LipidClass = LPE -- SumLength = 18 | -0.001 | 0.001 |
| SumDBs = 5 -- SumLength = 40 | -0.0 | 0.0 | LipidClass = DG -- SumDBs = 4 | -0.0 | 0.0 | LipidClass = LPC -- SumDBs = 0 | -0.0 | 0.0 | LipidClass = PS -- SumDBs = 5 | -0.001 | 0.001 |
| SumDBs = 2 -- SumLength = 18 | -0.0 | 0.0 | LipidClass = PS -- SumDBs = 7 | -0.0 | 0.0 | Headgroup = PC -- LipidClass = PC | -0.0 | 0.0 | Headgroup = PS -- SumDBs = 5 | -0.001 | 0.001 |
| SumDBs = 1 -- SumLength = 42 | -0.0 | 0.0 | Headgroup = PS -- SumDBs = 7 | -0.0 | 0.0 | LipidClass = DG -- SumLength = 34 | -0.0 | 0.0 | Headgroup = PE -- SumLength = 36 | -0.001 | 0.001 |
| LipidClass = TG -- SumDBs = 5 | -0.0 | 0.0 | LipidClass = SM -- SumDBs = 1 | -0.0 | 0.0 | LipidClass = PG -- SumDBs = 4 | -0.0 | 0.0 | LipidClass = PC -- SumDBs = 0 | -0.001 | 0.001 |
| Headgroup = PC -- SumDBs = 2 | -0.0 | 0.0 | LipidClass = PC -- SumDBs = 1 | -0.0 | 0.0 | SumDBs = 6 -- SumLength = 22 | -0.0 | 0.0 | LipidClass = PC -- SumDBs = 7 | -0.001 | 0.001 |
| SumDBs = 2 -- SumLength = 32 | -0.0 | 0.0 | SumDBs = 4 -- SumLength = 20 | -0.0 | 0.0 | Headgroup = -- SumLength = 36 | -0.0 | 0.0 | LipidClass = PE -- SumDBs = 3 | -0.001 | 0.001 |
| LipidClass = PE -- SumLength = 34 | -0.0 | 0.0 | Headgroup = CE -- SumDBs = 2 | -0.0 | 0.0 | LipidClass = DG -- SumLength = 36 | -0.0 | 0.0 | SumDBs = 6 -- SumLength = 54 | -0.001 | 0.001 |
| Headgroup = PE -- SumLength = 34 | -0.0 | 0.0 | LipidClass = CE -- SumDBs = 2 | -0.0 | 0.0 | Headgroup = -- LipidClass = DG | -0.0 | 0.0 | LipidClass = DG -- SumDBs = 0 | -0.001 | 0.001 |
| LipidClass = PE -- SumLength = 36 | -0.0 | 0.0 | SumDBs = 3 -- SumLength = 20 | -0.0 | 0.0 | SumDBs = 10 -- SumLength = 60 | -0.0 | 0.0 | SumDBs = 8 -- SumLength = 58 | -0.001 | 0.001 |
| Headgroup = PC -- SumDBs = 5 | -0.0 | 0.0 | LipidClass = PG -- SumLength = 36 | -0.0 | 0.0 | Headgroup = CE -- LipidClass = CE | -0.0 | 0.0 | LipidClass = PC -- SumDBs = 1 | -0.001 | 0.001 |
| SumDBs = 6 -- SumLength = 52 | -0.0 | 0.0 | Headgroup = PG -- SumLength = 36 | -0.0 | 0.0 | SumDBs = 6 -- SumLength = 40 | -0.0 | 0.0 | SumDBs = 6 -- SumLength = 52 | -0.001 | 0.001 |
| Headgroup = -- SumLength = 42 | -0.0 | 0.0 | SumDBs = 2 -- SumLength = 18 | -0.0 | 0.0 | Headgroup = PG -- LipidClass = LPG | -0.0 | 0.0 | LipidClass = LPE -- SumDBs = 1 | -0.001 | 0.001 |
| LipidClass = Cer -- SumLength = 42 | -0.0 | 0.0 | SumDBs = 9 -- SumLength = 58 | -0.0 | 0.0 | Headgroup = PC -- LipidClass = LPC | -0.0 | 0.0 | SumDBs = 6 -- SumLength = 56 | -0.001 | 0.001 |
| Headgroup = PS -- SumDBs = 7 | -0.0 | 0.0 | SumDBs = 6 -- SumLength = 54 | -0.0 | 0.0 | Headgroup = PC -- LipidClass = PC O- | -0.0 | 0.0 | SumDBs = 2 -- SumLength = 50 | -0.001 | 0.001 |
| LipidClass = PS -- SumDBs = 7 | -0.0 | 0.0 | Headgroup = PE -- LipidClass = PE | -0.0 | 0.0 | LipidClass = TG -- SumDBs = 10 | -0.0 | 0.0 | LipidClass = LPE -- SumLength = 20 | -0.001 | 0.001 |
| LipidClass = LPG -- SumDBs = 3 | -0.0 | 0.0 | LipidClass = PE -- SumDBs = 5 | -0.0 | 0.0 | Headgroup = -- SumDBs = 10 | -0.0 | 0.0 | LipidClass = LPE -- SumDBs = 4 | -0.001 | 0.001 |
| SumDBs = 1 -- SumLength = 18 | -0.0 | 0.0 | SumDBs = 10 -- SumLength = 58 | -0.0 | 0.0 | LipidClass = PC O- -- SumLength = 36 | -0.0 | 0.0 | Headgroup = PE -- SumLength = 20 | -0.001 | 0.001 |
| SumDBs = 1 -- SumLength = 36 | -0.0 | 0.0 | Headgroup = PE -- LipidClass = PE O- | -0.0 | 0.0 | Headgroup = CE -- SumDBs = 2 | -0.0 | 0.0 | LipidClass = PC -- SumDBs = 5 | -0.0 | 0.0 |
| LipidClass = PC O- -- SumDBs = 2 | -0.0 | 0.0 | Headgroup = PS -- LipidClass = PS | -0.0 | 0.0 | LipidClass = CE -- SumDBs = 2 | -0.0 | 0.0 | SumDBs = 7 -- SumLength = 54 | -0.0 | 0.0 |
| SumDBs = 3 -- SumLength = 38 | -0.0 | 0.0 | SumDBs = 12 -- SumLength = 62 | -0.0 | 0.0 | LipidClass = SM -- SumLength = 40 | -0.0 | 0.0 | SumDBs = 4 -- SumLength = 54 | -0.0 | 0.0 |
| Headgroup = PE -- SumDBs = 0 | -0.0 | 0.0 | Headgroup = PE -- LipidClass = LPE | -0.0 | 0.0 | Headgroup = PS -- SumDBs = 0 | -0.0 | 0.0 | LipidClass = PE -- SumDBs = 1 | -0.0 | 0.0 |
| LipidClass = LPC -- SumDBs = 2 | -0.0 | 0.0 | Headgroup = -- SumDBs = 13 | -0.0 | 0.0 | SumDBs = 0 -- SumLength = 38 | -0.0 | 0.0 | SumDBs = 2 -- SumLength = 54 | -0.0 | 0.0 |
| SumDBs = 7 -- SumLength = 56 | -0.0 | 0.0 | LipidClass = TG -- SumDBs = 13 | -0.0 | 0.0 | LipidClass = PS -- SumDBs = 0 | -0.0 | 0.0 | LipidClass = TG -- SumDBs = 5 | -0.0 | 0.0 |
| LipidClass = LPE -- SumDBs = 0 | -0.0 | 0.0 | Headgroup = PC -- LipidClass = SM | -0.0 | 0.0 | LipidClass = Cer -- SumDBs = 3 | -0.0 | 0.0 | LipidClass = PE O- -- SumDBs = 3 | -0.0 | 0.0 |
| SumDBs = 6 -- SumLength = 58 | -0.0 | 0.0 | Headgroup = P -- LipidClass = PA | -0.0 | 0.0 | Headgroup = PI -- LipidClass = PI | -0.0 | 0.0 | SumDBs = 4 -- SumLength = 50 | -0.0 | 0.0 |
| LipidClass = SM -- SumDBs = 1 | -0.0 | 0.0 | SumDBs = 7 -- SumLength = 56 | -0.0 | 0.0 | LipidClass = PC O- -- SumDBs = 0 | -0.0 | 0.0 | SumDBs = 0 -- SumLength = 34 | -0.0 | 0.0 |
| LipidClass = CE -- SumDBs = 3 | -0.0 | 0.0 | Headgroup = PG -- LipidClass = PG | -0.0 | 0.0 | LipidClass = PC O- -- SumLength = 32 | -0.0 | 0.0 | SumDBs = 2 -- SumLength = 40 | -0.0 | 0.0 |
| Headgroup = CE -- SumDBs = 3 | -0.0 | 0.0 | Headgroup = CE -- SumDBs = 3 | -0.0 | 0.0 | SumDBs = 3 -- SumLength = 20 | -0.0 | 0.0 | Headgroup = PC -- SumLength = 33 | -0.0 | 0.0 |
| LipidClass = PC O- -- SumDBs = 5 | -0.0 | 0.0 | LipidClass = CE -- SumDBs = 3 | -0.0 | 0.0 | Headgroup = -- LipidClass = TG | -0.0 | 0.0 | SumDBs = 1 -- SumLength = 33 | -0.0 | 0.0 |
| Headgroup = PE -- SumDBs = 8 | -0.0 | 0.0 | Headgroup = PC -- LipidClass = PC | -0.0 | 0.0 | SumDBs = 2 -- SumLength = 48 | -0.0 | 0.0 | LipidClass = SM -- SumLength = 33 | -0.0 | 0.0 |
| LipidClass = PE -- SumDBs = 8 | -0.0 | 0.0 | Headgroup = PS -- SumDBs = 0 | -0.0 | 0.0 | LipidClass = TG -- SumLength = 58 | -0.0 | 0.0 | LipidClass = TG -- SumDBs = 7 | -0.0 | 0.0 |
| SumDBs = 4 -- SumLength = 56 | -0.0 | 0.0 | SumDBs = 0 -- SumLength = 38 | -0.0 | 0.0 | Headgroup = -- SumLength = 58 | -0.0 | 0.0 | Headgroup = PC -- SumDBs = 5 | -0.0 | 0.0 |
| SumDBs = 2 -- SumLength = 54 | -0.0 | 0.0 | LipidClass = PS -- SumDBs = 0 | -0.0 | 0.0 | LipidClass = PS -- SumLength = 40 | -0.0 | 0.0 | LipidClass = PE -- SumDBs = 7 | -0.0 | 0.0 |
| LipidClass = PC -- SumDBs = 7 | -0.0 | 0.0 | SumDBs = 13 -- SumLength = 62 | -0.0 | 0.0 | Headgroup = PS -- SumLength = 40 | -0.0 | 0.0 | Headgroup = -- SumDBs = 9 | -0.0 | 0.0 |
| Headgroup = PI -- SumDBs = 7 | -0.0 | 0.0 | Headgroup = -- SumLength = 50 | -0.0 | 0.0 | Headgroup = -- LipidClass = Cer | -0.0 | 0.0 | LipidClass = TG -- SumDBs = 9 | -0.0 | 0.0 |
| LipidClass = PI -- SumDBs = 7 | -0.0 | 0.0 | LipidClass = TG -- SumLength = 50 | -0.0 | 0.0 | SumDBs = 12 -- SumLength = 60 | -0.0 | 0.0 | SumDBs = 1 -- SumLength = 38 | -0.0 | 0.0 |
| LipidClass = TG -- SumLength = 50 | -0.0 | 0.0 | Headgroup = -- SumDBs = 6 | -0.0 | 0.0 | LipidClass = CE -- SumDBs = 4 | -0.0 | 0.0 | Headgroup = PC -- SumDBs = 0 | -0.0 | 0.0 |
| Headgroup = -- SumLength = 50 | -0.0 | 0.0 | SumDBs = 11 -- SumLength = 62 | -0.0 | 0.0 | Headgroup = CE -- SumDBs = 4 | -0.0 | 0.0 | LipidClass = CE -- SumDBs = 1 | -0.0 | 0.0 |
| SumDBs = 5 -- SumLength = 56 | -0.0 | 0.0 | LipidClass = TG -- SumLength = 56 | -0.0 | 0.0 | LipidClass = PS -- SumDBs = 3 | -0.0 | 0.0 | LipidClass = DG -- SumLength = 36 | -0.0 | 0.0 |
| LipidClass = SM -- SumLength = 36 | -0.0 | 0.0 | Headgroup = -- SumLength = 56 | -0.0 | 0.0 | Headgroup = PS -- SumDBs = 3 | -0.0 | 0.0 | Headgroup = -- SumLength = 36 | -0.0 | 0.0 |
| Headgroup = PG -- SumDBs = 4 | -0.0 | 0.0 | Headgroup = -- LipidClass = DG | -0.0 | 0.0 | SumDBs = 9 -- SumLength = 60 | -0.0 | 0.0 | SumDBs = 6 -- SumLength = 36 | -0.0 | 0.0 |
| Headgroup = PC -- SumLength = 33 | -0.0 | 0.0 | Headgroup = CE -- LipidClass = CE | -0.0 | 0.0 | Headgroup = -- SumDBs = 12 | -0.0 | 0.0 | SumDBs = 9 -- SumLength = 60 | -0.0 | 0.0 |
| SumDBs = 1 -- SumLength = 33 | -0.0 | 0.0 | LipidClass = LPC -- SumDBs = 1 | -0.0 | 0.0 | LipidClass = TG -- SumDBs = 12 | -0.0 | 0.0 | LipidClass = TG -- SumLength = 48 | -0.0 | 0.0 |
| LipidClass = SM -- SumLength = 33 | -0.0 | 0.0 | Headgroup = PG -- LipidClass = LPG | -0.0 | 0.0 | SumDBs = 4 -- SumLength = 56 | -0.0 | 0.0 | Headgroup = -- SumLength = 48 | -0.0 | 0.0 |
| Headgroup = PC -- SumLength = 18 | -0.0 | 0.0 | Headgroup = PC -- LipidClass = LPC | -0.0 | 0.0 | SumDBs = 11 -- SumLength = 60 | -0.0 | 0.0 | Headgroup = CE -- SumDBs = 1 | -0.0 | 0.0 |
| LipidClass = LPC -- SumLength = 18 | -0.0 | 0.0 | Headgroup = PC -- LipidClass = PC O- | -0.0 | 0.0 | SumDBs = 8 -- SumLength = 60 | -0.0 | 0.0 | SumDBs = 7 -- SumLength = 56 | -0.0 | 0.0 |
| Headgroup = -- SumDBs = 6 | -0.0 | 0.0 | SumDBs = 3 -- SumLength = 48 | -0.0 | 0.0 | Headgroup = PC -- SumLength = 34 | -0.0 | 0.0 | Headgroup = -- SumLength = 34 | -0.0 | 0.0 |
| SumDBs = 0 -- SumLength = 34 | -0.0 | 0.0 | Headgroup = -- SumDBs = 11 | -0.0 | 0.0 | SumDBs = 3 -- SumLength = 40 | -0.0 | 0.0 | SumDBs = 4 -- SumLength = 34 | -0.0 | 0.0 |
| LipidClass = PC -- SumDBs = 5 | -0.0 | 0.0 | LipidClass = TG -- SumDBs = 11 | -0.0 | 0.0 | Headgroup = -- SumDBs = 7 | -0.0 | 0.0 | Headgroup = PE -- SumDBs = 8 | -0.0 | 0.0 |
| LipidClass = PE -- SumDBs = 3 | -0.0 | 0.0 | Headgroup = PI -- LipidClass = PI | -0.0 | 0.0 | SumDBs = 7 -- SumLength = 60 | -0.0 | 0.0 | LipidClass = PE -- SumDBs = 8 | -0.0 | 0.0 |
| LipidClass = PG -- SumLength = 36 | -0.0 | 0.0 | Headgroup = PS -- SumDBs = 5 | -0.0 | 0.0 | LipidClass = PS -- SumDBs = 1 | -0.0 | 0.0 | LipidClass = LPC -- SumDBs = 1 | -0.0 | 0.0 |
| Headgroup = PG -- SumLength = 36 | -0.0 | 0.0 | LipidClass = PS -- SumDBs = 5 | -0.0 | 0.0 | Headgroup = PS -- SumDBs = 1 | -0.0 | 0.0 | SumDBs = 3 -- SumLength = 38 | -0.0 | 0.0 |
| LipidClass = LPC -- SumDBs = 1 | -0.0 | 0.0 | Headgroup = -- LipidClass = TG | -0.0 | 0.0 | SumDBs = 6 -- SumLength = 58 | -0.0 | 0.0 | LipidClass = LPG -- SumDBs = 2 | -0.0 | 0.0 |
| SumDBs = 9 -- SumLength = 56 | -0.0 | 0.0 | Headgroup = PC -- SumDBs = 0 | -0.0 | 0.0 | LipidClass = PE -- SumLength = 40 | -0.0 | 0.0 | LipidClass = TG -- SumLength = 46 | -0.0 | 0.0 |
| Headgroup = PG -- SumLength = 18 | -0.0 | 0.0 | Headgroup = -- LipidClass = Cer | -0.0 | 0.0 | SumDBs = 0 -- SumLength = 34 | -0.0 | 0.0 | Headgroup = -- SumLength = 46 | -0.0 | 0.0 |
| LipidClass = LPG -- SumLength = 18 | -0.0 | 0.0 | Headgroup = -- SumDBs = 14 | -0.0 | 0.0 | SumDBs = 1 -- SumLength = 48 | -0.0 | 0.0 | LipidClass = LPE -- SumDBs = 6 | -0.0 | 0.0 |
| Headgroup = -- SumLength = 46 | -0.0 | 0.0 | SumDBs = 14 -- SumLength = 62 | -0.0 | 0.0 | LipidClass = TG -- SumDBs = 11 | -0.0 | 0.0 | LipidClass = LPE -- SumLength = 22 | -0.0 | 0.0 |
| LipidClass = TG -- SumLength = 46 | -0.0 | 0.0 | LipidClass = TG -- SumDBs = 14 | -0.0 | 0.0 | Headgroup = -- SumDBs = 11 | -0.0 | 0.0 | Headgroup = PE -- SumLength = 22 | -0.0 | 0.0 |
| SumDBs = 5 -- SumLength = 20 | -0.0 | 0.0 | SumDBs = 5 -- SumLength = 48 | -0.0 | 0.0 | LipidClass = DG -- SumDBs = 6 | -0.0 | 0.0 | SumDBs = 7 -- SumLength = 42 | -0.0 | 0.0 |
| LipidClass = CE -- SumDBs = 5 | -0.0 | 0.0 | Headgroup = -- SumDBs = 10 | -0.0 | 0.0 | SumDBs = 4 -- SumLength = 58 | -0.0 | 0.0 | LipidClass = PG -- SumDBs = 6 | -0.0 | 0.0 |
| Headgroup = CE -- SumDBs = 5 | -0.0 | 0.0 | LipidClass = TG -- SumDBs = 10 | -0.0 | 0.0 | Headgroup = -- SumLength = 48 | -0.0 | 0.0 | LipidClass = PI -- SumDBs = 7 | -0.0 | 0.0 |
| SumDBs = 3 -- SumLength = 50 | -0.0 | 0.0 | LipidClass = LPE -- SumDBs = 0 | -0.0 | 0.0 | LipidClass = TG -- SumLength = 48 | -0.0 | 0.0 | Headgroup = PI -- SumDBs = 7 | -0.0 | 0.0 |
| SumDBs = 4 -- SumLength = 34 | -0.0 | 0.0 | SumDBs = 4 -- SumLength = 48 | -0.0 | 0.0 | Headgroup = -- SumLength = 41 | -0.0 | 0.0 | SumDBs = 7 -- SumLength = 52 | -0.0 | 0.0 |
| Headgroup = PG -- SumDBs = 1 | -0.0 | 0.0 | SumDBs = 5 -- SumLength = 34 | -0.0 | 0.0 | LipidClass = Cer -- SumLength = 41 | -0.0 | 0.0 | SumDBs = 8 -- SumLength = 60 | -0.0 | 0.0 |
| SumDBs = 4 -- SumLength = 50 | -0.0 | 0.0 | LipidClass = PC -- SumLength = 36 | -0.0 | 0.0 | SumDBs = 2 -- SumLength = 38 | -0.0 | 0.0 | SumDBs = 7 -- SumLength = 60 | -0.0 | 0.0 |
| SumDBs = 6 -- SumLength = 36 | -0.0 | 0.0 | SumDBs = 13 -- SumLength = 60 | -0.0 | 0.0 | Headgroup = PC -- SumDBs = 4 | -0.0 | 0.0 | SumDBs = 5 -- SumLength = 50 | -0.0 | 0.0 |
| Headgroup = PE -- SumDBs = 2 | -0.0 | 0.0 | SumDBs = 3 -- SumLength = 58 | -0.0 | 0.0 | SumDBs = 5 -- SumLength = 34 | -0.0 | 0.0 | SumDBs = 5 -- SumLength = 20 | -0.0 | 0.0 |
| LipidClass = PG -- SumDBs = 0 | -0.0 | 0.0 | SumDBs = 11 -- SumLength = 58 | -0.0 | 0.0 | LipidClass = PE -- SumDBs = 5 | -0.0 | 0.0 | LipidClass = CE -- SumDBs = 5 | -0.0 | 0.0 |
| SumDBs = 8 -- SumLength = 60 | -0.0 | 0.0 | SumDBs = 2 -- SumLength = 38 | -0.0 | 0.0 | LipidClass = PC O- -- SumDBs = 6 | -0.0 | 0.0 | LipidClass = TG -- SumDBs = 0 | -0.0 | 0.0 |
| Headgroup = PI -- SumLength = 32 | -0.0 | 0.0 | SumDBs = 3 -- SumLength = 56 | -0.0 | 0.0 | SumDBs = 8 -- SumLength = 42 | -0.0 | 0.0 | Headgroup = CE -- SumDBs = 5 | -0.0 | 0.0 |
| LipidClass = PI -- SumDBs = 0 | -0.0 | 0.0 | SumDBs = 4 -- SumLength = 46 | -0.0 | 0.0 | Headgroup = PE -- SumLength = 42 | -0.0 | 0.0 | LipidClass = PE -- SumLength = 32 | -0.0 | 0.0 |
| LipidClass = PI -- SumLength = 32 | -0.0 | 0.0 | SumDBs = 8 -- SumLength = 42 | -0.0 | 0.0 | LipidClass = PE -- SumLength = 42 | -0.0 | 0.0 | Headgroup = PE -- SumLength = 32 | -0.0 | 0.0 |
| SumDBs = 4 -- SumLength = 36 | -0.0 | 0.0 | SumDBs = 3 -- SumLength = 60 | -0.0 | 0.0 | LipidClass = PG -- SumDBs = 0 | -0.0 | 0.0 | Headgroup = PE -- SumLength = 42 | -0.0 | 0.0 |
| LipidClass = PE O- -- SumDBs = 6 | -0.0 | 0.0 | Headgroup = -- SumDBs = 7 | -0.0 | 0.0 | SumDBs = 2 -- SumLength = 58 | -0.0 | 0.0 | LipidClass = PE -- SumLength = 42 | -0.0 | 0.0 |
| SumDBs = 7 -- SumLength = 52 | -0.0 | 0.0 | LipidClass = LPE O- -- SumDBs = 1 | 0.0 | 0.0 | SumDBs = 13 -- SumLength = 60 | -0.0 | 0.0 | SumDBs = 8 -- SumLength = 54 | -0.0 | 0.0 |
| SumDBs = 2 -- SumLength = 50 | -0.0 | 0.0 | Headgroup = PE -- LipidClass = LPE O- | 0.0 | 0.0 | SumDBs = 4 -- SumLength = 60 | -0.0 | 0.0 | SumDBs = 1 -- SumLength = 48 | -0.0 | 0.0 |
| LipidClass = TG -- SumDBs = 0 | -0.0 | 0.0 | LipidClass = LPE O- -- SumLength = 16 | 0.0 | 0.0 | SumDBs = 3 -- SumLength = 60 | -0.0 | 0.0 | LipidClass = PE -- SumDBs = 0 | -0.0 | 0.0 |
| SumDBs = 8 -- SumLength = 54 | -0.0 | 0.0 | SumDBs = 2 -- SumLength = 60 | 0.0 | 0.0 | SumDBs = 2 -- SumLength = 46 | -0.0 | 0.0 | Headgroup = -- SumLength = 58 | -0.0 | 0.0 |
| LipidClass = PE -- SumDBs = 2 | -0.0 | 0.0 | SumDBs = 6 -- SumLength = 48 | 0.0 | 0.0 | SumDBs = 7 -- SumLength = 50 | -0.0 | 0.0 | LipidClass = TG -- SumLength = 58 | -0.0 | 0.0 |
| SumDBs = 8 -- SumLength = 58 | -0.0 | 0.0 | SumDBs = 2 -- SumLength = 58 | 0.0 | 0.0 | SumDBs = 10 -- SumLength = 56 | -0.0 | 0.0 | SumDBs = 0 -- SumLength = 48 | -0.0 | 0.0 |
| SumDBs = 2 -- SumLength = 46 | -0.0 | 0.0 | SumDBs = 4 -- SumLength = 60 | 0.0 | 0.0 | SumDBs = 3 -- SumLength = 46 | -0.0 | 0.0 | SumDBs = 9 -- SumLength = 58 | -0.0 | 0.0 |
| SumDBs = 9 -- SumLength = 60 | -0.0 | 0.0 | SumDBs = 1 -- SumLength = 58 | 0.0 | 0.0 | SumDBs = 2 -- SumLength = 60 | -0.0 | 0.0 | SumDBs = 6 -- SumLength = 58 | -0.0 | 0.0 |
| SumDBs = 5 -- SumLength = 50 | -0.0 | 0.0 | SumDBs = 7 -- SumLength = 50 | 0.0 | 0.0 | LipidClass = LPE O- -- SumDBs = 1 | 0.0 | 0.0 | SumDBs = 2 -- SumLength = 48 | -0.0 | 0.0 |
| SumDBs = 1 -- SumLength = 48 | -0.0 | 0.0 | SumDBs = 9 -- SumLength = 56 | 0.0 | 0.0 | Headgroup = PE -- LipidClass = LPE O- | 0.0 | 0.0 | SumDBs = 6 -- SumLength = 50 | -0.0 | 0.0 |
| SumDBs = 6 -- SumLength = 60 | -0.0 | 0.0 | SumDBs = 6 -- SumLength = 36 | 0.0 | 0.0 | LipidClass = LPE O- -- SumLength = 16 | 0.0 | 0.0 | SumDBs = 1 -- SumLength = 46 | -0.0 | 0.0 |
| SumDBs = 1 -- SumLength = 46 | -0.0 | 0.0 | SumDBs = 6 -- SumLength = 50 | 0.0 | 0.0 | SumDBs = 6 -- SumLength = 48 | 0.0 | 0.0 | SumDBs = 2 -- SumLength = 46 | -0.0 | 0.0 |
| SumDBs = 0 -- SumLength = 48 | -0.0 | 0.0 | SumDBs = 0 -- SumLength = 46 | 0.0 | 0.0 | SumDBs = 2 -- SumLength = 56 | 0.0 | 0.0 | SumDBs = 4 -- SumLength = 46 | -0.0 | 0.0 |
| SumDBs = 7 -- SumLength = 54 | -0.0 | 0.0 | LipidClass = PG -- SumDBs = 0 | 0.0 | 0.0 | SumDBs = 1 -- SumLength = 58 | 0.0 | 0.0 | SumDBs = 0 -- SumLength = 46 | -0.0 | 0.0 |
| SumDBs = 7 -- SumLength = 60 | -0.0 | 0.0 | Headgroup = PE -- SumDBs = 0 | 0.0 | 0.0 | SumDBs = 0 -- SumLength = 48 | 0.0 | 0.0 | SumDBs = 6 -- SumLength = 60 | -0.0 | 0.0 |
| Headgroup = PE -- SumDBs = 3 | -0.0 | 0.0 | SumDBs = 2 -- SumLength = 56 | 0.0 | 0.0 | SumDBs = 3 -- SumLength = 58 | 0.0 | 0.0 | Headgroup = PI -- SumLength = 32 | -0.0 | 0.0 |
| LipidClass = PE O- -- SumDBs = 2 | -0.0 | 0.0 | LipidClass = PE -- SumDBs = 0 | 0.0 | 0.0 | SumDBs = 1 -- SumLength = 46 | 0.0 | 0.0 | LipidClass = PI -- SumLength = 32 | -0.0 | 0.0 |
| SumDBs = 4 -- SumLength = 58 | -0.0 | 0.0 | SumDBs = 3 -- SumLength = 46 | 0.0 | 0.0 | SumDBs = 7 -- SumLength = 42 | 0.0 | 0.0 | LipidClass = PI -- SumDBs = 0 | -0.0 | 0.0 |
| SumDBs = 10 -- SumLength = 56 | -0.0 | 0.0 | SumDBs = 10 -- SumLength = 56 | 0.0 | 0.0 | SumDBs = 11 -- SumLength = 58 | 0.0 | 0.0 | SumDBs = 3 -- SumLength = 48 | -0.0 | 0.0 |
| SumDBs = 3 -- SumLength = 46 | -0.0 | 0.0 | SumDBs = 4 -- SumLength = 58 | 0.0 | 0.0 | SumDBs = 6 -- SumLength = 60 | 0.0 | 0.0 | SumDBs = 2 -- SumLength = 56 | -0.0 | 0.0 |
| SumDBs = 0 -- SumLength = 46 | -0.0 | 0.0 | LipidClass = LPI -- SumDBs = 0 | 0.0 | 0.0 | SumDBs = 6 -- SumLength = 50 | 0.0 | 0.0 | SumDBs = 10 -- SumLength = 56 | -0.0 | 0.0 |
| SumDBs = 2 -- SumLength = 56 | -0.0 | 0.0 | SumDBs = 6 -- SumLength = 60 | 0.0 | 0.0 | LipidClass = PC -- SumDBs = 0 | 0.0 | 0.0 | SumDBs = 3 -- SumLength = 46 | -0.0 | 0.0 |
| LipidClass = PE -- SumLength = 32 | -0.0 | 0.0 | LipidClass = PI -- SumLength = 32 | 0.0 | 0.0 | Headgroup = -- SumDBs = 14 | 0.0 | 0.0 | SumDBs = 11 -- SumLength = 60 | -0.0 | 0.0 |
| Headgroup = PE -- SumLength = 32 | -0.0 | 0.0 | LipidClass = PI -- SumDBs = 0 | 0.0 | 0.0 | LipidClass = TG -- SumDBs = 14 | 0.0 | 0.0 | SumDBs = 5 -- SumLength = 48 | -0.0 | 0.0 |
| SumDBs = 2 -- SumLength = 48 | -0.0 | 0.0 | Headgroup = PI -- SumLength = 32 | 0.0 | 0.0 | SumDBs = 14 -- SumLength = 62 | 0.0 | 0.0 | LipidClass = PE O- -- SumDBs = 6 | -0.0 | 0.0 |
| Headgroup = -- SumLength = 48 | -0.0 | 0.0 | LipidClass = PE -- SumLength = 32 | 0.0 | 0.0 | SumDBs = 0 -- SumLength = 46 | 0.0 | 0.0 | SumDBs = 1 -- SumLength = 58 | -0.0 | 0.0 |
| LipidClass = TG -- SumLength = 48 | -0.0 | 0.0 | Headgroup = PE -- SumLength = 32 | 0.0 | 0.0 | Headgroup = PI -- SumLength = 32 | 0.0 | 0.0 | SumDBs = 7 -- SumLength = 50 | -0.0 | 0.0 |
| Headgroup = PE -- SumLength = 42 | -0.0 | 0.0 | Headgroup = PC -- SumDBs = 4 | 0.0 | 0.0 | LipidClass = PI -- SumDBs = 0 | 0.0 | 0.0 | SumDBs = 11 -- SumLength = 58 | -0.0 | 0.0 |
| LipidClass = PE -- SumLength = 42 | -0.0 | 0.0 | SumDBs = 8 -- SumLength = 54 | 0.0 | 0.0 | LipidClass = PI -- SumLength = 32 | 0.0 | 0.0 | SumDBs = 4 -- SumLength = 48 | -0.0 | 0.0 |
| SumDBs = 7 -- SumLength = 42 | -0.0 | 0.0 | SumDBs = 1 -- SumLength = 46 | 0.0 | 0.0 | LipidClass = PE -- SumDBs = 1 | 0.0 | 0.0 | SumDBs = 6 -- SumLength = 48 | -0.0 | 0.0 |
| SumDBs = 6 -- SumLength = 50 | -0.0 | 0.0 | SumDBs = 11 -- SumLength = 60 | 0.0 | 0.0 | LipidClass = TG -- SumDBs = 0 | 0.0 | 0.0 | SumDBs = 2 -- SumLength = 60 | -0.0 | 0.0 |
| SumDBs = 3 -- SumLength = 58 | -0.0 | 0.0 | SumDBs = 3 -- SumLength = 50 | 0.0 | 0.0 | LipidClass = PE O- -- SumDBs = 3 | 0.0 | 0.0 | LipidClass = LPE O- -- SumLength = 16 | 0.0 | 0.0 |
| SumDBs = 1 -- SumLength = 58 | -0.0 | 0.0 | Headgroup = PI -- SumDBs = 0 | 0.0 | 0.0 | SumDBs = 10 -- SumLength = 58 | 0.0 | 0.0 | LipidClass = LPE O- -- SumDBs = 1 | 0.0 | 0.0 |
| SumDBs = 8 -- SumLength = 42 | -0.0 | 0.0 | SumDBs = 5 -- SumLength = 50 | 0.0 | 0.0 | LipidClass = PE -- SumDBs = 0 | 0.0 | 0.0 | Headgroup = PE -- LipidClass = LPE O- | 0.0 | 0.0 |
| LipidClass = PE -- SumDBs = 0 | -0.0 | 0.0 | SumDBs = 2 -- SumLength = 46 | 0.0 | 0.0 | LipidClass = TG -- SumDBs = 13 | 0.0 | 0.0 | SumDBs = 4 -- SumLength = 60 | 0.0 | 0.0 |
| LipidClass = LPE O- -- SumLength = 16 | 0.0 | 0.0 | LipidClass = PG -- SumDBs = 1 | 0.0 | 0.0 | Headgroup = -- SumDBs = 13 | 0.0 | 0.0 | SumDBs = 3 -- SumLength = 42 | 0.0 | 0.0 |
| LipidClass = LPE O- -- SumDBs = 1 | 0.0 | 0.0 | SumDBs = 0 -- SumLength = 48 | 0.0 | 0.0 | Headgroup = PE -- SumDBs = 8 | 0.0 | 0.0 | SumDBs = 2 -- SumLength = 58 | 0.0 | 0.0 |
| Headgroup = PE -- LipidClass = LPE O- | 0.0 | 0.0 | SumDBs = 7 -- SumLength = 52 | 0.0 | 0.0 | LipidClass = PE -- SumDBs = 8 | 0.0 | 0.0 | SumDBs = 2 -- SumLength = 38 | 0.0 | 0.0 |
| SumDBs = 4 -- SumLength = 46 | 0.0 | 0.0 | LipidClass = TG -- SumDBs = 0 | 0.0 | 0.0 | SumDBs = 4 -- SumLength = 48 | 0.0 | 0.0 | SumDBs = 3 -- SumLength = 60 | 0.0 | 0.0 |
| SumDBs = 7 -- SumLength = 50 | 0.0 | 0.0 | LipidClass = PE -- SumLength = 42 | 0.0 | 0.0 | SumDBs = 11 -- SumLength = 62 | 0.0 | 0.0 | SumDBs = 3 -- SumLength = 58 | 0.0 | 0.0 |
| SumDBs = 2 -- SumLength = 60 | 0.0 | 0.0 | Headgroup = PE -- SumLength = 42 | 0.0 | 0.0 | SumDBs = 13 -- SumLength = 62 | 0.0 | 0.0 | SumDBs = 9 -- SumLength = 56 | 0.0 | 0.0 |
| SumDBs = 6 -- SumLength = 48 | 0.0 | 0.0 | SumDBs = 10 -- SumLength = 60 | 0.0 | 0.0 | SumDBs = 4 -- SumLength = 46 | 0.0 | 0.0 | SumDBs = 13 -- SumLength = 60 | 0.0 | 0.0 |
| SumDBs = 3 -- SumLength = 60 | 0.0 | 0.0 | SumDBs = 7 -- SumLength = 42 | 0.0 | 0.0 | Headgroup = CE -- SumDBs = 5 | 0.0 | 0.0 | SumDBs = 5 -- SumLength = 34 | 0.0 | 0.0 |
| LipidClass = PE O- -- SumLength = 34 | 0.0 | 0.0 | LipidClass = TG -- SumDBs = 9 | 0.0 | 0.0 | LipidClass = CE -- SumDBs = 5 | 0.0 | 0.0 | LipidClass = CE -- SumLength = 22 | 0.0 | 0.0 |
| SumDBs = 4 -- SumLength = 60 | 0.0 | 0.0 | Headgroup = -- SumDBs = 9 | 0.0 | 0.0 | Headgroup = PE -- SumLength = 32 | 0.0 | 0.0 | LipidClass = CE -- SumDBs = 6 | 0.0 | 0.0 |
| SumDBs = 2 -- SumLength = 58 | 0.0 | 0.0 | Headgroup = CE -- SumDBs = 5 | 0.0 | 0.0 | LipidClass = PE -- SumLength = 32 | 0.0 | 0.0 | Headgroup = CE -- SumDBs = 6 | 0.0 | 0.0 |
| SumDBs = 13 -- SumLength = 60 | 0.0 | 0.0 | LipidClass = CE -- SumDBs = 5 | 0.0 | 0.0 | SumDBs = 12 -- SumLength = 62 | 0.0 | 0.0 | Headgroup = CE -- SumLength = 22 | 0.0 | 0.0 |
| LipidClass = TG -- SumDBs = 1 | 0.0 | 0.0 | SumDBs = 5 -- SumLength = 20 | 0.0 | 0.0 | SumDBs = 5 -- SumLength = 20 | 0.0 | 0.0 | SumDBs = 4 -- SumLength = 58 | 0.0 | 0.0 |
| SumDBs = 11 -- SumLength = 58 | 0.0 | 0.0 | SumDBs = 7 -- SumLength = 38 | 0.0 | 0.0 | LipidClass = PE O- -- SumLength = 36 | 0.0 | 0.0 | SumDBs = 8 -- SumLength = 42 | 0.0 | 0.0 |
| SumDBs = 4 -- SumLength = 48 | 0.0 | 0.0 | Headgroup = PI -- SumDBs = 7 | 0.0 | 0.0 | SumDBs = 5 -- SumLength = 50 | 0.0 | 0.0 | LipidClass = TG -- SumDBs = 14 | 0.0 | 0.0 |
| SumDBs = 3 -- SumLength = 20 | 0.0 | 0.0 | LipidClass = PI -- SumDBs = 7 | 0.0 | 0.0 | Headgroup = PC -- SumLength = 32 | 0.0 | 0.0 | SumDBs = 14 -- SumLength = 62 | 0.0 | 0.0 |
| SumDBs = 14 -- SumLength = 62 | 0.0 | 0.0 | LipidClass = PE -- SumLength = 40 | 0.0 | 0.0 | SumDBs = 5 -- SumLength = 48 | 0.0 | 0.0 | Headgroup = -- SumDBs = 14 | 0.0 | 0.0 |
| LipidClass = TG -- SumDBs = 14 | 0.0 | 0.0 | SumDBs = 4 -- SumLength = 34 | 0.0 | 0.0 | Headgroup = -- SumDBs = 9 | 0.0 | 0.0 | LipidClass = PG -- SumDBs = 0 | 0.0 | 0.0 |
| Headgroup = -- SumDBs = 14 | 0.0 | 0.0 | LipidClass = TG -- SumLength = 46 | 0.0 | 0.0 | LipidClass = TG -- SumDBs = 9 | 0.0 | 0.0 | SumDBs = 4 -- SumLength = 56 | 0.0 | 0.0 |
| SumDBs = 5 -- SumLength = 48 | 0.0 | 0.0 | Headgroup = -- SumLength = 46 | 0.0 | 0.0 | SumDBs = 0 -- SumLength = 32 | 0.0 | 0.0 | SumDBs = 10 -- SumLength = 58 | 0.0 | 0.0 |
| SumDBs = 3 -- SumLength = 48 | 0.0 | 0.0 | SumDBs = 4 -- SumLength = 50 | 0.0 | 0.0 | SumDBs = 8 -- SumLength = 54 | 0.0 | 0.0 | LipidClass = TG -- SumDBs = 11 | 0.0 | 0.0 |
| LipidClass = PC -- SumLength = 32 | 0.0 | 0.0 | SumDBs = 1 -- SumLength = 48 | 0.0 | 0.0 | Headgroup = -- SumLength = 46 | 0.0 | 0.0 | Headgroup = -- SumDBs = 11 | 0.0 | 0.0 |
| SumDBs = 11 -- SumLength = 62 | 0.0 | 0.0 | SumDBs = 4 -- SumLength = 36 | 0.0 | 0.0 | LipidClass = TG -- SumLength = 46 | 0.0 | 0.0 | SumDBs = 10 -- SumLength = 60 | 0.0 | 0.0 |
| SumDBs = 13 -- SumLength = 62 | 0.0 | 0.0 | SumDBs = 7 -- SumLength = 54 | 0.0 | 0.0 | SumDBs = 9 -- SumLength = 56 | 0.0 | 0.0 | SumDBs = 11 -- SumLength = 62 | 0.0 | 0.0 |
| SumDBs = 12 -- SumLength = 62 | 0.0 | 0.0 | LipidClass = Cer -- SumLength = 34 | 0.0 | 0.0 | SumDBs = 7 -- SumLength = 52 | 0.0 | 0.0 | SumDBs = 13 -- SumLength = 62 | 0.0 | 0.0 |
| Headgroup = -- SumDBs = 9 | 0.0 | 0.0 | SumDBs = 7 -- SumLength = 60 | 0.0 | 0.0 | LipidClass = PC -- SumLength = 40 | 0.0 | 0.0 | SumDBs = 12 -- SumLength = 62 | 0.0 | 0.0 |
| LipidClass = TG -- SumDBs = 9 | 0.0 | 0.0 | SumDBs = 1 -- SumLength = 38 | 0.0 | 0.0 | LipidClass = TG -- SumLength = 56 | 0.0 | 0.0 | Headgroup = -- SumDBs = 13 | 0.0 | 0.0 |
| LipidClass = TG -- SumDBs = 13 | 0.0 | 0.0 | LipidClass = PE O- -- SumDBs = 3 | 0.0 | 0.0 | Headgroup = -- SumLength = 56 | 0.0 | 0.0 | LipidClass = TG -- SumDBs = 13 | 0.0 | 0.0 |
| Headgroup = -- SumDBs = 13 | 0.0 | 0.0 | Headgroup = -- SumLength = 36 | 0.0 | 0.0 | SumDBs = 4 -- SumLength = 34 | 0.0 | 0.0 | LipidClass = TG -- SumDBs = 10 | 0.0 | 0.0 |
| SumDBs = 5 -- SumLength = 34 | 0.0 | 0.0 | LipidClass = DG -- SumLength = 36 | 0.0 | 0.0 | LipidClass = PC -- SumDBs = 2 | 0.0 | 0.0 | Headgroup = -- SumDBs = 10 | 0.0 | 0.0 |
| SumDBs = 2 -- SumLength = 38 | 0.0 | 0.0 | SumDBs = 2 -- SumLength = 48 | 0.0 | 0.0 | LipidClass = PG -- SumLength = 36 | 0.0 | 0.0 | LipidClass = PE -- SumDBs = 5 | 0.0 | 0.0 |
| SumDBs = 6 -- SumLength = 54 | 0.0 | 0.0 | LipidClass = TG -- SumLength = 48 | 0.0 | 0.0 | Headgroup = PG -- SumLength = 36 | 0.0 | 0.0 | Headgroup = -- SumLength = 60 | 0.0 | 0.0 |
| SumDBs = 11 -- SumLength = 60 | 0.0 | 0.0 | Headgroup = -- SumLength = 48 | 0.0 | 0.0 | SumDBs = 3 -- SumLength = 38 | 0.0 | 0.0 | LipidClass = TG -- SumLength = 60 | 0.0 | 0.0 |
| SumDBs = 1 -- SumLength = 50 | 0.0 | 0.0 | LipidClass = PE O- -- SumLength = 36 | 0.0 | 0.0 | SumDBs = 2 -- SumLength = 54 | 0.0 | 0.0 | LipidClass = TG -- SumDBs = 8 | 0.0 | 0.0 |
| Headgroup = -- LipidClass = Cer | 0.0 | 0.0 | SumDBs = 8 -- SumLength = 60 | 0.0 | 0.0 | SumDBs = 3 -- SumLength = 48 | 0.0 | 0.0 | Headgroup = -- SumDBs = 8 | 0.0 | 0.0 |
| SumDBs = 1 -- SumLength = 16 | 0.0 | 0.0 | LipidClass = PE -- SumDBs = 1 | 0.0 | 0.0 | SumDBs = 6 -- SumLength = 56 | 0.0 | 0.0 | LipidClass = PC -- SumLength = 40 | 0.0 | 0.0 |
| Headgroup = -- LipidClass = TG | 0.0 | 0.0 | Headgroup = PG -- SumLength = 20 | 0.0 | 0.0 | LipidClass = DG -- SumDBs = 0 | 0.0 | 0.0 | LipidClass = PC O- -- SumDBs = 5 | 0.0 | 0.0 |
| Headgroup = PI -- LipidClass = PI | 0.0 | 0.0 | LipidClass = LPG -- SumDBs = 4 | 0.0 | 0.0 | SumDBs = 9 -- SumLength = 58 | 0.0 | 0.0 | SumDBs = 3 -- SumLength = 56 | 0.0 | 0.0 |
| SumDBs = 0 -- SumLength = 38 | 0.0 | 0.0 | LipidClass = LPG -- SumLength = 20 | 0.0 | 0.0 | LipidClass = PI -- SumDBs = 7 | 0.0 | 0.0 | LipidClass = TG -- SumLength = 52 | 0.0 | 0.0 |
| Headgroup = PS -- SumDBs = 0 | 0.0 | 0.0 | SumDBs = 4 -- SumLength = 56 | 0.0 | 0.0 | Headgroup = PI -- SumDBs = 7 | 0.0 | 0.0 | Headgroup = -- SumLength = 52 | 0.0 | 0.0 |
| LipidClass = PS -- SumDBs = 0 | 0.0 | 0.0 | LipidClass = Cer -- SumDBs = 3 | 0.0 | 0.0 | SumDBs = 1 -- SumLength = 33 | 0.0 | 0.0 | Headgroup = -- LipidClass = Cer | 0.0 | 0.0 |
| SumDBs = 10 -- SumLength = 60 | 0.0 | 0.0 | SumDBs = 1 -- SumLength = 16 | 0.0 | 0.0 | LipidClass = SM -- SumLength = 33 | 0.0 | 0.0 | Headgroup = -- LipidClass = TG | 0.0 | 0.0 |
| Headgroup = PC -- LipidClass = PC O- | 0.0 | 0.0 | LipidClass = PE -- SumDBs = 8 | 0.0 | 0.0 | Headgroup = PC -- SumLength = 33 | 0.0 | 0.0 | Headgroup = PI -- LipidClass = PI | 0.0 | 0.0 |
| Headgroup = PC -- LipidClass = LPC | 0.0 | 0.0 | Headgroup = PE -- SumDBs = 8 | 0.0 | 0.0 | SumDBs = 6 -- SumLength = 38 | 0.0 | 0.0 | SumDBs = 5 -- SumLength = 56 | 0.0 | 0.0 |
| Headgroup = PG -- LipidClass = LPG | 0.0 | 0.0 | LipidClass = PG -- SumDBs = 4 | 0.0 | 0.0 | Headgroup = -- SumLength = 52 | 0.0 | 0.0 | LipidClass = TG -- SumLength = 62 | 0.0 | 0.0 |
| LipidClass = PE O- -- SumDBs = 3 | 0.0 | 0.0 | Headgroup = PE -- SumLength = 22 | 0.0 | 0.0 | LipidClass = TG -- SumLength = 52 | 0.0 | 0.0 | Headgroup = -- SumLength = 62 | 0.0 | 0.0 |
| Headgroup = CE -- LipidClass = CE | 0.0 | 0.0 | LipidClass = LPE -- SumLength = 22 | 0.0 | 0.0 | SumDBs = 3 -- SumLength = 52 | 0.0 | 0.0 | LipidClass = CE -- SumDBs = 2 | 0.0 | 0.0 |
| Headgroup = -- LipidClass = DG | 0.0 | 0.0 | LipidClass = LPE -- SumDBs = 6 | 0.0 | 0.0 | SumDBs = 3 -- SumLength = 42 | 0.0 | 0.0 | Headgroup = CE -- SumDBs = 2 | 0.0 | 0.0 |
| Headgroup = -- SumDBs = 11 | 0.0 | 0.0 | LipidClass = SM -- SumLength = 33 | 0.0 | 0.0 | SumDBs = 4 -- SumLength = 54 | 0.0 | 0.0 | Headgroup = PS -- SumDBs = 0 | 0.0 | 0.0 |
| LipidClass = TG -- SumDBs = 11 | 0.0 | 0.0 | SumDBs = 1 -- SumLength = 33 | 0.0 | 0.0 | Headgroup = -- SumDBs = 0 | 0.0 | 0.0 | SumDBs = 0 -- SumLength = 38 | 0.0 | 0.0 |
| Headgroup = PC -- LipidClass = PC | 0.0 | 0.0 | Headgroup = PC -- SumLength = 33 | 0.0 | 0.0 | LipidClass = TG -- SumLength = 62 | 0.0 | 0.0 | LipidClass = PS -- SumDBs = 0 | 0.0 | 0.0 |
| SumDBs = 10 -- SumLength = 58 | 0.0 | 0.0 | SumDBs = 3 -- SumLength = 38 | 0.0 | 0.0 | Headgroup = -- SumLength = 62 | 0.0 | 0.0 | Headgroup = PC -- LipidClass = PC O- | 0.0 | 0.0 |
| LipidClass = CE -- SumLength = 20 | 0.0 | 0.0 | SumDBs = 6 -- SumLength = 58 | 0.0 | 0.0 | SumDBs = 8 -- SumLength = 58 | 0.0 | 0.0 | Headgroup = PC -- LipidClass = LPC | 0.0 | 0.0 |
| Headgroup = CE -- SumLength = 20 | 0.0 | 0.0 | SumDBs = 9 -- SumLength = 60 | 0.0 | 0.0 | LipidClass = TG -- SumDBs = 5 | 0.0 | 0.0 | Headgroup = PG -- LipidClass = LPG | 0.0 | 0.0 |
| Headgroup = PG -- LipidClass = PG | 0.0 | 0.0 | SumDBs = 5 -- SumLength = 36 | 0.0 | 0.0 | SumDBs = 4 -- SumLength = 40 | 0.0 | 0.0 | Headgroup = CE -- LipidClass = CE | 0.0 | 0.0 |
| LipidClass = TG -- SumLength = 62 | 0.0 | 0.0 | SumDBs = 5 -- SumLength = 56 | 0.0 | 0.0 | LipidClass = PE -- SumDBs = 7 | 0.0 | 0.0 | Headgroup = -- LipidClass = DG | 0.0 | 0.0 |
| Headgroup = -- SumLength = 62 | 0.0 | 0.0 | LipidClass = PC -- SumDBs = 7 | 0.0 | 0.0 | SumDBs = 6 -- SumLength = 36 | 0.0 | 0.0 | LipidClass = LPI -- SumDBs = 0 | 0.0 | 0.0 |
| LipidClass = PE -- SumDBs = 5 | 0.0 | 0.0 | LipidClass = PG -- SumDBs = 3 | 0.0 | 0.0 | SumDBs = 8 -- SumLength = 56 | 0.0 | 0.0 | Headgroup = PI -- SumDBs = 0 | 0.0 | 0.0 |
| Headgroup = P -- LipidClass = PA | 0.0 | 0.0 | SumDBs = 7 -- SumLength = 58 | 0.0 | 0.0 | SumDBs = 4 -- SumLength = 50 | 0.0 | 0.0 | LipidClass = PE O- -- SumLength = 36 | 0.0 | 0.0 |
| Headgroup = -- SumLength = 52 | 0.0 | 0.0 | LipidClass = PE -- SumDBs = 3 | 0.0 | 0.0 | LipidClass = PC -- SumLength = 32 | 0.0 | 0.0 | Headgroup = PC -- LipidClass = PC | 0.0 | 0.0 |
| LipidClass = TG -- SumLength = 52 | 0.0 | 0.0 | SumDBs = 2 -- SumLength = 40 | 0.0 | 0.0 | SumDBs = 7 -- SumLength = 54 | 0.0 | 0.0 | Headgroup = PG -- LipidClass = PG | 0.0 | 0.0 |
| LipidClass = PC -- SumDBs = 0 | 0.0 | 0.0 | SumDBs = 8 -- SumLength = 58 | 0.0 | 0.0 | LipidClass = LPG -- SumLength = 18 | 0.0 | 0.0 | LipidClass = SM -- SumLength = 42 | 0.0 | 0.0 |
| Headgroup = PC -- LipidClass = SM | 0.0 | 0.0 | LipidClass = PG -- SumDBs = 6 | 0.0 | 0.0 | Headgroup = PG -- SumLength = 18 | 0.0 | 0.0 | Headgroup = PC -- SumLength = 42 | 0.0 | 0.0 |
| Headgroup = PE -- LipidClass = LPE | 0.0 | 0.0 | LipidClass = PC -- SumDBs = 0 | 0.0 | 0.0 | LipidClass = LPE -- SumLength = 22 | 0.0 | 0.0 | Headgroup = P -- LipidClass = PA | 0.0 | 0.0 |
| Headgroup = PS -- LipidClass = PS | 0.0 | 0.0 | LipidClass = PE O- -- SumLength = 34 | 0.001 | 0.001 | Headgroup = PE -- SumLength = 22 | 0.0 | 0.0 | Headgroup = PG -- SumLength = 36 | 0.0 | 0.0 |
| LipidClass = PE -- SumDBs = 1 | 0.0 | 0.0 | LipidClass = DG -- SumDBs = 1 | 0.001 | 0.001 | LipidClass = LPE -- SumDBs = 6 | 0.0 | 0.0 | LipidClass = PG -- SumLength = 36 | 0.0 | 0.0 |
| Headgroup = PE -- LipidClass = PE O- | 0.0 | 0.0 | SumDBs = 2 -- SumLength = 54 | 0.001 | 0.001 | LipidClass = PE -- SumDBs = 3 | 0.0 | 0.0 | Headgroup = PC -- LipidClass = SM | 0.0 | 0.0 |
| Headgroup = PC -- SumLength = 41 | 0.0 | 0.0 | LipidClass = PE O- -- SumDBs = 2 | 0.001 | 0.001 | LipidClass = DG -- SumDBs = 5 | 0.0 | 0.0 | Headgroup = PE -- LipidClass = LPE | 0.0 | 0.0 |
| LipidClass = SM -- SumLength = 41 | 0.0 | 0.0 | SumDBs = 1 -- SumLength = 50 | 0.001 | 0.001 | Headgroup = PC -- SumLength = 42 | 0.0 | 0.0 | Headgroup = PS -- LipidClass = PS | 0.0 | 0.0 |
| SumDBs = 3 -- SumLength = 56 | 0.0 | 0.0 | Headgroup = CE -- SumLength = 22 | 0.001 | 0.001 | LipidClass = SM -- SumLength = 42 | 0.0 | 0.0 | LipidClass = TG -- SumDBs = 1 | 0.0 | 0.0 |
| SumDBs = 9 -- SumLength = 58 | 0.0 | 0.0 | Headgroup = CE -- SumDBs = 6 | 0.001 | 0.001 | SumDBs = 6 -- SumLength = 52 | 0.0 | 0.0 | Headgroup = PE -- LipidClass = PE O- | 0.0 | 0.0 |
| SumDBs = 12 -- SumLength = 60 | 0.0 | 0.0 | LipidClass = CE -- SumDBs = 6 | 0.001 | 0.001 | SumDBs = 2 -- SumLength = 32 | 0.0 | 0.0 | LipidClass = LPG -- SumLength = 18 | 0.0 | 0.0 |
| Headgroup = PE -- LipidClass = PE | 0.0 | 0.0 | LipidClass = CE -- SumLength = 22 | 0.001 | 0.001 | Headgroup = PE -- SumDBs = 3 | 0.0 | 0.0 | Headgroup = PG -- SumLength = 18 | 0.0 | 0.0 |
| LipidClass = DG -- SumDBs = 4 | 0.0 | 0.0 | Headgroup = -- SumLength = 60 | 0.001 | 0.001 | Headgroup = -- SumDBs = 3 | 0.0 | 0.0 | LipidClass = SM -- SumLength = 34 | 0.0 | 0.0 |
| LipidClass = Cer -- SumLength = 41 | 0.0 | 0.0 | LipidClass = TG -- SumLength = 60 | 0.001 | 0.001 | LipidClass = PE -- SumDBs = 6 | 0.0 | 0.0 | Headgroup = PG -- SumDBs = 1 | 0.0 | 0.0 |
| Headgroup = -- SumLength = 41 | 0.0 | 0.0 | SumDBs = 6 -- SumLength = 52 | 0.001 | 0.001 | LipidClass = SM -- SumDBs = 3 | 0.0 | 0.0 | Headgroup = PG -- SumLength = 20 | 0.0 | 0.0 |
| Headgroup = -- SumDBs = 10 | 0.0 | 0.0 | LipidClass = SM -- SumLength = 36 | 0.001 | 0.001 | LipidClass = PC -- SumDBs = 6 | 0.0 | 0.0 | LipidClass = LPG -- SumLength = 20 | 0.0 | 0.0 |
| LipidClass = TG -- SumDBs = 10 | 0.0 | 0.0 | SumDBs = 0 -- SumLength = 34 | 0.001 | 0.001 | LipidClass = PC -- SumDBs = 7 | 0.0 | 0.0 | LipidClass = LPG -- SumDBs = 4 | 0.0 | 0.0 |
| Headgroup = PI -- SumDBs = 0 | 0.0 | 0.0 | LipidClass = PE -- SumDBs = 2 | 0.001 | 0.001 | LipidClass = LPC -- SumDBs = 1 | 0.0 | 0.0 | LipidClass = PG -- SumDBs = 4 | 0.0 | 0.0 |
| Headgroup = -- SumLength = 36 | 0.0 | 0.0 | LipidClass = TG -- SumDBs = 5 | 0.001 | 0.001 | Headgroup = -- SumDBs = 8 | 0.0 | 0.0 | SumDBs = 12 -- SumLength = 60 | 0.0 | 0.0 |
| LipidClass = DG -- SumLength = 36 | 0.0 | 0.0 | Headgroup = PG -- SumDBs = 4 | 0.001 | 0.001 | LipidClass = TG -- SumDBs = 8 | 0.0 | 0.0 | SumDBs = 4 -- SumLength = 40 | 0.0 | 0.0 |
| Headgroup = CE -- SumDBs = 4 | 0.0 | 0.0 | Headgroup = PE -- SumDBs = 3 | 0.001 | 0.001 | Headgroup = CE -- SumDBs = 1 | 0.0 | 0.0 | Headgroup = PE -- LipidClass = PE | 0.001 | 0.001 |
| LipidClass = CE -- SumDBs = 4 | 0.0 | 0.0 | LipidClass = PE O- -- SumDBs = 7 | 0.001 | 0.001 | LipidClass = CE -- SumDBs = 1 | 0.0 | 0.0 | SumDBs = 1 -- SumLength = 50 | 0.001 | 0.001 |
| LipidClass = TG -- SumLength = 60 | 0.0 | 0.0 | LipidClass = LPC -- SumDBs = 2 | 0.001 | 0.001 | Headgroup = PG -- SumDBs = 1 | 0.0 | 0.0 | SumDBs = 7 -- SumLength = 58 | 0.001 | 0.001 |
| Headgroup = -- SumLength = 60 | 0.0 | 0.0 | SumDBs = 2 -- SumLength = 32 | 0.001 | 0.001 | LipidClass = TG -- SumDBs = 7 | 0.0 | 0.0 | LipidClass = TG -- SumLength = 54 | 0.001 | 0.001 |
| LipidClass = TG -- SumDBs = 12 | 0.0 | 0.0 | LipidClass = TG -- SumDBs = 1 | 0.001 | 0.001 | LipidClass = PE O- -- SumDBs = 2 | 0.0 | 0.0 | Headgroup = -- SumLength = 54 | 0.001 | 0.001 |
| Headgroup = -- SumDBs = 12 | 0.0 | 0.0 | LipidClass = TG -- SumDBs = 7 | 0.001 | 0.001 | Headgroup = -- SumDBs = 5 | 0.001 | 0.001 | LipidClass = DG -- SumDBs = 1 | 0.001 | 0.001 |
| LipidClass = PS -- SumDBs = 5 | 0.0 | 0.0 | Headgroup = -- SumLength = 58 | 0.001 | 0.001 | LipidClass = PE -- SumLength = 36 | 0.001 | 0.001 | SumDBs = 3 -- SumLength = 20 | 0.001 | 0.001 |
| Headgroup = PS -- SumDBs = 5 | 0.0 | 0.0 | LipidClass = TG -- SumLength = 58 | 0.001 | 0.001 | LipidClass = PS -- SumDBs = 4 | 0.001 | 0.001 | Headgroup = -- SumDBs = 12 | 0.001 | 0.001 |
| SumDBs = 2 -- SumLength = 40 | 0.0 | 0.0 | LipidClass = PC -- SumLength = 32 | 0.001 | 0.001 | Headgroup = PS -- SumDBs = 4 | 0.001 | 0.001 | LipidClass = TG -- SumDBs = 12 | 0.001 | 0.001 |
| LipidClass = LPI -- SumDBs = 0 | 0.0 | 0.0 | LipidClass = PE -- SumLength = 36 | 0.001 | 0.001 | Headgroup = PS -- SumDBs = 2 | 0.001 | 0.001 | Headgroup = CE -- SumDBs = 4 | 0.001 | 0.001 |
| LipidClass = DG -- SumLength = 34 | 0.0 | 0.0 | Headgroup = -- SumLength = 41 | 0.001 | 0.001 | LipidClass = PS -- SumDBs = 2 | 0.001 | 0.001 | LipidClass = PE -- SumLength = 40 | 0.001 | 0.001 |
| LipidClass = PC -- SumLength = 40 | 0.0 | 0.0 | LipidClass = Cer -- SumLength = 41 | 0.001 | 0.001 | LipidClass = PS -- SumDBs = 5 | 0.001 | 0.001 | LipidClass = CE -- SumDBs = 4 | 0.001 | 0.001 |
| SumDBs = 4 -- SumLength = 40 | 0.0 | 0.0 | LipidClass = PC -- SumDBs = 5 | 0.001 | 0.001 | Headgroup = PS -- SumDBs = 5 | 0.001 | 0.001 | Headgroup = -- SumDBs = 7 | 0.001 | 0.001 |
| Headgroup = -- SumDBs = 8 | 0.0 | 0.0 | SumDBs = 6 -- SumLength = 56 | 0.001 | 0.001 | LipidClass = PE O- -- SumLength = 34 | 0.001 | 0.001 | LipidClass = PS -- SumDBs = 1 | 0.001 | 0.001 |
| LipidClass = TG -- SumDBs = 8 | 0.0 | 0.0 | SumDBs = 4 -- SumLength = 54 | 0.001 | 0.001 | Headgroup = PC -- SumDBs = 3 | 0.001 | 0.001 | Headgroup = PS -- SumDBs = 1 | 0.001 | 0.001 |
| LipidClass = LPG -- SumLength = 20 | 0.0 | 0.0 | LipidClass = LPC -- SumLength = 20 | 0.001 | 0.001 | Headgroup = PE -- SumLength = 36 | 0.001 | 0.001 | SumDBs = 4 -- SumLength = 20 | 0.001 | 0.001 |
| LipidClass = LPG -- SumDBs = 4 | 0.0 | 0.0 | LipidClass = LPC -- SumDBs = 4 | 0.001 | 0.001 | Headgroup = PI -- SumLength = 18 | 0.001 | 0.001 | LipidClass = Cer -- SumDBs = 3 | 0.001 | 0.001 |
| Headgroup = PG -- SumLength = 20 | 0.0 | 0.0 | Headgroup = PC -- SumLength = 20 | 0.001 | 0.001 | LipidClass = LPI -- SumLength = 18 | 0.001 | 0.001 | LipidClass = CE -- SumDBs = 3 | 0.001 | 0.001 |
| LipidClass = CE -- SumDBs = 2 | 0.0 | 0.0 | LipidClass = LPG -- SumDBs = 3 | 0.001 | 0.001 | SumDBs = 7 -- SumLength = 56 | 0.001 | 0.001 | Headgroup = CE -- SumDBs = 3 | 0.001 | 0.001 |
| LipidClass = DG -- SumLength = 38 | 0.0 | 0.0 | LipidClass = PG -- SumLength = 32 | 0.001 | 0.001 | SumDBs = 1 -- SumLength = 38 | 0.001 | 0.001 | Headgroup = -- SumLength = 56 | 0.001 | 0.001 |
| Headgroup = CE -- SumDBs = 2 | 0.0 | 0.0 | Headgroup = PG -- SumLength = 32 | 0.001 | 0.001 | Headgroup = PE -- SumLength = 18 | 0.001 | 0.001 | LipidClass = TG -- SumLength = 56 | 0.001 | 0.001 |
| LipidClass = CE -- SumLength = 18 | 0.0 | 0.0 | LipidClass = DG -- SumDBs = 0 | 0.001 | 0.001 | LipidClass = LPE -- SumLength = 18 | 0.001 | 0.001 | Headgroup = PG -- SumDBs = 6 | 0.001 | 0.001 |
| Headgroup = CE -- SumLength = 18 | 0.0 | 0.0 | LipidClass = PE -- SumDBs = 7 | 0.001 | 0.001 | Headgroup = PC -- SumDBs = 2 | 0.001 | 0.001 | Headgroup = PC -- SumDBs = 6 | 0.001 | 0.001 |
| LipidClass = PE O- -- SumDBs = 4 | 0.0 | 0.0 | Headgroup = PE -- SumLength = 36 | 0.001 | 0.001 | Headgroup = PG -- SumDBs = 6 | 0.001 | 0.001 | SumDBs = 5 -- SumLength = 54 | 0.001 | 0.001 |
| LipidClass = DG -- SumDBs = 7 | 0.0 | 0.0 | Headgroup = PE -- SumDBs = 2 | 0.001 | 0.001 | SumDBs = 6 -- SumLength = 54 | 0.001 | 0.001 | LipidClass = PC O- -- SumLength = 40 | 0.001 | 0.001 |
| SumDBs = 3 -- SumLength = 40 | 0.0 | 0.0 | Headgroup = -- SumDBs = 0 | 0.001 | 0.001 | Headgroup = PG -- SumDBs = 3 | 0.001 | 0.001 | LipidClass = PG -- SumDBs = 3 | 0.001 | 0.001 |
| Headgroup = PE -- SumLength = 16 | 0.0 | 0.0 | LipidClass = PE O- -- SumDBs = 6 | 0.001 | 0.001 | Headgroup = P -- SumDBs = 1 | 0.001 | 0.001 | LipidClass = PC O- -- SumDBs = 6 | 0.001 | 0.001 |
| LipidClass = LPE -- SumLength = 16 | 0.0 | 0.0 | SumDBs = 3 -- SumLength = 18 | 0.001 | 0.001 | LipidClass = PA -- SumDBs = 1 | 0.001 | 0.001 | SumDBs = 3 -- SumLength = 52 | 0.001 | 0.001 |
| SumDBs = 8 -- SumLength = 56 | 0.0 | 0.0 | Headgroup = -- SumLength = 38 | 0.001 | 0.001 | LipidClass = PC -- SumDBs = 4 | 0.001 | 0.001 | LipidClass = DG -- SumLength = 38 | 0.001 | 0.001 |
| Headgroup = PS -- SumDBs = 3 | 0.001 | 0.001 | SumDBs = 5 -- SumLength = 52 | 0.001 | 0.001 | Headgroup = -- SumLength = 32 | 0.001 | 0.001 | LipidClass = SM -- SumLength = 40 | 0.001 | 0.001 |
| LipidClass = PS -- SumDBs = 3 | 0.001 | 0.001 | Headgroup = PS -- SumDBs = 2 | 0.001 | 0.001 | LipidClass = DG -- SumLength = 32 | 0.001 | 0.001 | Headgroup = PG -- SumDBs = 4 | 0.001 | 0.001 |
| SumDBs = 3 -- SumLength = 52 | 0.001 | 0.001 | LipidClass = PS -- SumDBs = 2 | 0.001 | 0.001 | LipidClass = PE -- SumDBs = 2 | 0.001 | 0.001 | LipidClass = LPG -- SumLength = 22 | 0.001 | 0.001 |
| LipidClass = SM -- SumLength = 38 | 0.001 | 0.001 | LipidClass = PE -- SumLength = 34 | 0.001 | 0.001 | Headgroup = -- SumLength = 42 | 0.001 | 0.001 | LipidClass = LPG -- SumDBs = 6 | 0.001 | 0.001 |
| LipidClass = SM -- SumLength = 40 | 0.001 | 0.001 | LipidClass = SM -- SumDBs = 3 | 0.001 | 0.001 | LipidClass = Cer -- SumLength = 42 | 0.001 | 0.001 | Headgroup = PG -- SumLength = 22 | 0.001 | 0.001 |
| Headgroup = PI -- SumDBs = 3 | 0.001 | 0.001 | Headgroup = PG -- SumLength = 16 | 0.001 | 0.001 | SumDBs = 3 -- SumLength = 50 | 0.001 | 0.001 | LipidClass = DG -- SumDBs = 7 | 0.001 | 0.001 |
| LipidClass = PI -- SumDBs = 3 | 0.001 | 0.001 | LipidClass = LPG -- SumDBs = 0 | 0.001 | 0.001 | SumDBs = 5 -- SumLength = 52 | 0.001 | 0.001 | SumDBs = 8 -- SumLength = 56 | 0.001 | 0.001 |
| LipidClass = LPG -- SumDBs = 6 | 0.001 | 0.001 | LipidClass = LPG -- SumLength = 16 | 0.001 | 0.001 | Headgroup = PG -- SumDBs = 0 | 0.001 | 0.001 | LipidClass = DG -- SumDBs = 6 | 0.001 | 0.001 |
| LipidClass = LPG -- SumLength = 22 | 0.001 | 0.001 | Headgroup = PG -- SumDBs = 0 | 0.001 | 0.001 | Headgroup = PC -- SumLength = 22 | 0.001 | 0.001 | Headgroup = -- SumDBs = 2 | 0.001 | 0.001 |
| Headgroup = PG -- SumLength = 22 | 0.001 | 0.001 | LipidClass = DG -- SumDBs = 5 | 0.001 | 0.001 | LipidClass = LPC -- SumDBs = 6 | 0.001 | 0.001 | LipidClass = CE -- SumLength = 20 | 0.001 | 0.001 |
| LipidClass = CE -- SumLength = 22 | 0.001 | 0.001 | Headgroup = PG -- SumDBs = 3 | 0.001 | 0.001 | LipidClass = LPC -- SumLength = 22 | 0.001 | 0.001 | Headgroup = CE -- SumLength = 20 | 0.001 | 0.001 |
| LipidClass = CE -- SumDBs = 6 | 0.001 | 0.001 | LipidClass = PC O- -- SumLength = 38 | 0.002 | 0.002 | SumDBs = 1 -- SumLength = 34 | 0.001 | 0.001 | LipidClass = PS -- SumLength = 38 | 0.001 | 0.001 |
| Headgroup = CE -- SumDBs = 6 | 0.001 | 0.001 | Headgroup = -- SumLength = 42 | 0.002 | 0.002 | LipidClass = LPG -- SumLength = 16 | 0.001 | 0.001 | Headgroup = PS -- SumLength = 38 | 0.001 | 0.001 |
| Headgroup = CE -- SumLength = 22 | 0.001 | 0.001 | LipidClass = Cer -- SumLength = 42 | 0.002 | 0.002 | LipidClass = LPG -- SumDBs = 0 | 0.001 | 0.001 | Headgroup = PE -- SumDBs = 0 | 0.001 | 0.001 |
| SumDBs = 5 -- SumLength = 54 | 0.001 | 0.001 | LipidClass = DG -- SumDBs = 3 | 0.002 | 0.002 | Headgroup = PG -- SumLength = 16 | 0.001 | 0.001 | LipidClass = PC O- -- SumLength = 36 | 0.001 | 0.001 |
| LipidClass = PC O- -- SumDBs = 1 | 0.001 | 0.001 | SumDBs = 3 -- SumLength = 42 | 0.002 | 0.002 | Headgroup = PI -- SumLength = 40 | 0.001 | 0.001 | SumDBs = 3 -- SumLength = 40 | 0.001 | 0.001 |
| LipidClass = PC O- -- SumLength = 34 | 0.001 | 0.001 | SumDBs = 1 -- SumLength = 34 | 0.002 | 0.002 | LipidClass = PI -- SumLength = 40 | 0.001 | 0.001 | LipidClass = PS -- SumDBs = 3 | 0.001 | 0.001 |
| LipidClass = PC -- SumDBs = 3 | 0.001 | 0.001 | LipidClass = PS -- SumLength = 36 | 0.002 | 0.002 | LipidClass = SM -- SumLength = 36 | 0.001 | 0.001 | Headgroup = PS -- SumDBs = 3 | 0.001 | 0.001 |
| LipidClass = LPG -- SumDBs = 1 | 0.001 | 0.001 | Headgroup = PS -- SumLength = 36 | 0.002 | 0.002 | Headgroup = PG -- SumLength = 32 | 0.001 | 0.001 | LipidClass = DG -- SumLength = 34 | 0.001 | 0.001 |
| SumDBs = 0 -- SumLength = 18 | 0.001 | 0.001 | SumDBs = 1 -- SumLength = 42 | 0.002 | 0.002 | LipidClass = PG -- SumLength = 32 | 0.001 | 0.001 | LipidClass = SM -- SumLength = 38 | 0.001 | 0.001 |
| Headgroup = PC -- SumDBs = 0 | 0.001 | 0.001 | LipidClass = PI -- SumDBs = 4 | 0.002 | 0.002 | Headgroup = -- SumDBs = 6 | 0.001 | 0.001 | LipidClass = PC O- -- SumDBs = 2 | 0.001 | 0.001 |
| LipidClass = DG -- SumDBs = 1 | 0.001 | 0.001 | LipidClass = PC O- -- SumDBs = 5 | 0.002 | 0.002 | LipidClass = PG -- SumDBs = 6 | 0.001 | 0.001 | SumDBs = 2 -- SumLength = 42 | 0.001 | 0.001 |
| Headgroup = PG -- SumDBs = 2 | 0.001 | 0.001 | LipidClass = PE O- -- SumLength = 40 | 0.002 | 0.002 | SumDBs = 4 -- SumLength = 36 | 0.001 | 0.001 | LipidClass = LPE -- SumDBs = 0 | 0.001 | 0.001 |
| LipidClass = PS -- SumDBs = 1 | 0.001 | 0.001 | LipidClass = TG -- SumDBs = 6 | 0.002 | 0.002 | LipidClass = TG -- SumDBs = 6 | 0.001 | 0.001 | LipidClass = PE O- -- SumDBs = 4 | 0.001 | 0.001 |
| Headgroup = PS -- SumDBs = 1 | 0.001 | 0.001 | Headgroup = PE -- SumDBs = 7 | 0.002 | 0.002 | LipidClass = DG -- SumDBs = 4 | 0.001 | 0.001 | LipidClass = LPC -- SumLength = 18 | 0.001 | 0.001 |
| Headgroup = PC -- SumDBs = 4 | 0.001 | 0.001 | Headgroup = -- SumLength = 32 | 0.002 | 0.002 | Headgroup = PE -- SumLength = 40 | 0.001 | 0.001 | Headgroup = PC -- SumLength = 18 | 0.001 | 0.001 |
| LipidClass = PC O- -- SumLength = 36 | 0.001 | 0.001 | LipidClass = DG -- SumLength = 32 | 0.002 | 0.002 | Headgroup = PG -- SumLength = 34 | 0.001 | 0.001 | Headgroup = -- SumDBs = 3 | 0.001 | 0.001 |
| Headgroup = PC -- SumLength = 42 | 0.001 | 0.001 | Headgroup = PE -- SumLength = 34 | 0.002 | 0.002 | LipidClass = PG -- SumLength = 34 | 0.001 | 0.001 | Headgroup = PI -- SumDBs = 4 | 0.001 | 0.001 |
| LipidClass = SM -- SumLength = 42 | 0.001 | 0.001 | LipidClass = PA -- SumDBs = 2 | 0.002 | 0.002 | Headgroup = PI -- SumDBs = 6 | 0.001 | 0.001 | LipidClass = LPE -- SumLength = 16 | 0.001 | 0.001 |
| LipidClass = PC -- SumDBs = 6 | 0.001 | 0.001 | Headgroup = P -- SumDBs = 2 | 0.002 | 0.002 | LipidClass = PI -- SumDBs = 6 | 0.001 | 0.001 | Headgroup = PE -- SumLength = 16 | 0.001 | 0.001 |
| Headgroup = PC -- SumLength = 38 | 0.001 | 0.001 | Headgroup = CE -- SumLength = 16 | 0.002 | 0.002 | LipidClass = PC O- -- SumDBs = 3 | 0.001 | 0.001 | LipidClass = PC O- -- SumDBs = 1 | 0.001 | 0.001 |
| LipidClass = DG -- SumDBs = 2 | 0.001 | 0.001 | LipidClass = CE -- SumDBs = 0 | 0.002 | 0.002 | LipidClass = PE O- -- SumLength = 40 | 0.001 | 0.001 | LipidClass = PC O- -- SumLength = 34 | 0.001 | 0.001 |
| SumDBs = 4 -- SumLength = 20 | 0.001 | 0.001 | LipidClass = CE -- SumLength = 16 | 0.002 | 0.002 | LipidClass = LPG -- SumDBs = 2 | 0.001 | 0.001 | LipidClass = DG -- SumDBs = 2 | 0.002 | 0.002 |
| LipidClass = DG -- SumDBs = 6 | 0.001 | 0.001 | Headgroup = CE -- SumDBs = 0 | 0.002 | 0.002 | Headgroup = PE -- SumDBs = 1 | 0.001 | 0.001 | LipidClass = PS -- SumLength = 40 | 0.002 | 0.002 |
| LipidClass = TG -- SumLength = 54 | 0.001 | 0.001 | SumDBs = 1 -- SumLength = 32 | 0.002 | 0.002 | LipidClass = LPE -- SumDBs = 1 | 0.001 | 0.001 | Headgroup = PS -- SumLength = 40 | 0.002 | 0.002 |
| Headgroup = -- SumLength = 54 | 0.001 | 0.001 | Headgroup = PE -- SumLength = 40 | 0.002 | 0.002 | LipidClass = PC O- -- SumLength = 38 | 0.001 | 0.001 | LipidClass = PC -- SumDBs = 2 | 0.002 | 0.002 |
| Headgroup = PE -- SumLength = 18 | 0.001 | 0.001 | Headgroup = -- SumDBs = 5 | 0.002 | 0.002 | SumDBs = 3 -- SumLength = 18 | 0.001 | 0.001 | Headgroup = PI -- SumDBs = 3 | 0.002 | 0.002 |
| LipidClass = LPE -- SumLength = 18 | 0.001 | 0.001 | SumDBs = 0 -- SumLength = 16 | 0.002 | 0.002 | Headgroup = -- SumLength = 38 | 0.001 | 0.001 | LipidClass = PI -- SumDBs = 3 | 0.002 | 0.002 |
| LipidClass = PS -- SumLength = 38 | 0.001 | 0.001 | SumDBs = 1 -- SumLength = 18 | 0.002 | 0.002 | Headgroup = PE -- SumDBs = 2 | 0.001 | 0.001 | SumDBs = 3 -- SumLength = 54 | 0.002 | 0.002 |
| Headgroup = PS -- SumLength = 38 | 0.001 | 0.001 | SumDBs = 4 -- SumLength = 52 | 0.002 | 0.002 | LipidClass = Cer -- SumDBs = 2 | 0.001 | 0.001 | LipidClass = CE -- SumLength = 18 | 0.002 | 0.002 |
| LipidClass = CE -- SumDBs = 1 | 0.001 | 0.001 | Headgroup = PS -- SumDBs = 8 | 0.002 | 0.002 | Headgroup = PC -- SumDBs = 6 | 0.001 | 0.001 | Headgroup = PC -- SumLength = 40 | 0.002 | 0.002 |
| Headgroup = CE -- SumDBs = 1 | 0.001 | 0.001 | LipidClass = PS -- SumDBs = 8 | 0.002 | 0.002 | LipidClass = CE -- SumDBs = 0 | 0.001 | 0.001 | Headgroup = CE -- SumLength = 18 | 0.002 | 0.002 |
| SumDBs = 3 -- SumLength = 54 | 0.001 | 0.001 | LipidClass = Cer -- SumLength = 38 | 0.002 | 0.002 | Headgroup = CE -- SumDBs = 0 | 0.001 | 0.001 | LipidClass = TG -- SumDBs = 3 | 0.002 | 0.002 |
| LipidClass = SM -- SumDBs = 2 | 0.001 | 0.001 | Headgroup = PE -- SumLength = 20 | 0.002 | 0.002 | LipidClass = PC O- -- SumDBs = 2 | 0.001 | 0.001 | SumDBs = 2 -- SumLength = 34 | 0.002 | 0.002 |
| Headgroup = PS -- SumDBs = 6 | 0.001 | 0.001 | LipidClass = LPE -- SumDBs = 4 | 0.002 | 0.002 | LipidClass = LPC -- SumLength = 20 | 0.001 | 0.001 | LipidClass = PE O- -- SumDBs = 5 | 0.002 | 0.002 |
| LipidClass = PS -- SumDBs = 6 | 0.001 | 0.001 | LipidClass = LPE -- SumLength = 20 | 0.002 | 0.002 | LipidClass = LPC -- SumDBs = 4 | 0.001 | 0.001 | SumDBs = 2 -- SumLength = 36 | 0.002 | 0.002 |
| LipidClass = PG -- SumDBs = 2 | 0.001 | 0.001 | SumDBs = 6 -- SumLength = 22 | 0.002 | 0.002 | Headgroup = PC -- SumLength = 20 | 0.001 | 0.001 | Headgroup = PC -- SumLength = 36 | 0.002 | 0.002 |
| LipidClass = PG -- SumLength = 38 | 0.001 | 0.001 | SumDBs = 1 -- SumLength = 36 | 0.002 | 0.002 | LipidClass = PI -- SumDBs = 5 | 0.001 | 0.001 | LipidClass = LPC -- SumDBs = 2 | 0.002 | 0.002 |
| Headgroup = PG -- SumLength = 38 | 0.001 | 0.001 | Headgroup = PC -- SumLength = 22 | 0.002 | 0.002 | Headgroup = PI -- SumDBs = 5 | 0.001 | 0.001 | SumDBs = 2 -- SumLength = 18 | 0.002 | 0.002 |
| LipidClass = PC O- -- SumLength = 40 | 0.001 | 0.001 | LipidClass = LPC -- SumLength = 22 | 0.002 | 0.002 | LipidClass = PS -- SumLength = 36 | 0.001 | 0.001 | LipidClass = PC -- SumDBs = 6 | 0.002 | 0.002 |
| LipidClass = PC -- SumDBs = 1 | 0.001 | 0.001 | LipidClass = LPC -- SumDBs = 6 | 0.002 | 0.002 | Headgroup = PS -- SumLength = 36 | 0.001 | 0.001 | Headgroup = PC -- SumLength = 38 | 0.002 | 0.002 |
| LipidClass = PE -- SumLength = 38 | 0.001 | 0.001 | SumDBs = 8 -- SumLength = 40 | 0.003 | 0.003 | LipidClass = LPG -- SumDBs = 3 | 0.002 | 0.002 | Headgroup = PE -- SumDBs = 5 | 0.002 | 0.002 |
| Headgroup = PI -- SumLength = 34 | 0.002 | 0.002 | Headgroup = PG -- SumDBs = 2 | 0.003 | 0.003 | SumDBs = 3 -- SumLength = 34 | 0.002 | 0.002 | LipidClass = PC -- SumDBs = 3 | 0.002 | 0.002 |
| LipidClass = PI -- SumLength = 34 | 0.002 | 0.002 | Headgroup = PC -- SumDBs = 5 | 0.003 | 0.003 | SumDBs = 1 -- SumLength = 16 | 0.002 | 0.002 | LipidClass = LPC -- SumDBs = 0 | 0.002 | 0.002 |
| SumDBs = 6 -- SumLength = 40 | 0.002 | 0.002 | Headgroup = PG -- SumLength = 34 | 0.003 | 0.003 | Headgroup = P -- SumDBs = 2 | 0.002 | 0.002 | LipidClass = LPI -- SumLength = 20 | 0.002 | 0.002 |
| Headgroup = PC -- SumDBs = 1 | 0.002 | 0.002 | LipidClass = PG -- SumLength = 34 | 0.003 | 0.003 | LipidClass = PA -- SumDBs = 2 | 0.002 | 0.002 | LipidClass = LPI -- SumDBs = 4 | 0.002 | 0.002 |
| LipidClass = PE O- -- SumDBs = 5 | 0.002 | 0.002 | LipidClass = PC -- SumLength = 34 | 0.003 | 0.003 | SumDBs = 2 -- SumLength = 50 | 0.002 | 0.002 | Headgroup = PI -- SumLength = 20 | 0.002 | 0.002 |
| LipidClass = LPE -- SumDBs = 1 | 0.002 | 0.002 | Headgroup = P -- SumDBs = 1 | 0.003 | 0.003 | LipidClass = Cer -- SumLength = 38 | 0.002 | 0.002 | LipidClass = PE O- -- SumLength = 38 | 0.002 | 0.002 |
| LipidClass = PC -- SumLength = 38 | 0.002 | 0.002 | LipidClass = PA -- SumDBs = 1 | 0.003 | 0.003 | Headgroup = PI -- SumDBs = 4 | 0.002 | 0.002 | LipidClass = Cer -- SumDBs = 1 | 0.002 | 0.002 |
| LipidClass = LPI -- SumLength = 20 | 0.002 | 0.002 | LipidClass = PA -- SumLength = 36 | 0.003 | 0.003 | SumDBs = 1 -- SumLength = 32 | 0.002 | 0.002 | SumDBs = 3 -- SumLength = 36 | 0.002 | 0.002 |
| Headgroup = PI -- SumLength = 20 | 0.002 | 0.002 | Headgroup = P -- SumLength = 36 | 0.003 | 0.003 | SumDBs = 1 -- SumLength = 42 | 0.002 | 0.002 | Headgroup = P -- SumLength = 34 | 0.002 | 0.002 |
| LipidClass = LPI -- SumDBs = 4 | 0.002 | 0.002 | SumDBs = 3 -- SumLength = 34 | 0.003 | 0.003 | LipidClass = TG -- SumLength = 50 | 0.002 | 0.002 | LipidClass = PA -- SumLength = 34 | 0.002 | 0.002 |
| LipidClass = TG -- SumDBs = 3 | 0.002 | 0.002 | LipidClass = Cer -- SumDBs = 2 | 0.003 | 0.003 | Headgroup = -- SumLength = 50 | 0.002 | 0.002 | LipidClass = TG -- SumDBs = 2 | 0.003 | 0.003 |
| Headgroup = PE -- SumDBs = 5 | 0.002 | 0.002 | Headgroup = -- SumDBs = 4 | 0.003 | 0.003 | LipidClass = SM -- SumLength = 41 | 0.002 | 0.002 | LipidClass = PC -- SumLength = 38 | 0.003 | 0.003 |
| Headgroup = PC -- SumLength = 16 | 0.002 | 0.002 | Headgroup = PC -- SumDBs = 3 | 0.003 | 0.003 | Headgroup = PC -- SumLength = 41 | 0.002 | 0.002 | LipidClass = LPG -- SumDBs = 1 | 0.003 | 0.003 |
| LipidClass = LPC -- SumLength = 16 | 0.002 | 0.002 | LipidClass = PG -- SumDBs = 2 | 0.003 | 0.003 | LipidClass = PE -- SumLength = 34 | 0.002 | 0.002 | LipidClass = PC -- SumLength = 36 | 0.003 | 0.003 |
| Headgroup = PE -- SumDBs = 1 | 0.002 | 0.002 | LipidClass = LPI -- SumLength = 16 | 0.003 | 0.003 | Headgroup = PS -- SumDBs = 7 | 0.002 | 0.002 | LipidClass = LPC -- SumLength = 16 | 0.003 | 0.003 |
| Headgroup = PC -- SumDBs = 6 | 0.002 | 0.002 | Headgroup = PI -- SumLength = 16 | 0.003 | 0.003 | LipidClass = PS -- SumDBs = 7 | 0.002 | 0.002 | Headgroup = PC -- SumLength = 16 | 0.003 | 0.003 |
| LipidClass = Cer -- SumDBs = 1 | 0.002 | 0.002 | LipidClass = PC O- -- SumDBs = 0 | 0.003 | 0.003 | LipidClass = SM -- SumDBs = 2 | 0.002 | 0.002 | Headgroup = PC -- SumDBs = 2 | 0.003 | 0.003 |
| Headgroup = -- SumDBs = 2 | 0.002 | 0.002 | LipidClass = PC O- -- SumLength = 32 | 0.003 | 0.003 | SumDBs = 1 -- SumLength = 18 | 0.002 | 0.002 | LipidClass = SM -- SumDBs = 1 | 0.003 | 0.003 |
| LipidClass = LPC -- SumDBs = 0 | 0.002 | 0.002 | LipidClass = SM -- SumLength = 34 | 0.003 | 0.003 | LipidClass = DG -- SumDBs = 3 | 0.002 | 0.002 | Headgroup = PG -- SumDBs = 5 | 0.003 | 0.003 |
| LipidClass = PG -- SumDBs = 5 | 0.002 | 0.002 | SumDBs = 6 -- SumLength = 38 | 0.004 | 0.004 | SumDBs = 0 -- SumLength = 16 | 0.002 | 0.002 | LipidClass = PG -- SumDBs = 5 | 0.003 | 0.003 |
| Headgroup = PG -- SumDBs = 5 | 0.002 | 0.002 | LipidClass = PC O- -- SumDBs = 7 | 0.004 | 0.004 | Headgroup = -- SumLength = 34 | 0.002 | 0.002 | LipidClass = PE -- SumLength = 38 | 0.003 | 0.003 |
| SumDBs = 2 -- SumLength = 34 | 0.002 | 0.002 | LipidClass = TG -- SumDBs = 4 | 0.004 | 0.004 | SumDBs = 4 -- SumLength = 52 | 0.002 | 0.002 | Headgroup = -- SumDBs = 1 | 0.003 | 0.003 |
| LipidClass = PA -- SumLength = 34 | 0.002 | 0.002 | LipidClass = PC O- -- SumDBs = 3 | 0.004 | 0.004 | LipidClass = PA -- SumLength = 36 | 0.002 | 0.002 | SumDBs = 0 -- SumLength = 18 | 0.003 | 0.003 |
| Headgroup = P -- SumLength = 34 | 0.002 | 0.002 | LipidClass = PE -- SumDBs = 6 | 0.004 | 0.004 | Headgroup = P -- SumLength = 36 | 0.002 | 0.002 | Headgroup = PC -- SumDBs = 1 | 0.003 | 0.003 |
| Headgroup = PE -- SumDBs = 4 | 0.002 | 0.002 | Headgroup = PI -- SumLength = 38 | 0.004 | 0.004 | LipidClass = PC O- -- SumDBs = 7 | 0.002 | 0.002 | SumDBs = 2 -- SumLength = 52 | 0.004 | 0.004 |
| Headgroup = PC -- SumLength = 40 | 0.002 | 0.002 | LipidClass = PI -- SumLength = 38 | 0.004 | 0.004 | Headgroup = PI -- LipidClass = LPI | 0.002 | 0.002 | Headgroup = PI -- SumLength = 34 | 0.004 | 0.004 |
| LipidClass = PE O- -- SumLength = 38 | 0.002 | 0.002 | Headgroup = PS -- SumDBs = 4 | 0.004 | 0.004 | LipidClass = Cer -- SumLength = 34 | 0.002 | 0.002 | LipidClass = PI -- SumLength = 34 | 0.004 | 0.004 |
| SumDBs = 4 -- SumLength = 38 | 0.002 | 0.002 | LipidClass = PS -- SumDBs = 4 | 0.004 | 0.004 | LipidClass = LPI -- SumLength = 16 | 0.002 | 0.002 | Headgroup = PI -- SumLength = 36 | 0.004 | 0.004 |
| LipidClass = PC O- -- SumDBs = 6 | 0.002 | 0.002 | Headgroup = PI -- SumDBs = 6 | 0.004 | 0.004 | Headgroup = PI -- SumLength = 16 | 0.002 | 0.002 | LipidClass = PI -- SumLength = 36 | 0.004 | 0.004 |
| LipidClass = TG -- SumDBs = 2 | 0.002 | 0.002 | LipidClass = PI -- SumDBs = 6 | 0.004 | 0.004 | LipidClass = TG -- SumDBs = 4 | 0.002 | 0.002 | SumDBs = 5 -- SumLength = 38 | 0.004 | 0.004 |
| SumDBs = 2 -- SumLength = 42 | 0.003 | 0.003 | Headgroup = PC -- SumDBs = 7 | 0.004 | 0.004 | SumDBs = 5 -- SumLength = 36 | 0.002 | 0.002 | Headgroup = P -- SumDBs = 4 | 0.004 | 0.004 |
| SumDBs = 2 -- SumLength = 52 | 0.003 | 0.003 | Headgroup = PI -- LipidClass = LPI | 0.004 | 0.004 | Headgroup = CE -- SumLength = 16 | 0.002 | 0.002 | LipidClass = PA -- SumDBs = 4 | 0.004 | 0.004 |
| LipidClass = PE -- SumDBs = 4 | 0.003 | 0.003 | Headgroup = PI -- SumLength = 18 | 0.004 | 0.004 | Headgroup = PE -- SumLength = 34 | 0.002 | 0.002 | SumDBs = 1 -- SumLength = 41 | 0.004 | 0.004 |
| LipidClass = Cer -- SumLength = 40 | 0.003 | 0.003 | LipidClass = LPI -- SumLength = 18 | 0.004 | 0.004 | LipidClass = CE -- SumLength = 16 | 0.002 | 0.002 | LipidClass = PS -- SumDBs = 6 | 0.004 | 0.004 |
| Headgroup = -- SumLength = 40 | 0.003 | 0.003 | LipidClass = PI -- SumDBs = 5 | 0.004 | 0.004 | LipidClass = PI -- SumDBs = 4 | 0.002 | 0.002 | Headgroup = PS -- SumDBs = 6 | 0.004 | 0.004 |
| Headgroup = -- SumDBs = 1 | 0.003 | 0.003 | Headgroup = PI -- SumDBs = 5 | 0.004 | 0.004 | Headgroup = PC -- SumDBs = 7 | 0.002 | 0.002 | Headgroup = PG -- SumLength = 38 | 0.005 | 0.005 |
| SumDBs = 2 -- SumLength = 36 | 0.003 | 0.003 | Headgroup = PC -- SumLength = 32 | 0.004 | 0.004 | LipidClass = PC -- SumLength = 34 | 0.003 | 0.003 | LipidClass = PG -- SumLength = 38 | 0.005 | 0.005 |
| LipidClass = PC O- -- SumDBs = 4 | 0.003 | 0.003 | SumDBs = 7 -- SumLength = 40 | 0.004 | 0.004 | LipidClass = PG -- SumDBs = 1 | 0.003 | 0.003 | LipidClass = PC O- -- SumDBs = 4 | 0.005 | 0.005 |
| SumDBs = 1 -- SumLength = 40 | 0.003 | 0.003 | LipidClass = PI -- SumLength = 40 | 0.004 | 0.004 | SumDBs = 7 -- SumLength = 38 | 0.003 | 0.003 | SumDBs = 6 -- SumLength = 40 | 0.005 | 0.005 |
| LipidClass = PA -- SumDBs = 4 | 0.003 | 0.003 | Headgroup = PI -- SumLength = 40 | 0.004 | 0.004 | LipidClass = PE O- -- SumDBs = 7 | 0.003 | 0.003 | LipidClass = PE -- SumDBs = 4 | 0.005 | 0.005 |
| Headgroup = P -- SumDBs = 4 | 0.003 | 0.003 | SumDBs = 0 -- SumLength = 32 | 0.005 | 0.005 | Headgroup = -- SumDBs = 4 | 0.004 | 0.004 | Headgroup = PE -- SumLength = 38 | 0.006 | 0.006 |
| SumDBs = 5 -- SumLength = 38 | 0.003 | 0.003 | LipidClass = PC -- SumDBs = 4 | 0.005 | 0.005 | LipidClass = LPI -- SumDBs = 1 | 0.004 | 0.004 | Headgroup = PE -- SumDBs = 4 | 0.006 | 0.006 |
| SumDBs = 1 -- SumLength = 41 | 0.003 | 0.003 | Headgroup = PC -- SumLength = 34 | 0.005 | 0.005 | Headgroup = PI -- SumDBs = 1 | 0.004 | 0.004 | Headgroup = PI -- SumDBs = 2 | 0.007 | 0.007 |
| Headgroup = PE -- SumLength = 38 | 0.004 | 0.004 | SumDBs = 2 -- SumLength = 41 | 0.005 | 0.005 | Headgroup = PE -- SumDBs = 7 | 0.004 | 0.004 | LipidClass = PI -- SumDBs = 2 | 0.007 | 0.007 |
| Headgroup = PI -- SumLength = 36 | 0.004 | 0.004 | SumDBs = 5 -- SumLength = 40 | 0.005 | 0.005 | LipidClass = PI -- SumLength = 38 | 0.004 | 0.004 | LipidClass = Cer -- SumLength = 40 | 0.007 | 0.007 |
| LipidClass = PI -- SumLength = 36 | 0.004 | 0.004 | Headgroup = PE -- SumDBs = 6 | 0.005 | 0.005 | Headgroup = PI -- SumLength = 38 | 0.004 | 0.004 | Headgroup = -- SumLength = 40 | 0.007 | 0.007 |
| Headgroup = PI -- SumDBs = 2 | 0.006 | 0.006 | LipidClass = LPI -- SumDBs = 1 | 0.008 | 0.008 | SumDBs = 2 -- SumLength = 41 | 0.004 | 0.004 | SumDBs = 1 -- SumLength = 40 | 0.008 | 0.008 |
| LipidClass = PI -- SumDBs = 2 | 0.006 | 0.006 | Headgroup = PI -- SumDBs = 1 | 0.008 | 0.008 | SumDBs = 7 -- SumLength = 40 | 0.004 | 0.004 | SumDBs = 4 -- SumLength = 38 | 0.009 | 0.009 |

  


##### Best Substructure Features

KO CHOW

KO HFCD

WT CHOW
